# Ensemblify: a user-friendly tool for generating ensembles of intrinsically disordered regions of AlphaFold and user-defined models

**DOI:** 10.1101/2025.08.26.672300

**Authors:** Nuno P. Fernandes, Tiago Gomes, Tiago N. Cordeiro

## Abstract

**Motivation:** Intrinsically disordered proteins (IDPs) and regions (IDRs) challenge structural characterization due to their dynamic conformational ensembles and lack of stable structure. Existing computational approaches for modelling these ensembles are either computationally intensive, limited in flexibility, or inaccessible to non-experts, especially when dealing with multi-domain or multi-chain proteins.

**Results:** We present Ensemblify, an open-source, user-friendly Python package for generating and analyzing conformational ensembles of IDPs/IDRs. Ensemblify uses a Monte Carlo algorithm coupled with neighbour-aware sampling of dihedral angles from curated or user-defined fragment libraries to explore conformational space. It directly incorporates information from AlphaFold’s confidence metrics as flexible energy restraints in PyRosetta to guide the sampling. It supports multi-chain and multi-domain proteins and can sample N-terminal, C-terminal, and inter-domain linkers while preserving folded regions. Ensemble quality can be validated and refined against experimental data such as SAXS via Bayesian/Maximum Entropy (BME) reweighting. Interactive dashboards provide in-depth structural analysis and comparison. Testing across 10 diverse proteins demonstrated Ensemblify’s accuracy, flexibility, and ability to recover experimentally observed structural features. Incorporating AlphaFold confidence metrics shows potential to improve the ensemble-data agreement.

**Availability:** Ensemblify is freely available at https://github.com/CordeiroLab/ensemblify, along with detailed installation instructions and usage tutorials. Ensemblify can be used for scripting through its Python API or directly through the provided command-line interface (CLI). Complete documentation is available within the source-code and CLI and on Ensemblify’s official documentation page (https://ensemblify.readthedocs.io).

**Supplementary information:** Supplementary data is available online.

## Introduction

Intrinsically disordered proteins (IDPs) are a unique class of proteins that lack a stable, well-defined three-dimensional structure under physiological conditions (Holehouse and Kragelund 2024). Unlike their folded counterparts, IDPs exist as highly dynamic conformational ensembles (Tesei *et al*. 2024) but often exhibit nonrandom structural heterogeneity that enables specific interactions with various biological partners (Tompa *et al*. 2015, Csizmok *et al*. 2016).

The emergence of AlphaFold has enabled accurate prediction of folded protein structures at the proteome scale (Jumper *et al*. 2021, Varadi *et al*. 2024). However, reliably modelling IDPs remains a major challenge (Ruff and Pappu 2021, Lane 2023). Notably, AlphaFold’s predicted Local Distance Difference Test (pLDDT) metric has emerged as a reliable indicator of disorder, with low pLDDT values correlating with unstructured regions (Alderson *et al*. 2023). Despite this, AlphaFold’s potential to identify and characterize disordered regions has been largely underutilized in probing the conformational landscapes of IDPs and intrinsically disordered regions (IDRs). Consequently, the conformational properties of IDPs/IDRs remain poorly understood and challenging to predict, also owing to their low sequence conservation and limited experimental characterization. Thus, ensemble generation remains essential for capturing their dynamic nature and for modelling their interactions with binding partners (Moses *et al*. 2023).

Exploring the conformational landscape of IDPs/IDRs is computationally intensive and often requires expert knowledge. The current state of the art has evolved into three main approaches: (i) structural database sampling (SDS), which sample dihedral angles from probability distributions derived from experimental data (Feldman and Hogue 2000, Ozenne *et al*. 2012, Estaña *et al*. 2019, Harmat, Dudola and Gáspári 2021, Teixeira *et al*. 2022, Liu *et al*. 2023, Pajkos *et al*. 2025); (ii) molecular dynamics (MD) simulations using physical force fields (Pietrek, Stelzl and Hummer 2020, Shrestha, Smith and Petridis 2021, Thomasen *et al*. 2024); and (iii) artificial intelligence/machine learning (AI/ML) methods (Del Alamo *et al*. 2022, Janson *et al*. 2023, Aupič *et al*. 2024, Wayment-Steele *et al*. 2024, Kalakoti and Wallner 2024, Wu *et al*. 2024). All-atom MD and advanced AI/ML methods, while offering high resolution and predictive power, are computationally expensive and typically require domain expertise. In contrast, coarse-grained (CG)-MD is more computationally efficient and allows access to broader conformational sampling of IDPs (Cao *et al*. 2024, Wang *et al*. 2025). However, this comes at the cost of reduced atomic resolution and underestimation of local structural detail, which can be critical for modelling IDPs accurately. SDS methods often prioritize accessibility through web servers (Ziegler *et al*. 2016, Pajkos *et al*. 2025) at the expense of versatility and user customization, or they offer flexibility with complex setups and limited documentation, making them inaccessible to less experienced researchers (Ozenne *et al*. 2012, Harmat, Dudola and Gáspári 2021, Teixeira *et al*. 2022, Liu *et al*. 2023). Moreover, current methods are usually not applicable to multi-domain or multi-chain proteins, whose structural dynamics remain challenging to model due to the IDRs that make up inter-domain linkers or terminal tails.

AlphaFold’s confidence metrics, such as the pLDDT and the predicted aligned error (PAE) (Guo *et al*. 2022, Brotzakis *et al*. 2025), convey structural information that could help overcome these limitations, but they are rarely exploited in ensemble generation. Likewise, the nonrandom secondary structure propensities in IDPs/IDRs are not always captured by available methods.

To address these challenges, we introduce Ensemblify, a Python package for generating protein conformational ensembles. Ensemblify is fast, versatile, open-source, and easily extensible, making it suitable to users with differing levels of expertise. It supports diverse protein structural architectures, directly incorporates information from AlphaFold’s confidence metrics, and realistically samples fractional secondary structure propensities in disordered regions. Additionally, it features robust documentation, a user-friendly HTML parameters form, and a command-line interface (CLI) that offers easy access to its key features. By combining ensemble generation, analysis, experimental validation, and refinement through user-friendly dashboards, Ensemblify provides a powerful, accessible resource for both research and teaching.

## Description and functionality

### Workflow & Implementation

To generate a conformational ensemble, Ensemblify requires a parameters file with mandatory input fields, including the input structure or sequence and a dihedral angle database (Fig. 1). This file can be easily created using the provided user-friendly HTML interface (Supplementary Fig. S6). The input structure can be a predicted model from AlphaFold, optionally including its PAE matrix, a user-defined model, or an amino acid sequence in the case of full IDPs (Fig. 1A). Users may also combine experimentally determined structures of folded protein domains with sequences of disordered linker or terminal regions to create a full-length starting structure. To do this, the user must provide the sequences of all protein regions (folded and disordered) in FASTA format, and the structures of folded domains as PDB files, in order from N- to C-terminal. Alternatively, a UniProt accession number can be provided in place of a structure and/or PAE matrix to automatically retrieve the corresponding file from the AlphaFold Protein Structure Database (Varadi *et al*. 2024).

**Figure 1.**
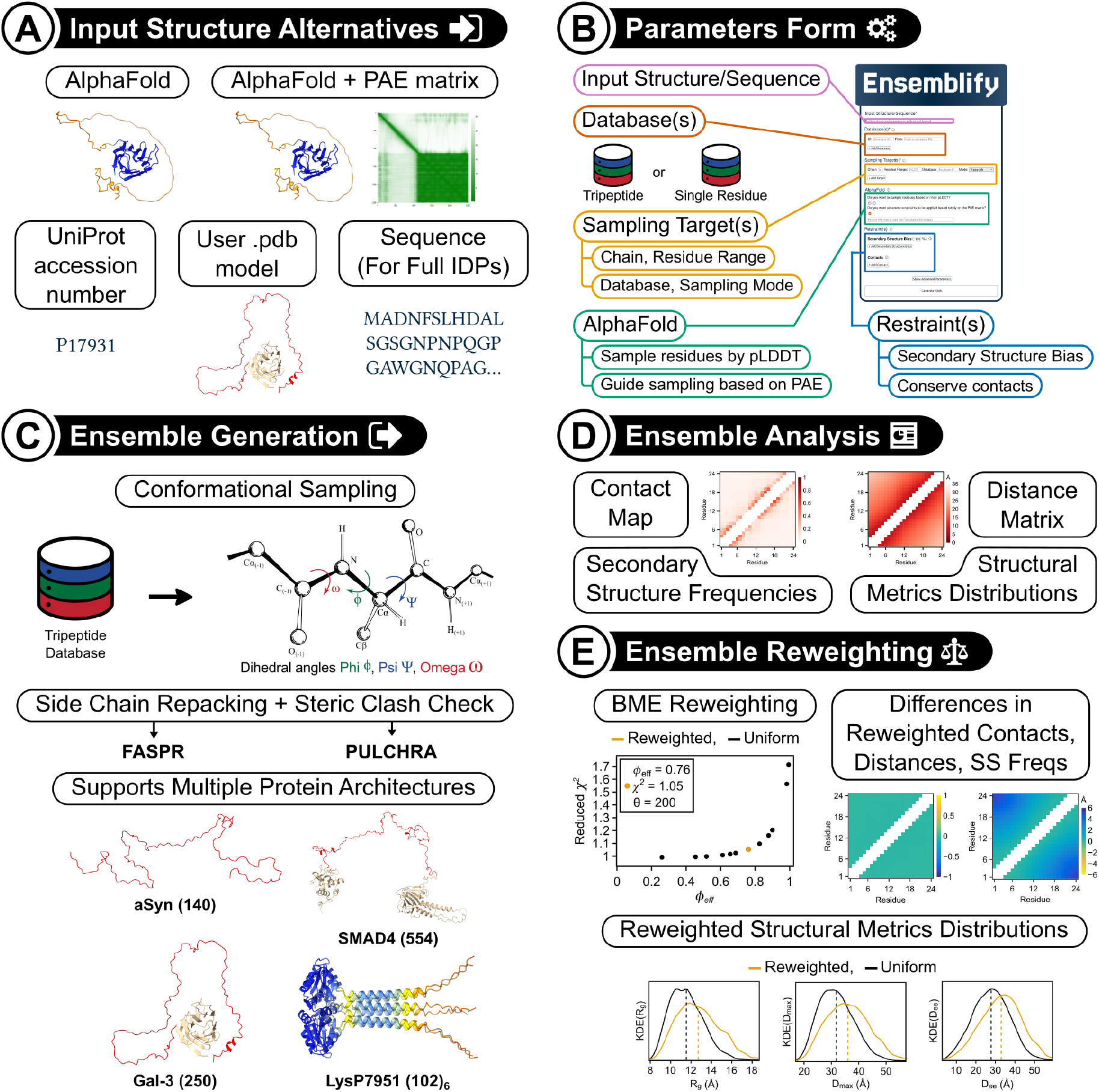
General workflow of the Ensemblify software tool. **A**. Input structure can be provided in various forms: an AF-2 model, with or without its PAE matrix, a UniProt accession number, a user-defined model, or a protein sequence. **B**. All mandatory inputs like the input structure, dihedral angle database and target sampling regions are defined through a user-friendly HTML form, which outputs a properly formatted YAML parameters file for ensemble generation. **C**. Based on the YAML file, conformational ensembles are generated by sampling dihedral angles from the user-provided database and inserting them into the protein backbone using a Monte Carlo algorithm. This is followed by side chain repacking using FASPR and a steric clash check using PULCHRA. Ensembles can sample flexible N- and/or C-terminal tails and/or inter-domain linkers across diverse protein architectures. **D**. Interactive HTML dashboards enable exploration of ensemble-averaged structural properties and intuitive data visualization. **E**. Experimental data (*e*.*g*. SAXS) can be used to reweight the ensemble via BME method. Structural metrics of reweighted ensembles are compared to their uniformly weighted counterparts through a dedicated reweighting dashboard.

Target sampling regions can be manually appointed for each protein chain in the input structure using the parameters file. Any regions not selected are restrained to their initial configuration (Fig. 1B). Each designated sampling region must be assigned a database and a sampling mode: tripeptide or single residue (Estaña *et al*. 2019, González-Delgado *et al*. 2022), which respectively dictate if neighbouring residues are considered or ignored during sampling. By default, all residues within the specified ranges are sampled. When using an AlphaFold model, users can also choose to automatically restrict sampling to residues within these regions with pLDDT below a certain threshold (default is 70).

Ensemble generation in Ensembify relies on PyRosetta, the Python interface for the widely used Rosetta molecular modelling suite (Chaudhury, Lyskov and Gray 2010). Prior to sampling, the input structure is processed by converting side-chains to a ‘centroid’ representation to increase sampling efficiency (Supplementary Information S1.1). Ensembles are generated through protein backbone conformational sampling (Fig. 1C, Supplementary Information S1.2). This process is guided by energy restraints that preserve the structure of folded domains and inter-chain contacts, with optional additional restraints to bias sampling towards desired secondary structure properties (Supplementary Information S1.2.2). During sampling, Ensemblify performs a Monte Carlo move targeting each residue within a defined sampling region, continuing until all residues have been targeted. In each move, the protein’s backbone is perturbed by sampling dihedral angles φ, ψ and ω from a normal distribution centered around values found in a dihedral angle database. Ensemblify offers a three-residue fragment dihedral angle database but also supports user-defined databases. Multiple databases can be used in the same sampling protocol, and strategies to reduce memory usage of large databases are detailed in Supplementary Information S1.2.3. After each Monte Carlo move, the energy of the newly generated conformation is evaluated using a Rosetta score function. This includes a weak Van der Waals repulsive term to penalize steric clashes and additional terms to penalize the violation of applied structural restraints. The move is then accepted or rejected according to the standard Metropolis criterion (Metropolis *et al*. 1953) (Supplementary Information S1.2.1, Supplementary Equation S1). Once all target residues have been sampled, the resulting structure undergoes a minimization protocol before output. To restore atomic detail, side chains of the sampled regions are repacked using FASPR, an ultra-fast and accurate program for deterministic protein side chain packing (Huang, Pearce and Zhang 2020). Final structures are then processed with PULCHRA (Rotkiewicz and Skolnick 2008) to check for steric clashes using a 2 Å threshold, and any structures with clashes are discarded. The sampling cycle is repeated until the desired number of valid sampled structures is reached. Strategies used to further increase variability between sampled structures are detailed in Supplementary Information S1.2.4.

Given a protein conformational ensemble, Ensemblify can create an interactive structural analysis dashboard that supports zooming, panning and trace selection for all figures (Figure 1D, Supplementary Fig. S7). When multiple ensembles are provided, they can be easily compared through a similar analysis dashboard (Supplementary Fig. S8). Analysis metrics include a contact frequency map with a 4.5 Å threshold, a C_α_ average distance matrix, and a secondary structure assignment frequency plot following the simplified DSSP (Kabsch and Sander 1983) classifications of α-helix, β-sheet and random coil. Additionally, the dashboard provides probability density distributions for the radius of gyration (*R*_*g*_), maximum C_α_ distance (*D*_*max*_) and end-to-end distance (*D*_*ee*_), with optional center mass distance distributions between any two user-defined protein regions. Calculation of additional analysis metrics can be easily implemented by exploiting Ensemblify’s modular architecture.

When provided with experimental small angle X-ray scattering (SAXS) data, Ensemblify can reweight an existing structural ensemble through a Bayesian/Maximum Entropy (BME) approach (Bottaro, Bengtsen and Lindorff-Larsen 2018) (Supplementary Information S2). This process outputs an interactive graphical dashboard that compares the original and reweighted ensembles based on the previously mentioned analysis metrics, as well as their agreement with SAXS data (Fig. 1E, Supplementary Fig. S9). The BME method involves finding the set of optimal conformer weights (Supplementary Equation S4) that achieve good agreement with experimental data (*i*.*e*., low reduced *χ*^2^; Supplementary Equation S5) while maximizing relative entropy (*i*.*e*., high *ϕ*_eff_ ; Supplementary Equations S6-S7). Tuning the scaling parameter *θ* in Supplementary Equation S4 results in different values of *χ*^2^ and *ϕ*_eff_. Consequently, plotting *ϕ*_eff_ versus *χ*^2^ as *θ* changes leads to the optimal value of *θ*, located at the point of maximum curvature, *i*.*e*. the “elbow” of the plot (Fig. 1E, Supplementary Fig. S5). Ensemble generation in Ensemblify is parallelized using Ray (Moritz *et al*. 2017), a distributed computing library for scaling Python applications. Ensemble analysis and reweighting calculations rely on the MDAnalysis (Michaud-Agrawal *et al*. 2011, Gowers *et al*. 2016), MDtraj (McGibbon *et al*. 2015), Pandas (McKinney 2010), NumPy (Harris *et al*. 2020) and SciPy (Virtanen *et al*. 2020) Python libraries in combination with the multiprocessing Python library to speed up calculations. Interactive graphical dashboards are created using Plotly (Inc 2015), a browser-based Python graphing library. All the calculated structural analysis data used in the creation of interactive dashboards, including back-calculated SAXS profiles and their fitting to experimental SAXS data, are provided as plain text files, allowing users to apply their own analysis pipelines.

### Applications

Ensemblify has been successfully applied to proteins of varying size and structural architectures (Supplementary Fig. S10, Supplementary Information S3.2), showcasing its ability to sample N-terminal and C-terminal disordered tails, as well as inter-domain linkers, while preserving the structure of folded domains and, for multi-chain proteins, inter-chain interfaces.

**Full IDPs** tested (Supplementary Figs S10A, S11-S12, Supplementary Information 3.2.1) include: (**1**) *Histatin5 (Hst5)*, an antimicrobial peptide found naturally in human saliva (Thomasen *et al*. 2024, Jephthah *et al*. 2019), and (**2**) *α-synuclein (aSyn)*, a presynaptic neuronal protein implicated in Parkinson’s disease (Thomasen *et al*. 2024, Ahmed *et al*. 2021).

**Multi-domain proteins with flexible linkers** (Supplementary Figs S10B, S13-S17, Supplementary Information 3.2.2): (**3**) A truncated construct of *cardiac myosin-binding protein C (cMyBP-C)*, which plays important roles in the muscle sarcomere by interacting with myosin and actin, composed of the tri-helix bundle of the m-domain connected to the C2 domain by a flexible linker (cMyBP-C_mTHB-C2_) (Thomasen *et al*. 2024, Michie *et al*. 2016); (**4**) *Linear tetraubiquitin (Ubq*_*4*_) (Thomasen *et al*. 2024, Jussupow *et al*. 2020); (***5****) A C-terminal truncated construct of TIA1*, a regulator of transcription and RNA translation (Thomasen *et al*. 2024, Sonntag *et al*. 2017); and (**6**) *SMAD4*, a mediator of TGF-β signal transduction (Gomes *et al*. 2021).

**Proteins with folded domains and long disordered tails** (Supplementary Figs S10C, S18-S20, Supplementary Information 3.2.3): (**7**) a construct of the SH4, Unique and SH3 domains of *non-receptor tyrosine kinase Src (USH3)* (Arbesú *et al*. 2017), implicated in cell signaling pathways related to cell growth, migration, invasion, and survival; (**8**) *Galectin-3 (Gal-3)* (Thomasen *et al*. 2024, Lin *et al*. 2017), a lectin that binds to β-galactoside and can regulate cell signaling; and (**9**) *heterogeneous nuclear ribonucleoprotein A1 (hnRNPA1)* (Thomasen *et al*. 2024, Martin *et al*. 2021), that has an intrinsically disordered low-complexity domain where mutations can lead to amyotrophic lateral sclerosis.

**Multi-chain protein** (Supplementary Figs S10D, S21, Supplementary Information 3.2.4): (**10**) a homohexamer construct of the C-terminal product subunit of the endolysin of *S. thermophilus* phage P7951 *(LysP7951)* (Jumper *et al*. 2021, Pinto *et al*. 2022).

A comprehensive description and analysis of these case studies is provided in Supplementary Information 3.2.

Theoretical SAXS profiles calculated from the ensembles of most of the tested proteins show reasonable agreement with experimental SAXS data, and, in all cases, this fitting improves after BME ensemble reweighting. The time required to generate ensembles of 10,000 structures per protein, along with references for the protein structures and corresponding experimental SAXS data, are provided in Supplementary Tables S3-S4.

Ensemblify offers a way to convert the structural information from AlphaFold’s PAE matrix (Guo *et al*. 2022, Brotzakis *et al*. 2025) into energy restraints that guide conformational sampling (Supplementary Information S1.2.5). We evaluated this approach by generating a Gal-3 ensemble from an AF-2 prediction, using both pLDDT and PAE metrics to inform the sampling process. This ensemble was compared to one generated from a user-defined model (Supplementary Information 3.3). The latter, composed predominantly of extended conformations (average *R*_*g*_ of ∼33 Å), initially showed poor agreement with experimental SAXS data (*χ*^2^ = 20.16), though this improved substantially following BME reweighting (*χ*^2^ _*reweighted*_ = 1.50). Incorporating AF-2-derived information into the sampling led to a more compact ensemble (average *Rg* of ∼24 Å) with better SAXS agreement (*χ*^2^ = 11.95). However, after BME reweighting, the fit (*χ*^2^_*reweighted*_ = 2.45) remained inferior to that of the ensemble generated without AF input — likely due to the presence of overly compact conformations. This motivated the generation of a third ensemble in which selected inter-residue energy restraints derived from the PAE matrix were attenuated by a factor of 30, as described in Supplementary Information S1.2.5. This adjustment yielded an ensemble of moderately compact conformations (average *Rg* of ∼26 Å), leading to improved SAXS agreement both before (*χ*^2^ = 4.96) and after BME reweighting (*χ*^2^ _*reweighted*_ = 1.32), surpassing the performance of the ensemble generated without AF-2 information. These results underscore the importance of tuning the scaling of PAE-derived restraints (parameter γ in Supplementary Equation S3) and highlight the potential for users to optimize this parameter to achieve better agreement with experimental data.

## Conclusions

We present Ensemblify, a user-friendly computational framework that enables the generation of conformational ensembles for IDPs/IDRs using neighbour-aware structural database sampling. It offers ways to combine the experimentally determined structured domains with disordered segments into unified hybrid globular/disordered protein structures, which can serve as sampling starting points and enable a holistic exploration of a protein’s conformational space. Ensembify seamlessly integrates ensemble analysis with experimental validation through interactive dashboards. It is also well-documented, intuitive, and modular, allowing for easy addition of new features.

Testing Ensemblify on a diverse benchmark set of ten proteins demonstrated that the generated ensembles closely match experimental SAXS data and accurately capture experimentally determined secondary structure propensities in disordered regions. In all cases, BME reweighting further improved agreement with experiments.

Unlike existing approaches, Ensemblify uniquely integrates AlphaFold-derived confidence metrics to inform conformational sampling, by using pLDDT scores to define flexible regions and converting PAE matrix values into tunable energy restraints. This strategy led to ensembles that better fit experimental SAXS data, highlighting the potential of leveraging AlphaFold towards improved ensemble accuracy.

By bridging predictive modelling with experimental validation in an accessible and extensible platform, Ensemblify holds promise as a valuable resource for both research and education in structural biology.

## Acknowledgements

We thank Manuel N. Melo (ITQB NOVA) for his insightful help, suggestions, and access to the *in-house* computer cluster. We thank the members of the Dynamic Structural Lab (ITQB NOVA) for testing this tool.

## Funding information

TNC is the recipient of the CEECIND/01443/2017 grant. National funds funded this work through FCT: Project MOSTMICRO-ITQB (UIDB/04612/2020, UIDP/04612/2020), FEDER Funds through COMPETE 2020 (0145-FEDER-007660), LS4FUTURE (LA/P/0087/2020), and a SR&TD project (PTDC/BIA-BFS/0391/2021).

## Notes

https://github.com/CordeiroLab/ensemblify

https://ensemblify.readthedocs.io

### Competing Interest Statement

The authors have declared no competing interest.

