## Supplementary Information for "Ensemblify: a user-friendly tool for generating ensembles of intrinsically disordered regions of AlphaFold and user-defined models"

#### Summary

Ensemblify is a user-friendly tool designed to generate conformational ensembles of intrinsically disordered regions (IDRs) within AlphaFold and user-defined protein models. It employs a Monte Carlo algorithm to explore protein backbone conformations by inserting new dihedral angles sampled from a tripeptide database derived from experimental structures.
The ensemble generation process involves converting all-atom models to a centroid representation for efficiency and manipulating the FoldTree morphology to prevent "lever arm effects." Structural restraints are applied to preserve functional elements (*e.g.*, interaction and structured domains), and Monte Carlo moves are accepted or rejected based on the Metropolis Criterion. Ensemblify can utilize one or more dihedral angle databases, taking care to minimize memory usage. Sampling variability is enhanced by randomizing residue targeting order and drawing new angles from a Gaussian distribution. Notably, AlphaFold confidence metrics (PAE and pLDDT) can be integrated to guide sampling, allowing for more freedom in less confident regions.
Beyond ensemble generation, Ensemblify allows for the calculation of averaged ensemble properties like Contact Maps, Distance Matrices, Secondary Structure Assignment Frequencies and probability distributions of commonly explored structural metrics, like the Radius of Gyration ($R_g$), the Maximum Distance ($D_{max}$) and the end-to-end distance ($D_{ee}$). Ensemblify also makes use of Bayesian/Maximum Entropy (BME) ensemble reweighting to align calculated averages with experimental data while minimizing alterations to the original ensemble. This is assessed by balancing the reduced chi-square ($\chi^2$) metric with the fraction of effective frames ($\phi_{eff}$), which quantifies deviation from weight uniformity.
Ensemblify provides interactive HTML forms and analysis dashboards with various structural visualizations. Ensemblify is broadly applicable to diverse protein architectures, including fully IDPs, multi-domain proteins with flexible linkers, and multi-chain systems.

### Supplementary Methods

#### 1. Ensemble Generation

Ensemble generation is performed using a Monte Carlo algorithm that iteratively perturbs backbone dihedral angles to efficiently explore the protein's conformational energy landscape in search of new conformations.

##### 1.1. Preparation of Sampling Starting Structure

###### 1.1.1. Conversion of Side-Chains to Centroid Representation

Performing protein simulations using all-atom models is computationally expensive, since the cost of calculating all pairwise atomic interactions scales quadratically with the number of atoms in a molecular system. Here, our goal is not to model detailed backbone-side-chain interactions (like in protein-docking protocols) but rather to quickly and efficiently sample a protein's possible structural configurations using a Monte Carlo algorithm. However, the conformational energy landscape of all-atom models is very rugged, which can lead to a high rejection rate of Monte Carlo moves and trapping in local minima. To overcome this, prior to conformational sampling, the input structure is converted to what Rosetta calls a “centroid” representation: the protein backbone remains fully atomic, while each residue's side-chain is represented as a single "pseudo-atom" of varying size and properties (polarity, charge, etc.), determined by the residue's identity (Rohl *et al.* 2004). Using this representation helps reduce the complexity of the system, smoothing the conformational energy landscape, which facilitates more efficient conformational sampling.

###### 1.1.2. Manipulating the FoldTree Morphology

The modelling of protein conformations in Rosetta is based on the principle that each atom in a molecular system can be uniquely defined through its internal coordinates, *i.e.* its bond length, bond angle and torsion angle relative to its neighbouring atoms. These internal coordinates can be translated into *(x, y, z)* Cartesian coordinates for all atoms of the modelled molecular system. A physics-based energy function can then be applied to the set of Cartesian coordinates defining the current state of the system to evaluate its energy score (Leaver-Fay *et al.* 2011).

In practice, when modelling protein conformational changes, the bond length and bond angle within and between protein residues do not change significantly, and can be considered fixed. This leaves only the torsion angle values as a relevant internal coordinate, reducing the number of degrees of freedom (DOF) associated with each atom from 3 to 1, which greatly improves the speed of energy score calculations (Leaver-Fay *et al.* 2011).

In Rosetta, the AtomTree is a data structure responsible for the translation of internal coordinates to Cartesian coordinates for each atom in the protein structure. It is a directed, acyclic, connected graph: there is one atom that acts as the root of the tree, and all other atoms have exactly one parent and zero or more children. An atom's position in the chain is specified in terms of their internal coordinates, relative to their parent atom's position. The FoldTree is a coarse-grained representation of the AtomTree that only specifies connections between residues and dictates how changes in torsion angles of protein residues propagate through the rest of the structure. The direction of the graph is set by choosing a root vertex

(*e.g.* the N-terminus of the protein) so that any movement in the upstream residues (residues that are close to N-terminus) will cause movements in residues downstream (closer to C-terminus) (Leaver-Fay *et al.* 2011).
Manipulating the FoldTree can help avoid unwanted “lever arm effects”, *i.e.* changes in the conformation of downstream regions triggered by small changes in torsion angles of upstream residues. Since in our Monte Carlo sampling protocol every move involves changes in residue torsion angles, we want to minimize moves that will be ultimately rejected by leading to the unfolding of conserved domains due to “lever arm effects”. To do this, we change the FoldTree aiming towards a morphology similar to the one detailed in (Wang, Bradley and Baker 2007) for protein docking with backbone minimization. The critical step is choosing residues for our root vertices that are closest to their protein chain’s "middle" residue while also being a part of a non-sampled region (see Supplementary Fig. S1). In the case of multichain proteins, residues that participate in interchain interactions that should be conserved (*e.g.* oligomerization sites) are prioritized. The goal is to make so the structurally conserved non-sampled regions are, whenever possible, "upstream" in the FoldTree, relative to the sampled regions. When sampling flexible regions, a proper FoldTree morphology minimizes the propagation of conformational changes to folded domains, leading to a lower occurrence of unwanted "lever arm effects" and a more efficient sampling protocol.

#### 1.2. Backbone Conformational Sampling

##### 1.2.1. Monte Carlo Acceptance Criterion

Each Monte Carlo move attempted on the current conformation is accepted with a probability calculated using the Metropolis Criterion (Supplementary Equation S1). If the energy of the new conformation is lower than or equal to the energy of the original conformation, the move is always accepted. If the energy of the new conformation is higher than the energy of the original conformation, the move is accepted with a probability expressed by the Boltzmann factor of the energy difference, $\exp(-\Delta E/k_b T)$, where ΔE is the difference between the energies of the new and original conformations, $k_b$ is the Boltzmann constant and $T$ is the temperature parameter. In practice, we take a random number between 0 and 1, and if it is lower than $\exp(-\Delta E/k_b T)$, we accept the move; otherwise, it is rejected (Metropolis *et al.* 1953, Allen and Tildesley 1987). If the move is rejected, a new backbone perturbation with different dihedral angle values is attempted. If after a number of attempts no move is accepted then that residue’s sampling is skipped and its dihedral angles remain unchanged. The use of the Metropolis criterion avoids local minima trapping through the occasional acceptance of worse solutions, resulting in a more efficient exploration of the protein’s conformational landscape.

##### 1.2.2. Application of Structural Restraints

After each Monte Carlo move, the energy of the resulting conformation is evaluated using a scoring function. Conformations that result in higher energy scores are more likely to be rejected when applying the Metropolis Criterion (Supplementary Equation S1). Thus, during conformational sampling of flexible regions, in order to preserve the structure of functionally relevant secondary structure elements, folded domains, or interchain contacts, energy restraints are created between their $C_\alpha$ constituting atoms so that the distance between them does not change significantly.

The creation of a restraint in Rosetta requires deciding what metric it is measuring and then defining how that measurement is translated into a scoring bonus or penalty. Here, we want to measure the distance between a pair of $C_\alpha$, and penalize structures where that distance deviates from its initial value. The more the distance between restrained $C_\alpha$ would change, the higher the value of their restraints, and more likely is the rejection of the Monte Carlo move that originated that change.
The score of a restraint applied to a pair of $C_\alpha$ is calculated using the function $f(x)$ in Supplementary Equation S2. Here, $x_0$ is the initial distance between the pair of $C_\alpha$, $x$ is the distance between the atoms after a Monte Carlo move, and σ is the width parameter of the harmonic potential that controls the strength of the restrain (Supplementary Fig. S2). The score of the restraint becomes higher (worse) the more the distance between the two $C_\alpha$ atoms changes. A tolerance parameter, *tol*, is added to the function definition, adding a flat region to the center of the harmonic potential that allows the distance between two restrained atoms to change before the score of their conformation begins increasing. In practice, this means assigning a range of values of x defined by the tolerance parameter, [$x_0$-*tol*, $x_0$+*tol*], for which the value of the restraint function $f(x)$ is zero.

##### 1.2.3. Optimizing database memory usage

The dihedral angles three-residue fragment (tripeptide) database used in this work was originally created and published by González-Delgado *et al*. As described in (González-Delgado *et al.* 2022), it was built by extracting dihedral angles from structures taken from the SCOPe (Fox, Brenner and Chandonia 2014, Chandonia, Fox and Brenner 2019) 2.07 release, a curated database of high-resolution experimentally determined protein structures. In total, 6,740,433 tripeptide dihedral angle values were extracted, making up the *all* dataset. A structurally filtered dataset, *coil*, was generated by removing tripeptides that participated in α-helices or β-strands, reducing the number of tripeptide dihedral angle values to 3,141,877.
The tripeptide database was originally published as two sets, *all* and *coil*, each containing 20 files, one for each amino acid residue, with the dihedral angles φ, ψ and ω (in radians) of tripeptides where that amino acid was the central residue. The following steps were conducted equally for each of the two datasets, *all* and *coil*.
First, the 20 database files were concatenated into a single Pandas (McKinney 2010) DataFrame, which was then stripped of columns not relevant to our work, with 12 columns remaining: 9 containing the φ, ψ and ω angles for each residue of the triplet (in radians) and 3 with the 3-letter code of each residue making up the fragment, in order. The latter columns were condensed into a single column, with a text string representing the tripeptide fragment (but using the 1-letter code, so fewer bytes were required for storage). This resulted in a DataFrame with 10 columns: 9 containing the φ, ψ and ω angles for each residue of the triplet (in radians), and 1 with the string identification of the fragment they make up using the amino acid 1-letter code.
Finally, since the sampled database had to be kept in memory during computations, it was of utmost importance to reduce its size in memory to the minimum amount possible. This was achieved through the assignment of more appropriate data types to each column of the Pandas DataFrame. For convenience, in a DataFrame, the default data type for numeric values with decimal places is the double-precision floating-point format (*float64*). This format assigns 8 bytes (64 bits) of memory for each value, storing values with a high precision of approximately 16 decimal digits. The single-precision floating-point format, *float32*, assigns

only 4 bytes (32 bits) of memory to each value, resulting in a lower precision of approximately 7 decimal digits. In our case, dihedral angles are stored in radians, which means they are stored in the range [-π, π]. Thus, accurate representation of the decimal places of each value is important to differentiate between them. However, only a small fraction of rows in the database contained values that exceeded 7 decimal places (0.263% in *all* and 0.285% in *coil*), and so only a small number of rows were at the risk of becoming unreliable if we made the swap to the *float32* data type. The low chances for data loss together with the fact that half the computer memory was now required (as described in Supplementary Table S1), motivated the conversion of the columns storing dihedral angle values to the *float32* data type.

The fragment column in the database was originally assigned the *object* data type in the Pandas DataFrame. It is the most versatile Pandas data type, but also the one that allocates the most memory. Fortunately, Pandas offers an alternative that proved useful for our use case. The *category* data type holds an internal mapping that assigns integer values to all the unique values found in a column, and then stores the values in that column in *int8* format, the most space efficient format for integer storage in Pandas. This is only useful if the number of unique values in our column is lower than the total number of values, or else the use of the *category* could actually lead to an increase in memory usage, since we would be including all the original object string identifiers with the addition of all the integer values assigned to them. In our case, the number of unique values in the tripeptide fragment column could never surpass 8000 ($20^3$), which in both cases would be only a small fraction of the total number of values (0.119% in *all* and 0.255% in *coil*). Thus, it was appropriate to convert our fragment column to a *category* data type, which is further supported by the 96.5% reduction in memory size (see Supplementary Table S1).

##### 1.2.4. Increasing Sampling Variability

Two strategies are used to increase the variability of the Monte Carlo sampling on the protein backbone. First, before sampling a target region, the order in which residues within that region will be targeted is randomized. Second, when retrieving values from the dihedral angle database, new dihedral angles are randomly drawn from a Gaussian distribution centred on the database, with a user-defined percentage variance (defaults to 10%).

##### 1.2.5. Using AlphaFold confidence metrics to bias conformational sampling

The present work introduces a method to  convert the structural information contained in the AlphaFold PAE matrix (Guo *et al.* 2022, Brotzakis *et al.* 2025) into energy restraints. By default, the restraints described with Supplementary Equation S2 are only applied between atoms that constitute either non-sampled regions or regions that make up a protein interface. However, when the user decides to use the PAE matrix output by AlphaFold to guide the conformational sampling process, the PAE values are considered instead for the selection of targets for restraint application. First, the PAE matrix is filtered, keeping only values below a user-defined cutoff (default is 10 Å). Then, distance restraints are applied between each pair of residues found in the filtered matrix, with the value for the σ parameter in the restraint function being equal to the PAE value of that residue pair (Supplementary Equation S3). Since the PAE values assigned to residue pairs that make up folded domains are usually very low (< 2-3 Å), the intra-domain pairwise restraints required for conserving the structure of folded domains during sampling are always included. The strength of each added restraint scales with the PAE value of the residue pair it is applied to, as described in Supplementary

Figs. S3, S4 and Supplementary Table S2. The larger the error associated with the relative position of a residue pair, the less influence the violation of that restraint has on increasing the conformation's score. The value for the pLDDT of both residues involved in a PAE-based restraint is also considered: when one residue has high pLDDT and the other has low pLDDT and either one of them will be sampled, the restraints between them are made weaker through a $\gamma$ scaling parameter (default is 30). This is done by simply multiplying the PAE value for that residue pair by $\gamma$ prior to restraint creation (see Supplementary Equation S3 and Supplementary Table S2, Scenario 2). This ensures that regions of low local confidence retain higher conformational freedom while still remaining tethered to positions consistent with AlphaFold's PAE metric.
To avoid overspecification of rigid restraints when many PAE values fall below the cutoff, a "flatten" option can be applied, replacing all PAE values below the cutoff with a uniform value. This promotes permissive restraints that facilitate adequate conformational sampling while maintaining structural coherence.

### 2. Bayesian/Maximum Entropy (BME) Ensemble Reweighting

A Bayesian/Maximum Entropy (BME) approach is used to reweight generated ensembles so calculated averages are closer to experimental values while minimally changing the original ensemble. As described in (Bottaro, Bengtsen and Lindorff-Larsen 2018), this can be done by finding the set of optimal weights that minimizes the function $\mathcal{L}$ described in Supplementary Equation S4, where $(w_1 \ldots w_n)$ are the statistical weights assigned to each structure of the ensemble, $\chi^2$ measures the agreement with experimental data (Supplementary Equation S5), $S_{REL}$ measures the deviation from the initial set of weights $w^0$ (Supplementary Equation S6), $m$ is the number of experimental measurements, $n$ is the size of the generated ensemble and $\theta$ is a scaling parameter. The reduced $\chi^2$ calculation follows Supplementary Equation S5, where $F_i^{EXP}$ is the experimental measurement, $F_i(x_j)$ is the theoretical measurement calculated from each conformer, and $\sigma_i^2$ is the uncertainty associated with $F_i^{EXP}$ which includes experimental errors and inaccuracies introduced by the calculation of $F_i(x_j)$. Ensemblify supports the use of SAXS data for ensemble reweighting, for which the reduced $\chi^2$ is often used as a method of comparing calculated to experimental data (Gomes *et al.* 2020).
The $\theta$ parameter must be adequately tuned so that the reweighted ensemble has good agreement with experimental data (low $\chi^2$) while conserving the most information possible contained in the initial ensemble (high $S_{REL}$). The reweighting procedure is repeated for different values of $\theta$, starting from a set of initial weights $w^0 = 1/n$ for each structure, where $n$ is the size of the generated ensemble. At high values of $\theta$, the changes to the initial weights are minimal and the agreement to experimental data is lowest. As $\theta$ is decreased, the weights change to find a combination that improves the agreement with experimental data (decrease $\chi^2$), becoming less uniform. The decrease in uniformity of the weights corresponds to a decrease in the number of structures that effectively contribute to the calculated averages, and can be quantified by the fraction of effective frames (in our case, structures), $\phi_{eff}$ (Supplementary Equation S7).
Hence, it is useful to plot $\phi_{eff}$ versus $\chi^2$ to evaluate the balance between a good agreement with experimental data (low $\chi^2$) and the minimal perturbation of the initial ensemble (high $\phi_{eff}$). In practice, a good value of $\theta$ will be located at the point of maximum curvature, *i.e.* the "elbow" of this plot (highlighted data point in Supplementary Fig. S5).

### Supplementary Results

#### 3.1. Ensemblify offers user-friendly, interactive interfaces

#### 3.2. Ensemblify is applicable to multiple protein structural architectures

A set of proteins with varying size and structural architectures (Supplementary Figure S10) was used to test the developed framework, showcasing its ability to sample N- and C-terminal tails and inter-domain linkers, while conserving the structure of folded domains and, in the case of multichain proteins, interchain protein interfaces. Tested proteins were divided into groups according to their structural architecture: full IDPs (Supplementary Fig. S10A), folded domains connected by flexible linkers (Supplementary Fig. S10B), folded domains connected to long flexible tails (Supplementary Fig. S10C) and multichain proteins (Supplementary Fig. S10D).

For each of the tested proteins, after generating a conformational ensemble, averaged ensemble properties were calculated. These included average distance matrices, secondary structure assignment frequency plots and probability distributions for structural metrics like the radius of gyration ($R_g$), maximum $C_\alpha$ distance ($D_{max}$) and end-to-end distance ($D_{ee}$).

Subsequently, each ensemble was reweighted using a Bayesian/Maximum Entropy (BME) approach (Supplementary Information S2) using experimental SAXS data measured for each protein. The theoretical SAXS profiles calculated from both the uniformly weighted and reweighted ensembles were fitted to experimental SAXS profiles and their agreement was compared. In most cases, reweighting significantly improved agreement with experimental data. For some of the tested protein systems, the agreement with experimental data prior to reweighting was already quite good, so after reweighting the improvement was not as significant. Despite these differences, in all cases the reweighted ensemble retained a significant amount of information from the uniformly weighted ensemble, as shown by the average value of 0.65 for the fraction of effective structures ($\phi_{\mathrm{eff}}$) across all protein systems. Finally, averaged ensemble structural properties were recalculated using the new weights for each conformer, and compared with the ones calculated from the uniformly weighted ensembles, enabling a quantitative view of the effect of ensemble reweighting.

Time taken to generate an ensemble of 10000 structures for each protein construct is described in Supplementary Tables S3 and S4, along with references for the sampling starting structures and experimental SAXS data used in BME reweighting. Ensemble generation took between ~5 minutes and ~3 days, with proteins with a larger number of residues generally taking more time. This is expected, since a larger number of sampling targets require more Monte Carlo moves, with more energy calculations required for each move. Moreover, if folded domains whose structure should remain conserved are also larger, each move will be accompanied by a greater number of energy calculations when determining the possible violation of spatial restraints.

#### 3.2.1. Fully intrinsically disordered proteins

##### 3.2.1.1. Histatin 5

Histatin5 (Hst5) is a fully disordered, histidine rich antimicrobial peptide found naturally in human saliva that does not show any distinct structure in aqueous solutions (Oppenheim *et al.* 1988). However, it adopts an α-helical conformation when dissolved in organic solvents commonly used in protein structural studies, such as dimethylsulfoxide (DMSO) and trifluoroethanol (TFE) (Melino *et al.* 1999, Raj, Marcus and Sukumaran 1998). It has also been shown that the disruption of this α-helix, particularly in the 14-19 region, leads to a significant reduction in Hst5's candidacidal activity, highlighting the importance of this secondary structure for the protein's function (Raj, Marcus and Sukumaran 1998).

The averaged structural properties calculated over the conformational ensemble generated for Hst5, before and after BME reweighting, are summarized in Supplementary Fig. S11. Reweighting improved the agreement with SAXS experimental data, as denoted by a reduction of the $\chi^2$ value from 1.98 to 1.05 (Supplementary Fig. S11A). The relatively low $\chi^2$ value prior to reweighting indicates that the behaviour of Hst5 in solution seemed to be already somewhat captured through the uniform ensemble. The fitting of theoretical SAXS curves calculated from the uniform and reweighted ensemble to the experimental SAXS profile supports this claim, as in both cases the fitting is quite similar (Supplementary Fig. S11B).

The probability distributions for the $R_g$, $D_{max}$ and $D_{ee}$ calculated from the Hst5 uniform and reweighted ensembles are shown in Supplementary Fig. S11C. Comparing them between the uniform and reweighted ensembles shows that, in all cases, the reweighted ensemble favours more extended conformations, with both the distributions and their averages shifting towards higher values. The average value for the $R_g$ was found to be 11.54±0.02 Å for the uniform ensemble and 12.71±0.03 Å for the reweighted ensemble. These values are both in accordance with values found in the literature for atomistic molecular dynamics simulations of Hst5 using different force fields, which lie in the 11-13 Å interval (Fagerberg 2022, Jephthah *et al.* 2021, Henriques, Cragnell and Skepö 2015, Shrestha, Smith and Petridis 2021, Thomasen *et al.* 2024). It has been shown elsewhere that an ensemble comprised of more extended conformations of Hst5 (average $R_g$ between 12-13 Å) shows better agreement with experimental SAXS data (Shrestha, Smith and Petridis 2021), which is in line with what was observed here for the reweighted ensemble.

The distance matrices calculated from the Hst5 uniform and reweighted ensembles are shown in Supplementary Fig. S11D. The distance matrix for the uniform ensemble is typical of fully flexible proteins, as there are no apparent preferences for inter-residue distances throughout the sequence. After reweighting, more extended conformations are favoured, increasing the average distance between residue pairs that are far apart in the protein sequence by 4-6 Å.

The secondary structure assignment frequency plots calculated from the Hst5 uniform and reweighted ensembles are shown in Supplementary Fig. S11E. The regions around residues 3-7 and 14-19 are assigned an α-helical secondary structure in ~20% of structures, while the rest of the protein is consistently classified as a random coil. These results show that in the 4-24 region of Hst5 there seems to be a structural propensity to form α-helices, which is in line with what has been observed experimentally (Raj, Marcus and Sukumaran 1998, Melino *et al.* 1999).

##### 3.2.1.2. α-synuclein

α-synuclein (aSyn) is a fully disordered cytosolic protein enriched in presynaptic terminals. Under physiological conditions, aSyn is normally in an unfolded state, but adopts an α-helical conformation in the presence of membranes with acidic phospholipid head-groups and/or high curvature (Tofaris 2022, Chen *et al.* 2021, Hijaz and Volpicelli-Daley 2020). The accumulation of aggregated forms of aSyn in neuronal cells is a hallmark of synucleinopathies like Parkinson's disease (Volles and Lansbury 2003). The structure of aSyn is commonly divided into three domains: the N-terminal domain (residues 1-87), the non-amyloid component (NAC) domain (residues 61-95) and the C-terminal domain (residues 96-140) (Emamzadeh 2016). The N-terminal domain is a positively charged region that has a tendency to form α-helical structures when interacting with lipid bilayers (Giasson *et al.* 2001).

The averaged structural properties calculated over the conformational ensemble generated for aSyn, before and after BME reweighting, are summarized in Supplementary Fig. S12. The uniform aSyn ensemble was reweighted with BME using a value of 75 for the $\theta$ parameter, which improved the agreement with SAXS experimental data, as denoted by a reduction of the $\chi^2$ value from 1.19 to 1.04 (Supplementary Fig. S12A). The low $\chi^2$ value prior to reweighting indicates that the behaviour of aSyn in solution seemed to be already relatively well captured through the uniform ensemble. The fitting of theoretical SAXS curves calculated from the uniform and reweighted ensemble to the experimental SAXS profile supports this claim, as in both cases the fitting is quite similar (Supplementary Fig. S12B), although there is a marginal improvement for the reweighted ensemble.

The probability distributions for the $R_g$, $D_{max}$ and $D_{ee}$ calculated from the aSyn uniform and reweighted ensembles are shown in Supplementary Fig. S12C. Comparing them between the uniform and reweighted ensembles shows that, in all three cases, there is only a slight increase in the average values for the reweighted ensemble, with only minor changes to the shape of the distributions. The general similarity between the uniform and reweighted ensembles makes sense considering that the uniform ensemble already had a great fit to SAXS data ($\chi^2$ = 1.19). The average value for the $R_g$ was found to be 35.93±0.07 Å for the uniform ensemble and 36.15±0.12 Å for the reweighted ensemble. These values are both in accordance with values found in the literature for aSyn, which include coarse-grained molecular dynamics simulations with rescaled force fields, with average $R_g$ values in the 30-40 Å interval (Thomasen *et al.* 2024, Ramis *et al.* 2019, Pan, Mu and Chen 2023), and an ensemble of aSyn refined with NMR data, with an average $R_g$ also in the 30-40 Å interval (Schwalbe *et al.* 2014). Additionally, the average $R_g$ value calculated from the uniform ensemble (35.93±0.07 Å) is quite close to the $R_g$ value from experimental SAXS data (~35Å) (Piana *et al.* 2015).

The distance matrices calculated from the aSyn uniform and reweighted ensembles are shown in Supplementary Fig. S12D. After reweighting, conformations that are very slightly more extended are favoured, increasing the average distance between the N- and C-terminal regions by ~2 Å. This matches the results obtained for the structural metrics probability distributions, where the reweighted ensemble shows a preference for slightly more extended conformations.

The secondary structure assignment frequency plots calculated from the aSyn uniform and reweighted ensembles are shown in Supplementary Fig. S12E. In some regions, up to 20-30% of structures are assigned an α-helical secondary structure, with residue 19 having the highest frequency of α-helical assignment (~35%). This result is in agreement with what was concluded in a structural study of α-synuclein using NMR, which marked the 6-37 region as having a strong tendency to form α-helices (Eliezer *et al.* 2001).

##### 3.2.2. Multidomain proteins connected by flexible linkers

###### 3.2.2.1. Cardiac myosin-binding protein C

Myosin-binding protein C (MyBP-C) is a protein that plays important structural and regulatory roles in the muscle sarcomere by interacting with myosin and actin. MyBP-C is expressed in multiple isoforms in striated muscles, with the cardiac isoform (cMyBP-C) being linked to hypertrophic cardiomyopathy, a prevalent disease which can cause disability or death due to heart failure or stroke (Doh *et al.* 2022). cMyBP-C is a large, highly modular protein, composed of 11 domains (C0-C10): 8 immunoglobulin-like domains (C0-C5, C7, C10) and 3 fibronectin-type III domains (C6, C8, C9). The C1 and C2 domains are linked by ~100 residues comprising the “myosin binding motif” or m-domain, which is mostly unstructured except for a small tri-helix bundle (THB) at its C-terminal end (Howarth *et al.* 2012). The THB of the m-domain is connected to the C2 domain by a short flexible linker of 7 residues, and both the THB and the linker are highly conserved among MyBP-C isoforms. Many mutations occurring in the THB to C2 region can lead to pathology, suggesting an important role in cMyBP-C function (Michie *et al.* 2016).

In this work, a conformational ensemble was generated for a construct containing the THB of the m-domain connected to the C2 domain by the small disordered linker (cMyBP-C$_{mTHB\text{-}C2}$) (see Supplementary Fig. S10B). This construct is composed of 137 residues, containing a small 5 residue flexible tail (from truncating the m domain) followed by the THB (residues 6-42), the short disordered linker (residues 43-49) and the C2 domain (residues 50-137).

The averaged structural properties calculated over the conformational ensemble generated for cMyBP-C$_{mTHB\text{-}C2}$, before and after BME reweighting, are summarized in Supplementary Fig. S13. The uniform cMyBP-C$_{mTHB\text{-}C2}$ ensemble was reweighted with BME using a value of 400 for the $\theta$ parameter, which improved the agreement with SAXS experimental data, as denoted by a reduction of the $\chi^2$ value from 6.73 to 1.07 (Supplementary Fig. S13A). The fitting of the theoretical SAXS curve calculated from the reweighted ensemble was significantly better compared to the uniform ensemble, especially for high values of $q$ (Supplementary Fig. S13B).

The probability distributions for the $R_g$, $D_{max}$ and $D_{ee}$ calculated from the cMyBP-C$_{mTHB\text{-}C2}$ uniform and reweighted ensembles are shown in Supplementary Fig. S13C. Comparing them between the uniform and reweighted ensembles shows that the reweighted ensemble favours more compact conformations, with both the shape and averages of the distributions shifting towards lower distance values. Additionally, it is clear by looking at the $R_g$ and $D_{max}$ distributions that more compact conformations are being more populated, denoted by a clear peak located at smaller distance values. The obtained $R_g$ distributions show values between 14-28 Å, with an average of 20.23±0.03 Å for the uniform ensemble and 18.65±0.04 Å for the reweighted ensemble. The $D_{max}$ distributions show values between 40-85 Å, with an average of 62.19±0.09 Å for the uniform ensemble and 57.13±0.15 Å for the reweighted ensemble. For both the $R_g$ and $D_{max}$, the shape and average value of the distributions obtained for the uniform ensemble match the results obtained elsewhere for an ensemble of the same cMyBP-C$_{mTHB\text{-}C2}$ construct, derived from NMR structures (Michie *et al.* 2016). In that study, the obtained $R_g$ distribution presented values between 15-28 Å, with an average of ~20 Å, while the $D_{max}$ distribution presented values between 45-90 Å, with an average of

~62.5 Å. Additionally, in the same study, the results obtained after applying a different reweighting method to the ensemble were similar to the ones obtained here after BME reweighting. There was an increase in the population of compact structures, with a clear peak in the reweighted distributions around ~18 Å for the $R_g$ and ~57 Å for the $D_{max}$ (Michie *et al.* 2016). The results obtained here for the $R_g$ distribution also match other results found in the literature for an ensemble of the same cMyBP-$C_{mTHB\text{-}C2}$ construct, generated using molecular dynamics simulations, which showed values between 15-28 Å and an average $R_g$ of ~21 Å (Thomasen *et al.* 2024).

The distance matrices calculated from the uniform and reweighted ensembles of cMyBP-$C_{mTHB\text{-}C2}$ are shown in Supplementary Fig. S13D. The distance matrix for the uniform ensemble clearly represents the folded THB region (residues 6-42), followed by the short disordered linker (residues 43-49) and finally the larger C2 structured domain (residues 50-137). Reweighting favours conformations that place the THB and C2 domains closer together, decreasing the average distance between the two domains by 4-6 Å.

The secondary structure assignment frequency plots calculated from the cMyBP-$C_{mTHB\text{-}C2}$ uniform and reweighted ensembles are shown in Supplementary Fig. S13E. They clearly indicate the presence of three α-helical regions in the THB conserved among all structures, as well as a region majorly composed of β-sheets and a small α-helical region, characteristic of the C2 domain, which is also conserved. The small flexible region linking the THB to the C2 domain seems to have some tendency to adopt an α-helical conformation (seen in ~20% of structures), aligning with predictions of the secondary structure propensity of this region (Michie *et al.* 2016).

###### 3.2.2.2. Tetraubiquitin

Ubiquitin (Ub) is a highly conserved 76 residue protein that can be found in most tissues of eukaryotic organisms. Ubiquitination is a reversible post-translational modification in which an Ub monomer is attached to a substrate protein. Ub functions as a molecular tag that marks proteins for either degradation by the proteasome or to participate in a wide variety of other functions (DNA repair, transcription, endocytosis, membrane transport, protein localization and antigen processing), in a proteasome independent manner (Ande, Chen and Maddika 2009). During ubiquitination, the C-terminus of a Ub monomer is covalently attached to either a lysine or the N-terminus of a substrate protein. Then, the Ub monomer itself can be ubiquitinated at one of its seven lysine residues (K6, K11, K27, K29, K33, K48, and K63) or its N-terminal methionine residue (M1). This can lead to the formation of a polyubiquitin chain, whose length and topology ultimately determine its biological signal within the cell . A polyubiquitin chain in which all Ub monomers are connected through their M1 residues is known as a linear ubiquitin chain. Several proteins have been described to interact with linear ubiquitin chains via specific Ub-binding domains (Jung *et al.* 2024).

Characterizing the conformational space of polyubiquitin chains is essential for understanding their physiological behaviour. In this work, an ensemble was generated for linear tetraubiquitin ($Ubq_4$), a polyubiquitin chain with four Ub monomers connected through their M1 residues. This construct has 304 amino acid residues, with four Ub domains (residues 1-72, 77-148, 153-224 and 229-300), connected by short 4 residue linkers, with the last domain being followed by a short 4 residue flexible tail (which would link the possible next Ub monomer).

The averaged structural properties calculated over the conformational ensemble generated for $Ubq_4$, before and after BME reweighting, are summarized in Supplementary Fig. S14. The uniform $Ubq_4$ ensemble was reweighted with BME using a value of 50 for the $\theta$

parameter, which improved the agreement with SAXS experimental data, as denoted by a reduction of the $\chi^2$ value from 2.03 to 1.22 (Supplementary Fig. S14A). The relatively low $\chi^2$ value prior to reweighting indicates that the behaviour of $Ubq_4$ in solution seemed to be already somewhat captured through the uniform ensemble. The fitting of theoretical SAXS curves calculated from the uniform and reweighted ensembles to the experimental SAXS profile supports this claim, as in both cases the fitting is quite similar (Supplementary Fig. S14B).

The probability distributions for the $R_g$, $D_{max}$ and $D_{ee}$ calculated from the $Ubq_4$ uniform and reweighted ensembles are shown in Supplementary Fig. S14C. Comparing them between the uniform and reweighted ensembles shows that in all cases there are minimal differences. However, there is a minor preference in the reweighted ensemble for slightly more extended conformations, which can be seen by both the average values and the distributions shifting slightly towards higher distance values. The slight preference for more extended conformations of $Ubq_4$ after reweighting could be due to the fact that this type of Ub oligomerization usually leads to more extended, "open" conformations, as opposed to the ones found in a K48-linked Ub chain (Jussupow *et al.* 2020). For the uniform ensemble, the obtained $R_g$ distribution shows values between 20-43 Å, with an average of 32.34±0.04 Å, while the $D_{ee}$ distribution shows values between 10-150 Å, with an average of 89.179±0.24 Å. For both of these metrics, similar results can be found in other studies, where ensembles of $Ubq_4$ were generated using coarse-grained molecular dynamics simulations (Jussupow *et al.* 2020, Thomasen *et al.* 2024). In these studies, values of $R_g$ were found between 18-43 Å, with averages close to ~33 Å, with $D_{ee}$ values between 10-170 Å, with an average around ~90 Å.

The distance matrices calculated from the $Ubq_4$ uniform and reweighted ensembles are shown in Supplementary Fig. S14D. The distance matrix for the uniform ensemble clearly differentiates between the four Ub monomers, and it is possible to see that the distance between domains increases the further apart their positions are in the ubiquitin chain. After reweighting, conformations that are slightly more extended are favoured, with distances Ub(1-3) and Ub(1-4) increasing by 2-4 Å.

The secondary structure assignment frequency plots calculated from the $Ubq_4$ uniform and reweighted ensembles are shown in Supplementary Fig. S14E. They clearly indicate the presence of four Ub domains, whose structure is conserved among all conformers. There do not seem to be any significant propensities for regular secondary structure in any of the linker regions.

###### 3.2.2.3. T-cell intracellular antigen-1

T-cell intracellular antigen-1 (TIA1) is an RNA-binding protein that has multiple roles within cells, including the regulation of alternative splicing of various pre-mRNA and the mediation and suppression of mRNA translation in the cytoplasm during eukaryotic cellular stress response (Wang *et al.* 2014, Fritzsching *et al.* 2020). Native TIA1 has three RNA recognition motifs (RRM1, RRM2, and RRM3) connected by flexible linkers and followed by a C-terminal disordered domain of ~100 residues enriched in glutamine and asparagine (Q-rich domain). A mutation within the Q-rich domain is a diagnostic marker of Welander distal myopathy, and several other TIA1 mutations are related with various pathologies, including amyotrophic lateral sclerosis (Fritzsching *et al.* 2020). Despite its importance, little is known about the disordered Q-rich domain and its effect on the full length structure, mainly due to experimental challenges. The lack of experimental SAXS data that included this domain forced the use of a truncated construct of TIA1 in this work, without the Q-rich domain, which will now be referred simply as TIA1 (Sonntag *et al.* 2017). This construct has 275 residues, being comprised of a short flexible 5 residue tail, followed by the RRM1, RRM2 and RRM3 motifs (residues 6-82, 95-172 and 190-275, respectively), with a 12 residue flexible linker connect RRM1 and RRM2 and a 17 residue flexible linker connecting RRM2 and RRM3.

The averaged structural properties calculated over the conformational ensemble generated for TIA1, before and after BME reweighting, are summarized in Supplementary Fig. S15. The uniform TIA1 ensemble was reweighted with BME using a value of 200 for the $\theta$ parameter, which improved the agreement with SAXS experimental data, as denoted by a reduction of the $\chi^2$ value from 7.25 to 1.24 (Supplementary Fig. S15A). The fitting of theoretical SAXS curves calculated from the reweighted ensemble was significantly better compared to the uniform ensemble, especially for low values of $q$ (Supplementary Fig. S15B).

The probability distributions for the $R_g$, $D_{max}$ and $D_{ee}$ calculated from the TIA1 uniform and reweighted ensembles are shown in Supplementary Fig. S15C). Comparing them between the uniform and reweighted ensembles shows that in all cases there are minimal differences. For the $R_g$ there are small noticeable differences between the two distributions: while both show values between 18-44 Å, the reweighted one slightly favours more compact conformations. This leads to a change of the average $R_g$ value from 27.83±0.03 Å in the uniform distribution to 27.28±0.05 Å in the reweighted distribution. The average $R_g$ value for the uniform ensemble matches experimental SAXS data of TIA1 (Sonntag *et al.* 2017). Additionally, the results obtained for the $R_g$ distribution of the uniform ensemble match ones found for ensembles of TIA1 generated from coarse-grained molecular dynamics simulations using optimized force fields, which also presented values between 18-44 Å, with averages around 28 Å (Larsen *et al.* 2020, Thomasen *et al.* 2024). This highlights the ability of the present work to generate, without specific parametrization, results that are par with other approaches optimized for disordered regions.

The distance matrices calculated from the TIA1 uniform and reweighted ensembles are shown in Supplementary Fig. S15D. The distance matrix for the uniform ensemble clearly shows the differentiation between the RRM1, RRM2 and RRM3 domains of TIA1, connected by flexible linkers. After reweighting, we can detect small changes in the distances between domains, with RRM1-RRM2 and RRM2-RRM3 becoming closer together by ~2 Å, while RRM1-RRM3 move further apart by ~1 Å. Although these values are quite small, the tendency for the distance between the RRM1 and RRM3 domains to increase after BME reweighting using SAXS data has also been observed elsewhere (Larsen *et al.* 2020). This is counteracted by the approximation of the RRM1-RRM2 and RRM2-RRM3 domains, which results in a more globally compact structure, as denoted by the aforementioned reduction in the average $R_g$ value.

The secondary structure assignment frequency plots calculated from the TIA1 uniform and reweighted ensembles are shown in Supplementary Fig. S15E. They clearly highlight the structural pattern characteristic of the RRM domains, with alternating β-sheets and α-helices, and this structure is conserved among all conformers. The flexible linkers do not seem to have any significant secondary structure tendencies, with α-helices being assigned only in ~5% of conformers.

###### 3.2.2.4. “Mothers against decapentaplegic homolog 4”

The “mothers against decapentaplegic homolog 4” (SMAD4) belongs to a family of tumor suppressor genes that act as the principal regulators of the transforming growth factor beta (TGF-β) signalling pathway. The activation of TGF-β receptors at the cellular membrane by the TGF-β hormone can produce opposite effects, such as the inhibition of cell proliferation or the promotion of cell growth. TGF-β signalling is therefore context-dependent, and is linked to the activity of SMAD proteins, including SMAD4, which acts as a mediator in the transmission of the cellular signal from the membrane receptor to the cell nucleus, mainly through oligomeric association with other SMAD proteins (Shah *et al.* 2023).

The translated SMAD4 protein has three domains: the Mad Homology 1 (MH1) domain at the N-terminus, the Mad Homology 2 (MH2) domain at the C-terminus, and a disordered linker region connecting the MH1 and MH2 domains. The MH1 domain binds DNA, while the MH2 domain and the linker region can interact with receptors, regulatory proteins and transcription co-factors, determining the outcome of TGF-β signalling (Gomes 2019). The MH1 and MH2 domains are closely linked: MH1 can disturb the interaction of MH2 with other SMAD proteins, while MH2 can prevent the binding of MH1 to cDNA (Shah *et al.* 2023). The linker region contains a SMAD-activation domain (SAD) at its C-terminal region, which is necessary for the activation of SMAD-dependent transcriptional responses (de Caestecker *et al.* 2000). Mutations in SMAD4 are found in several hereditary cancer predisposition syndromes and can promote the growth of cancer cells, being often observed after significant tumour progression, during metastasis.

Characterizing the conformational landscape of full-length SMAD4 is essential for a better understanding of its inter-domain dynamics, which could affect its oligomerization behaviour. Although SAD is part of the disordered linker of SMAD4, its structure was fixed during conformational sampling, as constructs with the structure of this domain in a folded state have been shown to more accurately conform with experimental SAXS data (Gomes 2019). In this work an ensemble was generated for a construct of full-length SMAD4, composed of 554 residues and containing the MH1 domain (residues 1-140) and SADMH2 domain (residues 271-554), connected by a long flexible linker.

The averaged structural properties calculated over the conformational ensemble generated for SMAD4, before and after BME reweighting, are summarized in Supplementary Fig. S16. The uniform SMAD4 ensemble was reweighted with BME using a value of 200 for the $\theta$ parameter, which improved the agreement with SAXS experimental data, as denoted by a reduction of the $\chi^2$ value from 4.37 to 1.09 (Supplementary Fig. S16A). The fitting of the theoretical SAXS curves calculated from the uniform and reweighted ensembles to experimental SAXS data highlights this improvement, especially for low values of $q$ (Supplementary Fig. S16B).

The probability distributions for the $R_g$, $D_{max}$ and $D_{ee}$ calculated from the SMAD4 uniform and reweighted ensembles are shown in Supplementary Fig. S16C. Comparing them between the uniform and reweighted ensembles shows that in all cases, despite no significant changes in the average values of the distributions, there are noticeable changes in their shape. In the reweighted distributions we can observe that both more compact and more extended conformations were more highly populated, with less importance given to conformers in the middle. This seems to indicate that the experimental SAXS data of SMAD4 can be better explained by a separation of conformations into two distinct groups: one more compact, where the MH1 and MH2 domains are closer together, and one more extended, where they are further apart. These results align with what has been shown in structural studies of SMAD4, which concluded that it exists in an "open-closed" equilibrium of conformations (Gomes *et al.* 2021, Gomes 2019).

The distance matrices calculated from the SMAD4 uniform and reweighted ensembles are shown in Supplementary Fig. S16D. The distance matrix for the uniform ensemble clearly shows the differentiation between the MH1 and SADMH2 domains, connected by a flexible

region. After reweighting, we can detect small changes in the distances between domains, with MH1 and SADMH2 being further away from each other by ~1 Å, while the linker region seems to adopt a more extended conformation near the SADMH2 region (+2 Å) and a more compact one near the MH1 domain (-1 Å).
The secondary structure assignment frequency plots calculated from the SMAD4 uniform and reweighted ensembles are shown in Supplementary Fig. S16E. The shown structural pattern of β-sheets and α-helices is consistent with the structure of the MH1 and MH2 domains, while the SAD domain's characteristic β-sheet architecture is also clearly represented. For the flexible linker, there seems to be a minor structural propensity to form α-helices, with this secondary structure being assigned throughout the region in around 15% of structures.

##### 3.2.3. Proteins with folded domains connected to long flexible tails

###### 3.2.3.1. Non-receptor tyrosine kinase Src

The Src family of non-receptor tyrosine kinases (SFK) is formed by at least nine members (Src, Fyn, Yes, Yrk, Fgr, Hck, Lyn, Blk, and Lck), implicated in cell signalling pathways related to cell growth, migration, invasion, and survival (Arbesú *et al.* 2017). Src, the most prominent member of the SFK family, plays a major role in cellular transduction pathways such as cell growth, differentiation, transcription, proliferation, adhesion, and survival and is associated with a variety of human cancers (Shrestha *et al.* 2019). SFKs share a common structural architecture with three folded domains (SH1, SH2 and SH3) and an N-terminal IDR that includes the SH4 domain and the Unique domain (UD). The latter is highly heterogeneous among SFK members, varying in both length and sequence. The structure and function of all three conserved domains of Src has been extensively studied: the SH1 domain contains the kinase catalytic centre, while SH2 and SH3 are both regulatory domains. On the other hand, the function of the disordered regions is still not thoroughly understood, nor its relation to the folded domains (Arbesú *et al.* 2017). It is known that SH4 regulates Src cellular localization by binding to the cellular membrane, while the UD mediates interactions with lipids, receptors and protein targets (Shrestha *et al.* 2019). Both the UD and SH4 regions are constrained around the SH3 domain but retain high flexibility, characteristic of a fuzzy intramolecular complex mediated by transient interactions. In this work a conformational ensemble was generated for a construct of Src containing only the disordered region connected to the SH3 domain (USH3). This construct was 154 residues long, composed of the disordered SH4 and Unique domains (residues 1-85), followed by the SH3 folded domain (residues 86-154).
The averaged structural properties calculated over the conformational ensemble generated for USH3, before and after BME reweighting, are summarized in Supplementary Fig. S17. The uniform USH3 ensemble was reweighted with BME using a value of 400 for the $\theta$ parameter, which improved the agreement with SAXS experimental data, as denoted by a reduction of the $\chi^2$ value from 5.71 to 1.10 (Supplementary Fig. S17A). The fitting of the theoretical SAXS curves calculated from the uniform and reweighted ensembles to experimental SAXS data highlights this improvement, especially for low values of $q$ (Supplementary Fig. S17B).
The probability distributions for the $R_g$, $D_{max}$ and $D_{ee}$ calculated from the USH3 uniform and reweighted ensembles are shown in Supplementary Fig. S17C. Comparing them between the uniform and reweighted ensembles shows that, in all three cases, the reweighted

ensemble favours more compact conformations, with the averages shifting towards lower distance values. The $R_g$ distribution for the uniform ensemble shows values between 18-52 Å, with an average of 29.46±0.06 Å. These results agree with what was found in a study where an ensemble of USH3 was generated using residue-specific torsion angle potentials, where $R_g$ values were between 20-52 Å with an average of ~30 Å (Arbesú *et al.* 2017). In the same study, a sub-ensemble that was more representative of experimental SAXS data was selected, and the $R_g$ distribution obtained after selection placed more importance on compact conformations, conveyed by the appearance of a clear peak in the distribution near ~20 Å. This was similar to the results obtained here after BME reweighting, where the reweighted distribution also shifted, showing a clear peak near ~23Å. Additionally, the $R_g$ distribution obtained here for the uniform ensemble of USH3 presents, in general, more extended conformations when compared to distributions of $R_g$ derived from ensembles of full length Src, generated using residue-specific torsion angle potentials (20-45 Å) or molecular dynamics simulations (26-32 Å) (Bernadó *et al.* 2008, Strelkova *et al.* 2024). This seems to indicate that the disordered region composed of the SH4 and Unique domains adopts more compact conformations when included in the structural context of the full length protein.

The distance matrices calculated from the USH3 uniform and reweighted ensembles are shown in Supplementary Fig. S17D. The distance matrix for the uniform ensemble clearly differentiates between the flexible SH4 and Unique domains at the N-terminus, followed by the folded SH3 domain at its C-terminus. After reweighting, conformations where the disordered tail is slightly closer to the SH3 domain seem to be preferred, decreasing the average distance between domains by ~2 Å, which is in line with the changes observed after reweighting for the $R_g$ distribution. These results agree with a structural study of USH3, which showed that the disordered N-terminal region is compacted around the SH3 domain (Arbesú *et al.* 2017).

The secondary structure assignment frequency plots calculated from the USH3 uniform and reweighted ensembles are shown in Supplementary Fig. S17E. They clearly indicate the presence of a disordered region followed by a folded domain, conserved among all conformers. The long disordered tail seems to show some propensity to form α-helices, especially in its N-terminal region, with some residues being assigned this secondary structure in close to 20% of conformers. These results align with a study where an ensemble of the disordered tail was generated using molecular dynamics, which identified several areas across the disordered sequence with tendency to form α-helices, near lipid-binding regions of SH4 (Shrestha *et al.* 2019).

###### 3.2.3.2. Galectin-3

Galectins are a family of lectins that bind to glycoconjugates containing β-galactose through their conserved carbohydrate recognition domain (CRD). Galectins are expressed in many animal tissues, being synthesized in the cytoplasm and exported to the extracellular medium. There, they can induce aggregation of or form lattices with glycoproteins or glycolipids on the cell surface, thus regulating cell activation, migration, adhesion, and signalling. Galectin-3 (Gal-3) is unique among the galectin protein family, with a structure containing an intrinsically disordered proline-rich N-terminal tail (NT) connected to a single CRD. While Gal-3 is mainly found in its monomeric form in solution, binding to cell surface glycoconjugates induces its oligomerization through mechanisms which are still poorly understood. Although initially proposed to occur through either CRD-CRD or NT-NT interactions, more recently it has been suggested that NT-CTD interactions are also essential for Gal-3 self-association (Ahmad *et al.* 2004, Lepur *et al.* 2012, Ippel *et al.* 2016,

Flores-Ibarra *et al.* 2018, Pally and Bhat 2021, Sato 2023a, Sato 2023b). Additionally, the fact that Gal-3 undergoes liquid-liquid phase separation is also thought to contribute to its oligomerization behaviour (Lin *et al.* 2017). In this work, an ensemble of conformations was generated for a construct of full length Gal-3, containing its disordered proline-rich N-terminal tail (NT) (residues 1-116) followed by its carbohydrate recognition domain (CRD) (residues 117-250).

The averaged structural properties calculated over the conformational ensemble generated for Gal-3, before and after BME reweighting, are summarized in Supplementary Fig. S18. The uniform Gal-3 ensemble was reweighted with BME using a value of 400 for the $\theta$ parameter, which significantly improved the agreement with SAXS experimental data, as denoted by a reduction of the $\chi^2$ value from 20.16 to 1.5 (Supplementary Fig. S18A). The fitting of the theoretical SAXS curves calculated from the uniform and reweighted ensembles to experimental SAXS data clearly highlights this improvement across all values of $q$ (Supplementary Fig. S18B).

The probability distributions for the $R_g$, $D_{max}$ and $D_{ee}$ calculated from the Gal-3 uniform and reweighted ensembles are shown in Supplementary Fig. S18C. Comparing them between the uniform and reweighted ensembles shows that, in all three cases, the reweighted ensemble favours more compact conformations, with the averages shifting towards lower distance values and the distributions populating compact conformations more prominently. This is especially true for the $R_g$ distribution, which in the uniform ensemble shows values between 20-60 Å with an average of 33.38±0.07 Å, while in the reweighted distribution there is clear preference for more compact structures, denoted by a clear peak at around ~25 Å, leading to a reduction in the average value to 29.07±0.11 Å. A similar change occurred in a different study, where resorting to an ensemble with more compact conformations (average $R_g$ of 27.66 Å) produced a better fit to Gal-3 experimental SAXS data (which indicated an $R_g$ value of 28.92 Å) (Lin *et al.* 2017). Additionally, there is an agreement between results obtained here for the uniform Gal-3 ensemble and the $R_g$ distribution obtained in a study where an ensemble of Gal-3 was generated using coarse-grained molecular dynamics simulations using force fields rescaled for flexible protein interactions (Thomasen *et al.* 2024). In this study, the calculated $R_g$ distribution showed values between 20-55 Å, with an average around ~ 35 Å. This shows that the presented sampling protocol can generate, without specific parametrization, results that are in line with other approaches optimized for disordered regions.

The distance matrices calculated from the Gal-3 uniform and reweighted ensembles are shown in Supplementary Fig. S18D. The distance matrix for the uniform ensemble clearly represents the long disordered tail of Gal-3 (NT) followed by its folded domain (CRD). After reweighting, conformations where the NT is closer to the CRD are preferred, with a reduction of ~10 Å in the average distance between them being observed. This suggests that the Gal-3 SAXS data is better explained by more compact conformations, where the NT tail is generally closer to the CRD domain, which could be due to a structural tendency for NT-CRD interactions. In fact, the monomeric behaviour of Gal-3 in the absence of ligands is usually tied to NT-CRD interactions, which deter Gal-3 oligomerization (Lin *et al.* 2017).

The secondary structure assignment frequency plots calculated from the Gal-3 uniform and reweighted ensembles are shown in Supplementary Fig. S18E. They clearly indicate the presence of the NT disordered region followed by the folded CRD domain, whose structure is fully conserved among all conformers. The NT region seems to show some propensity to form α-helices, especially in its N-terminal region, where around 30% of conformers are assigned this secondary structure. This is consistent with a structural study of Gal-3 that showed, using NMR, that the N-terminal region of the NT has a propensity to form transient α-helices, and that this region is crucial for interactions with the CRD (Ippel *et al.* 2016).

###### 3.2.3.3. Heterogeneous nuclear ribonucleoprotein A1

Heterogeneous nuclear ribonucleoprotein A1 (hnRNPA1) is an RNA-binding protein involved in alternative splicing regulation and mRNA transport. The structure of hnRNPA1 consists of two homologous folded RNA-recognition motif (RRM) domains that bind RNA and an intrinsically disordered low complexity glycine-rich domain (LCD). Given its central role for gene regulation, hnRNPA1 is implicated in many diseases, including viral infections, cancer and neurodegenerative diseases. Mutations in the LCD can lead to inclusion body myopathy and amyotrophic lateral sclerosis, two diseases characterized by solid deposits of RNA-binding proteins that lead to neurodegeneration (Martin *et al.* 2021). Under cellular stress conditions, hnRNPA1 is sequestered in cytoplasmic stress granules, which are formed via LLPS. While the disordered LCD has been shown to be both necessary and sufficient for inducing LLPS, it has been shown that electrostatic interactions between the RRM domains and the LCD are also important in mediating phase separation behaviour (Ritsch *et al.* 2022, Martin *et al.* 2021). In this work a conformational ensemble was generated for a hnRNPA1 construct with 314 residues containing a small flexible tail (residues 1-10), its two RRM domains (residues 11-89 and 105-179) connected by a small flexible linker (residues 90-104), followed by the LCD (residues 180-314).

The averaged structural properties calculated over the conformational ensemble generated for hnRNPA1, before and after BME reweighting, are summarized in Supplementary Fig. S19. The uniform hnRNPA1 ensemble was reweighted with BME using a value of 200 for the $\theta$ parameter, which improved the agreement with SAXS experimental data, as denoted by a reduction of the $\chi^2$ value from 2.63 to 1.06 (Supplementary Fig. S19A). The fitting of the theoretical SAXS curves calculated from the uniform and reweighted ensembles to experimental SAXS data clearly highlights this improvement across all values of $q$ (Supplementary Fig. S19B).

The probability distributions for the $R_g$, $D_{max}$ and $D_{ee}$ calculated from the hnRNPA1 uniform and reweighted ensembles are shown in Supplementary Fig. S19C. Comparing them between the uniform and reweighted ensembles shows that, in all three cases, the reweighted ensemble favours more compact conformations, with the averages shifting towards lower distance values and the distributions showing higher probability of adopting more compact conformations. This is evident in the $R_g$ distribution, which for the uniform ensemble shows values between 25-65 Å, with an average of 36.19±0.06 Å, while in the reweighted distribution there is a clear preference for more compact structures, as shown by the appearance of a defined peak around ~28 Å, shifting the average value to 32.00±0.10 Å. The $D_{max}$ distribution displays the same tendency, showing values between 60-220 Å in the uniform distribution, with an average of 115.87±0.22 Å, while the reweighted ensemble favours more compact conformations, as shown by the appearance of a defined peak around ~90 Å, leading to a reduction in the average value to 103.94±0.34 Å. It seems that after reweighting a group of more compact conformations was given a higher weight, while conformers presenting more extended conformations had their contribution reduced. Achieving a more detailed understanding of what differentiates these two groups could help elucidate the type of conformations hnRNPA1 tends to adopt in solution. Additionally, the average $R_g$ value obtained for the uniform ensemble agrees with the value obtained for an ensemble of hnRNPA1 generated through coarse-grained molecular dynamics simulations using force fields rescaled for flexible proteins, which showed values between 25-55 Å and an average $R_g$ of ~38 Å (Thomasen *et al.* 2024). This highlights the ability of the present work to generate, without specific parametrization, results that are on par with other approaches optimized for disordered regions.

The distance matrices calculated from the hnRNPA1 uniform and reweighted ensembles are shown in Supplementary Fig. S19D. The distance matrix for the uniform ensemble clearly represents the two folded RRM domains connected by a short flexible linker, followed by the flexible LCD. After reweighting, there is a clear preference for conformations where both the distance between the two RRM domains and the distance between each RRM and the LCD are smaller by about ~10 Å. This is in line with results obtained in structural studies of hnRNPA1, which concluded that the LCD adopts more compact conformations by interacting with the RRM domains (Ritsch *et al.* 2022, Martin *et al.* 2021, Levengood *et al.* 2024).

The secondary structure assignment frequency plots calculated from the hnRNPA1 uniform and reweighted ensembles are shown in Supplementary Fig. S19E. They clearly indicate the presence of two folded domains, conserved among all conformers, followed by a flexible region. The long disordered tail seems to show some propensity to form α-helices, especially in its N-terminal region right after the RRM domain, with around 30% of conformers being assigned this secondary structure. The same can be said for the short flexible linker that connects the two RRM domains, which also seems to adopt an α-helical conformation in around 30 % of conformers.

##### 3.2.3. Proteins with multiple chains

###### 3.2.3.1. Endolysin of *S. thermophilus* phage P7951

Phages, viruses that infect bacteria, are often under high selective pressure to reduce their genome and become more efficient in packing their genetic information. This can lead to the formation of genes with two in-frame overlapping open-reading frames (ORFs), resulting from the presence of an in-frame internal translation start site (iTSS). In other words, these genes can be thought of as having two alternative start codons, each with its own ribosome binding site. Such a gene can be expressed in two isoforms sharing some of their amino acid sequence: one full-length polypeptide encoded by the longest ORF and a shorter polypeptide, initiated at the iTSS, corresponding to a C-terminal part of the full-length product (Pinto *et al.* 2022).

Endolysins are the phage enzymes that cleave the host bacterial cell wall peptidoglycan to release the virion progeny. Some endolysins are encoded in genes that present an iTSS, resulting in two in-frame overlapping ORFs. The gene of endolysin LysP7951 of *S. thermophilus* phage P7951 (LysP7951) presents an iTSS, resulting in the formation of two polypeptides corresponding to a full-length product (FLP) and a C-terminal subproduct (CTP). The FLP contains both a catalytic domain at its N-terminal, responsible for peptidoglycan cleavage, and a cell-wall binding domain at its C-terminal, which corresponds to the CTP. Subunits of FLP and CTP have been shown to interact and form a multimeric complex whose presence is required for lytic activity against *S. thermophilus* (Pinto *et al.* 2022). However, the molecular mechanisms related with this oligomerization remain unsolved. A study on the multimeric organization of LysP7951 used gel filtration chromatography and mass spectrometry to conclude that LysP7951 presented a 1FLP:5CTP stoichiometry, suggesting that 6-7 CTP subunits were present in the complex. Assuming a homohexamer state, SAXS-driven *ab initio* reconstruction suggested a disklike shape with a slight central depression and protrusion on the opposite side, with an overall silhouette resembling a mushroom (Pinto *et al.* 2022).

An AF-2 prediction of a homohexamer construct of the LysP7951 CTP shows, for each protein chain, a folded domain (residues 1-56) connected by a short linker to a helical region with high confidence (residues 61-79), followed by a low confidence unstructured C-terminal region (see Supplementary Fig. S10D). Each folded domain is shown to be in contact with two neighbouring chains, forming a disklike shape similar to the one suggested by SAXS (Pinto *et al.* 2022). In this work, this AF-2 prediction was used to generate two ensembles. In both cases the interchain protein interfaces were conserved during sampling, in an effort to maintain the aforementioned disklike shape. Additionally, the helical regions were also conserved during sampling, as they were assigned high confidence values by AF and were capable of forming a similar structural protrusion to the one suggested by SAXS data (Pinto *et al.* 2022). The higher level of complexity associated with the LysP7951 system when compared to other tested proteins motivated the generation of two ensembles, each exploring a different region of the conformational space. The first ensemble, $LysP7951_{Open}$, assigned as sampling target regions both the short linker connecting the folded domain to the helical region (residues 57-60) and the low confidence C-terminal region (residues 80-102). The second ensemble, $LysP7951_{Closed}$, assigned as sampling target regions only the low confidence C-terminal region (residues 80-102). Essentially, in one ensemble the helical regions were allowed to move from their starting position in the AF-2 prediction, while in the other one they were kept in place. The results obtained for each individual ensemble were unsatisfactory: their initial agreement with experimental data was poor ($\chi^2$ > 4) and applying BME reweighting to reach a decent level of agreement with experimental data ($\chi^2$ < 2) lead to very low values for the $\phi_{eff}$ (0.36 and 0.19). This suggested that each ensemble contained a low portion of structures that were important for explaining experimental SAXS data of this construct, which results in a higher risk of overfitting. In order to arrive at a better representation of the conformational space of the protein, the two ensembles were combined into one larger ensemble, now referred to as LysP7951, increasing the pool of conformations available for reweighting.

The averaged structural properties calculated over the conformational ensemble generated for LysP7951, before and after BME reweighting, are summarized in Supplementary Fig. S20. The uniform LysP7951 ensemble was reweighted with BME using a value of 400 for the $\theta$ parameter, which improved the agreement with SAXS experimental data, as denoted by a reduction of the $\chi^2$ value from 4.22 to 1.82 (Supplementary Fig. S20A). The fitting of the theoretical SAXS curves calculated from the uniform and reweighted ensembles to experimental SAXS data highlights this improvement, especially for low values of $q$ (Supplementary Fig. S20B).

The probability distributions for the $R_g$, $D_{max}$ and $D_{ee}$ calculated from the LysP7951 uniform and reweighted ensembles are shown in Supplementary Fig. S20C. Comparing them between the uniform and reweighted ensembles shows that the reweighted ensemble favours more compact conformations, with the averages shifting towards lower distance values and the distributions showing higher probability of adoption of more compact conformations. For the uniform ensemble, the $D_{max}$ distribution showed values between 80-170 Å, with a maximum around 100 Å and an average value of 107.40±0.10 Å. After reweighting, the distribution presented values between 80-150 Å, with the maximum shifting to ~95 Å and the average value reduced to 103.02±0.17 Å, closer to the one determined for LysP7951 from experimental SAXS data (101±5 Å) (Pinto *et al.* 2022). The $R_g$ distribution for the uniform ensemble showed values between 28-36 Å, with a peak around 30.5 Å and an average value of 31.05±0.01 Å. After reweighting, the distribution presented values between 28-34 Å, with the peak shifting to ~29.5 Å and the reduced average value of 30.11±0.01 Å. Interestingly, reweighting lead to an average $R_g$ value that was now further away from one determined for LysP7951 from experimental SAXS data (30.75±0.20) (Pinto *et al.* 2022), which could indicate insufficient sampling (even after the combination of two different ensembles), possibly due to the lack of representation of the putative heptamer state in our generated ensembles.

The distance matrices calculated from the LysP7951 uniform and reweighted ensembles are shown in Supplementary Fig. S20D. The distance matrix for the uniform ensemble shows the division between different protein chains, clearly differentiating between the intra and interchain distances. Looking at the intrachain distances individually, it is clear that each monomer presents a folded domain followed by a more flexible region. Interchain distances show that folded domains of monomers that are neighbours in the LysP7951 hexamer construct are closer to each other, with the inter-domain distances increasing as they become further apart in the multichain sequence. Neighbouring helical regions, as well as the C-terminal disordered tails, also appear closer to each other than ones belonging to far away monomers. After reweighting, conformations where the disordered tails are closer to their respective folded domains by ~4 Å are preferred. Moreover, the average distance between both helical regions and the disordered tails of different chains also decreases by ~4 Å. These changes seem to support structures that more closely resemble the "mushroom" like topology proposed for LysP7951 in an earlier study (Pinto *et al.* 2022).

The secondary structure assignment frequency plots calculated from the LysP7951 uniform and reweighted ensembles are shown in Supplementary Fig. S20E. They clearly indicate, for each chain, the presence of a folded domain conserved among all conformers, followed by a α-helical region. Separating the two is a small region that is essentially always assigned a random coil conformation, and at the C-terminal the same can be observed, with α-helical conformations being assigned to this region only in ~5% of conformers.

It is clear that for LysP7951, despite using a "mixed" ensemble, it was not possible to arrive at a conformational ensemble that explained the experimental SAXS data as well as what was observed for other proteins, possibly due to the high level of complexity of the system. This means that it could prove fruitful to simply further increase the number of generated conformers, as a way to match the increased number of DOF associated with this multichain system, or explore different sampling strategies, such as varying the assignment of target regions for conformational sampling. Adding another ensemble into the mix, namely one that would represent the putative heptameric state of LysP7951, could also contribute to a better agreement with experimental SAXS data. SAXS is an ensemble method that encapsulates all the conformational states of a protein in solution, and even if the heptameric state is not present in the generated ensembles, it could be present in the experimental data.

#### 3.3. AlphaFold-guided sampling can improve ensemble accuracy and enable the study of specific structural states

The goal here was twofold: to determine if the information conveyed by AlphaFold's confidence metrics (Guo *et al.* 2022, Brotzakis *et al.* 2025) could be extracted and used to guide conformational sampling, and to understand if doing so would lead to a conformational ensemble that better aligned with SAXS experimental data.

To achieve this, the approach described in Supplementary Information S1.2.5 was applied to multidomain proteins whose AlphaFold prediction had a PAE matrix with low error values for residue pairs whose members belonged to different domains. By comparing the ensembles generated using this approach to those generated using user-defined models of these proteins, conclusions were drawn regarding the effectiveness of using AlphaFold confidence metrics to explore protein conformational space.

AlphaFold predictions were generated for two proteins: the β-galactose binding protein Galectin-3 (Gal-3) and the "mothers against decapentaplegic homolog 4" (SMAD4) transcription factor protein. Ensembles were generated for each protein using their predicted structures, incorporating information derived both from pLDDT and PAE values to bias conformational sampling. These results were compared to those obtained from ensembles generated using user-defined models of these proteins, described in Supplementary Information S3.1.3.2 (Gal-3) and Supplementary Information S3.1.2.4 (SMAD4).

##### 3.3.1. Galectin-3

Galectin-3 (Gal-3) is a β-galactose binding protein that plays a central role in many biological processes (see Supplementary Information S3.1.3.2). To generate ensembles of Gal-3, either a user-defined model (Thomasen *et al.* 2024) or an AF-2 prediction (Jumper *et al.* 2021) were used (Supplementary Fig. S21A).

The AF-2 prediction marks the disordered tail of Gal-3 (residues 1-113) as a low confidence region (pLDDT < 70), and places its N-terminal region (residues 1-30) near the folded CRD domain (residues 117-250). The accompanying PAE matrix shows low values between the N-terminal region and the CRD domain, indicating a high level of confidence in their relative position in the predicted structure. This is consistent with the fact that the N-terminal region of Gal-3 is known to interact with the CRD domain, and that this interaction is crucial for the protein's function (Lin *et al.* 2017).

Using the AF-2 prediction of Gal-3, two ensembles were generated. Both used low pLDDT values to determine target sampling regions and the PAE matrix to create inter-residue energy restraints that guided conformational sampling. One of the ensembles included these restraints without any further scaling ($Gal3_{\{pLDDT+PAE\}x1}$). The other made some of the energy restraints more permissive by applying a scaling factor of 30 to the PAE values ($Gal3_{\{pLDDT+PAE\}x30}$), using the method described in Supplementary Information S1.2.5 (Supplementary Fig. S21A).

The agreement between the three Gal-3 ensembles and experimental SAXS data was evaluated using the reduced $\chi^2$ metric, before and after BME reweighting (Supplementary Fig. S21B). The uniform Gal3 ensemble initially showed poor agreement with SAXS data ($\chi^2$ = 20.16). After reweighting, the fitting significantly improved, reducing the $\chi^2$ value to 1.5. Introducing AF-2 information into the sampling process improved the initial agreement with SAXS data, as shown by the uniform $Gal3_{\{pLDDT+PAE\}x1}$ ensemble performing better than the uniform Gal3 ensemble, presenting a $\chi^2$ value of 11.95. However, after reweighting, the $\chi^2$

value was reduced to only 2.45, which while still an improvement, did not achieve an equal level of agreement with experimental data as the one observed for the ensemble generated without using AF-2 information. Such behaviour seemed to be a consequence of the PAE-based energy restraints, which led to an ensemble that was too compact (see Supplementary Fig. S22). This motivated the hypothesis that maybe the PAE-based energy restraints were too restrictive, so a scaling factor of 30 was applied to the PAE values, aiming to loosen the contribution from AF-2 information. As a result, the uniform $Gal3_{\{pLDDT+PAE\}x30}$ ensemble showed the best agreement with experimental data out of all the uniform Gal-3 ensembles ($\chi^2$ = 4.96). After reweighting the fitting improved, lowering the $\chi^2$ value to 1.32, resulting in the best fit to experimental data out of all the Gal-3 ensembles, reweighted or otherwise. Moreover, the reweighted $Gal3_{\{pLDDT+PAE\}x30}$ ensemble retained the most information from its uniform ensemble, as shown by its $\phi_{eff}$ value of 0.84, significantly higher than the $\phi_{eff}$ value of 0.63 for the next best ensemble, Gal3.

Structural metrics probability density distributions for the $R_g$, $D_{ee}$ and $D_{max}$ were calculated for each ensemble, before and after BME reweighting (Supplementary Fig. S22). The uniform Gal3 ensemble presented the most extended conformations, with $R_g$ values spanning the 20-60 Å range and an average of 33.38±0.07 Å. After reweighting, more compact conformations were preferred, with the average $R_g$ dropping to 29.07±0.11 Å. In contrast, the uniform $Gal3_{\{pLDDT+PAE\}x1}$ ensemble showed a much more compact distribution, with $R_g$ values spanning the 20-38 Å range and an average of 24.19±0.02 Å, showing that using AF-2 information confines the conformational space. These differences can be attributed to the inter-residue energy restraints that were added between the CRD domain and the N-terminal region of Gal-3, as they presented low PAE values between them. Considering that they were close to each other in the predicted structure, the PAE-based energy restraints expectedly lead to the generation of a more compact ensemble. Unlike what was observed for Gal3, after reweighting there was a shift towards more extended conformations, with the average $R_g$ increasing to 26.62±0.08 Å. Both the Gal3 and $Gal3_{\{pLDDT+PAE\}x1}$ ensembles showed poor agreement with experimental SAXS data prior to reweighting, but for different reasons. For Gal3, the initial ensemble populated conformations that were too extended, while for $Gal3_{\{pLDDT+PAE\}x1}$ the initial ensemble was too compact. Applying BME reweighting led to both ensembles becoming closer to the experimental data, but while for Gal3 this involved a preference for more compact conformations, for $Gal3_{\{pLDDT+PAE\}x1}$ a shift towards more extended conformations was required.

The uniform $Gal3_{\{pLDDT+PAE\}x30}$ ensemble exhibited a distribution similar to the one observed for $Gal3_{\{pLDDT+PAE\}x1}$, albeit slightly less compact, with $R_g$ values spanning the 20-45 Å range and an average of 26.04±0.04 Å. After reweighting no major changes to the distribution were observed, with only a slight preference for more extended conformations, increasing the average $R_g$ to 27.96±0.08 Å. The introduction of a scaling factor for the PAE-based energy restraints in the $Gal3_{\{pLDDT+PAE\}x30}$ ensemble allowed for the sampling of slightly more extended conformations, while still maintaining a relatively compact ensemble. This helped in overcoming the excessive level of compactness observed in the $Gal3_{\{pLDDT+PAE\}x1}$ ensemble (where no scaling was applied), resulting in a much better fit to experimental SAXS data. This improvement is reflected in the $\chi^2$ values for each ensemble prior to reweighting (Gal3= 20.16, $Gal3_{\{pLDDT+PAE\}x1}$= 11.95, $Gal3_{\{pLDDT+PAE\}x30}$= 4.96).

These results highlight the $Gal3_{\{pLDDT+PAE\}x30}$ ensemble as the one that best explains the SAXS experimental data and thus the best candidate to represent the conformational variability of Gal-3 in solution. The introduction of a scaling factor for the PAE-based energy restraints proved critical for the generation of a more accurate ensemble. Considering the improvement observed with a scaling factor of 30, it is possible that a different value could lead to an even better fit to experimental data. A screening of scaling factor values could lead to an ensemble with the optimal fit to experimental data, where ideally we could retain close to 100% of the information contained in the initial ensemble, *i.e.* a $\phi_{\mathrm{eff}}$ value close to 1, but this was not pursued here. Nevertheless, these results highlight the possibility for future users to explore different scaling factors applied to their protein of interest in order to enhance their ensemble's alignment with SAXS data.

##### 3.3.2. "Mothers against decapentaplegic homolog 4"

"Mothers against decapentaplegic homolog 4" (SMAD4) is a transcription factor protein that plays a central role in the transforming growth factor-β (TGF-β) signalling pathway (see Supplementary Information S3.1.2.4). To generate ensembles of SMAD4, either a user-defined model (Gomes *et al.* 2021) or an AF-3 prediction (Abramson *et al.* 2024) were used (Supplementary Fig. S23A).

The AF-3 prediction assigns low pLDDT values (< 70) to the inter-domain disordered linker (residues 140-314), and places the MH1 (residues 10-141) and MH2 (residues 314-554) folded domains in close proximity. Interestingly, the SAD domain (residues 272-314), whose folded state is known to be important for SMAD4 function (Gomes *et al.* 2021), is included in the low confidence region, and is not assigned its characteristic secondary structure (Qin, Lam and Lin 1999). While most of the disordered linker is shown as a flexible ribbon around the two folded domains, a specific segment (residues 160~200) is assigned a β-strand secondary structure and placed close to the MH2 domain, seemingly contributing to β-sheet formation. The accompanying PAE matrix shows low values between this region of the disordered linker (residues 160~200) and the MH2 domain, indicating a high level of confidence in their relative position in the predicted structure.

Using the AF-3 prediction of SMAD4, two ensembles were generated, both relying on low pLDDT scores to define target sampling regions. In one ensemble, only the pLDDT was considered ($\mathrm{SMAD4_{pLDDT}}$). In the other ensemble, both pLDDT and PAE were used, with the PAE matrix being translated into inter-residue energy restraints that guided conformational sampling ($\mathrm{SMAD4_{pLDDT+PAE}}$). The strength of these restraints was reduced by a factor of 20, using the approach described in Supplementary Information S1.2.5.

The agreement between the three SMAD4 ensembles and experimental SAXS data was evaluated using the reduced $\chi^2$ metric, before and after BME reweighting (Supplementary Fig. S23B). The uniform SMAD4 ensemble initially showed a relatively poor agreement with experimental SAXS data ($\chi^2$ = 4.37). After reweighting the fitting significantly improved, reducing the $\chi^2$ value to 1.15. The uniform $\mathrm{SMAD4_{pLDDT}}$ ensemble presented a significantly worse initial agreement with SAXS data ($\chi^2$ = 9.22) when compared to the SMAD4 ensemble. It appears that including the SAD domain in the target sampling regions has a negative effect on the fitting to SAXS data. This agrees with previous findings that determined that keeping the SAD domain in its folded state results in a better agreement with experimental SAXS data of SMAD4 (Gomes *et al.* 2021). Although after reweighting the fitting of the $\mathrm{SMAD4_{pLDDT}}$ improved significantly ($\chi^2$ = 1.25), it still did not reach the level of agreement observed for the SMAD4 ensemble. Moreover, to achieve this improvement less information was retained from the initial $\mathrm{SMAD4_{pLDDT}}$ ensemble when compared to the SMAD4 ensemble, as shown by their $\phi_{\mathrm{eff}}$ values (0.52 and 0.75, respectively). The uniform $\mathrm{SMAD4_{pLDDT+PAE}}$ ensemble performed similarly to the SMAD4 ensemble, with a relatively poor agreement with SAXS data ($\chi^2$ = 5.56). After reweighting the fitting managed to improve significantly ($\chi^2$ = 1.43), but still fell short of the agreement observed for the SMAD4 ensemble, and retained less information from the initial ensemble ($\phi_{eff}$ = 0.54).

Structural metrics probability density distributions for the $R_g$, $D_{ee}$ and $D_{max}$ were calculated for each ensemble, before and after BME reweighting (Supplementary Fig. S24). The uniform SMAD4$_{pLDDT}$ ensemble presented the most extended conformations, with $R_g$ values spanning the 30-120 Å range and an average of 56.65±0.14 Å. Although the average $R_g$ value did not change significantly after reweighting (55.66±0.47 Å), there was a clear preference for more compact structures, denoted by the appearance of a maximum around 40 Å. The uniform SMAD4 ensemble showed a slightly more compact distribution, with $R_g$ values spanning the 30-100 Å range and an average of 50.05±0.11 Å. This is expected, as for this ensemble the SAD domain was kept in its folded state while for the SMAD4$_{pLDDT}$ ensemble this region was initialized in an unfolded state and was later targeted for conformational sampling. The SAD domain in its folded state is naturally more compact than in its unfolded state, and that might contribute to the difference observed in the average $R_g$ values of each ensemble. Similar to the SMAD4$_{pLDDT}$ ensemble, after reweighting the average $R_g$ value of the SMAD4 ensemble did not change significantly (49.66±0.25 Å), but a clear peak around 40 Å appeared in the distribution. The uniform SMAD4$_{pLDDT+PAE}$ ensemble was the most compact, with $R_g$ values spanning the 30-70 Å range and an average of 44.12±0.06 Å. This is due to the inter-residue energy restraints that were added between the MH2 domain and a section of the disordered linker region, since they presented low PAE values between them. Given that they were close to each other in the predicted structure, these restraints naturally lead to the generation of a more compact ensemble. After reweighting, there was a slight preference for more extended conformations in the resulting distribution, even if the average $R_g$ did not change significantly, increasing to 44.78±0.13 Å.

Unlike what was observed for Gal-3, using AF confidence metrics to guide the conformational sampling of SMAD4 did not lead to an ensemble that better aligned with experimental SAXS data. Here, extracting information from the PAE matrix and using it to guide the conformational sampling process leads to an ensemble that only captures more compact conformations of SMAD4. Considering that SMAD4 has been shown to exist in an "open-closed" conformational equilibrium (Gomes 2019), it seems that only the "closed" state is being captured by the PAE matrix. Additionally, the $R_g$ values associated with the SMAD4$_{pLDDT+PAE}$ ensemble (around ~42 Å) are similar to the ones previously experimentally observed for "closed" conformations of SMAD4 (Gomes *et al.* 2021). Thus, any ensemble generated using this information would be inherently biased towards this structural state. Assuming the information conveyed by the PAE matrix is correct, this approach could prove useful for studying the "closed" state of SMAD4, but not for exploring its full conformational variability. Similar to what was attempted for the LysP7951 protein system (see Supplementary Information S3.1.3.1), mixing the SMAD4$_{pLDDT+PAE}$ ensemble with more open conformations of SMAD4 could lead to a more accurate representation of the conformational variability of this protein and thus improve the agreement with experimental SAXS data.

### Supplementary Tables

**Table S1. Database memory size optimization for the *all* dataset.** Column name, size in the original database, size in the optimized database, difference (in percentage). All sizes are in Megabytes (MB) and were calculated using the Pandas *memory_usage* method with argument *deep=True*.

| Column | Size (MB) | | Difference (%) |
|---|---|---|---|
| | Original | Optimized | |
| Fragment | 385.69 | 13.57 | - 96.5 |
| φ Residue 1 | 51.43 | 25.71 | - 51 |
| ψ Residue 1 | | | |
| ω Residue 1 | | | |
| φ Residue 2 | | | |
| ψ Residue 2 | | | |
| ω Residue 2 | | | |
| φ Residue 3 | | | |
| ψ Residue 3 | | | |
| ω Residue 3 | | | |
| Total | 848.52 | 244.98 | - 79 |

**Table S2. pLDDT + PAE restraint strength scaling scenarios for a residue pair (X, Y).** The pLDDT value of residue X, the pLDDT value of residue Y, the PAE value between (X, Y) and a measure of the "strength" of the energy restraint between them, corresponding to σ in Supplementary Equation S2.

| Scenario | $pLDDT_X$ | $pLDDT_Y$ | $PAE_{\{X,Y\}}$ | Restraint "Strength" |
|---|---|---|---|---|
| 1 | High | High | Low | $PAE_{\{X,Y\}}$ |
| 2 | High | Low | Low | $PAE_{\{X,Y\}} \times \gamma$ |
| 3 | Low | Low | Low | No restraint (unlikely scenario) |
| | - | - | High | No restraint (discarded by PAE cutoff) |

**Table S3. Tested protein structural architectures.** Protein System, Number of amino acid residues ($N_r$), sampled residue regions, time to generate 10000 conformers, reference for the used initial structure and reference for the SAXS experimental data. All ensembles were generated on a machine running Ubuntu 22.04.4 LTS, with a 13th Gen Intel® Core™ i9-13900KF processor (32 cores) and 64 GB of RAM.

| Protein System | $N_r$ | Sampled regions | Time | Structure Reference | SAXS Reference |
|---|---|---|---|---|---|
| **Hst5** | 24 | 1-24 | 5 min | (Thomasen *et al.* 2024) | (Jephthah *et al.* 2019) |
| **aSyn** | 140 | 1-140 | 24 min | | (Ahmed *et al.* 2021) |
| **cMyBP-C$_{mTHB-C2}$** | 137 | 1-5, 43-49 | 30 min | | (Michie *et al.* 2016) |
| **Ubq$_4$** | 304 | 73-76, 149-152, 225-228, 301-304 | 4h 10 min | | (Jussupow *et al.* 2020) |
| **TIA1** | 275 | 1-5, 83-94, 173-189 | 1h 40 min | | (Sonntag *et al.* 2017) |
| **Gal-3** | 250 | 1-116 | 1h 45 min | | (Lin *et al.* 2017) |
| **hnRNPA1** | 314 | 1-10, 90-104, 180-314 | 2 h | | (Martin *et al.* 2021) |
| **SMAD4** | 554 | 140-271 | 1d 2h 40 min | (Gomes *et al.* 2021) | (Gomes *et al.* 2021) |
| **USH3** | 152 | 1-88, 146-152 | 30 min | (Arbesú *et al.* 2017) | (Arbesú *et al.* 2017) |
| **LysP7951** | 612 (102x6) | 59-102 (x6) | 2d 5h 22 min | (Jumper *et al.* 2021) (AF-2) | (Pinto *et al.* 2022) |

**Table S4. Tested protein structural architectures using AlphaFold-guided conformational sampling.** Protein System, Number of amino acid residues ($N_r$), sampled residue regions, time to generate 10000 conformers**,** reference for the used initial structure and reference for the SAXS experimental data. All ensembles were generated on a machine running Ubuntu 22.04.4 LTS, with a 13th Gen Intel® Core™ i9-13900KF processor (32 cores) and 64 GB of RAM.

<table>
<tr><th>Protein System</th><th>$N_r$</th><th>Sampled regions</th><th>Time</th><th>Structure Reference</th><th>SAXS Reference</th></tr>
<tr><td>SMAD4$_{\mathbf{pLDDT}}$</td><td>554</td><td rowspan="2">1-9, 140-180, 189-311, 471-490, 549-554</td><td>2d 5h 20 min</td><td rowspan="2">(Abramson et al. 2024) (AF-3)</td><td rowspan="2">(Gomes et al. 2021)</td></tr>
<tr><td>SMAD4$_{\mathbf{\{pLDDT+PAE\}x20}}$</td><td>554</td><td>9d 19h 22 min</td></tr>
<tr><td>Gal-3$_{\mathbf{\{pLDDT+PAE\}x1}}$</td><td>250</td><td>1-112</td><td>4h 22 min</td><td rowspan="2">(Jumper et al. 2021) (AF-2)</td><td rowspan="2">(Lin et al. 2017)</td></tr>
<tr><td>Gal-3$_{\mathbf{\{pLDDT+PAE\}x30}}$</td><td>250</td><td>1-112</td><td>2h 44 min</td></tr>
</table>

### Supplementary Equations

$$P(accept) = \begin{cases} \exp^{\frac{-\Delta E}{k_b T}} & \text{if } \Delta E \geq 0 \\ 1 & \text{if } \Delta E < 0 \end{cases}$$

**Supplementary Equation S1:** The Metropolis Criterion.

$$f(x) = \begin{cases} 0 & \text{if } x \in [x_0 - tol, x_0 + tol] \\ (\frac{x - x_0}{\sigma})^2 & \text{if } x \notin [x_0 - tol, x_0 + tol] \end{cases}$$

**Supplementary Equation S2:** Restraint function.

$$f(x) = \begin{cases} 0 & \text{if } x \in [x_0 - tol, x_0 + tol] \\ (\frac{x - x_0}{PAE_{\{X,Y\}} \times \gamma})^2 & \text{if } x \notin [x_0 - tol, x_0 + tol] \end{cases}$$

**Supplementary Equation S3:** Restraint function using PAE values.

$$\mathcal{L}(w_1 \dots w_n) = \frac{m}{2}\chi^2(w_1 \dots w_n) - \theta S_{\text{REL}}(w_1 \dots w_n)$$

**Supplementary Equation S4:** Function to minimize during BME reweighting.

$$\chi^2(w_1 \dots w_n) = \frac{1}{m} \sum_i^m \frac{\left(\sum_j^n w_j F_i(\mathbf{x}_j) - F_i^{\text{EXP}}\right)^2}{\sigma_i^2}$$

.

**Supplementary Equation S5:** Reduced $\chi^2$ between calculated and experimental values.

$$S_{\text{REL}} = -\sum_j^n w_j log\left(\frac{w_j}{w_j^0}\right)$$

**Supplementary Equation S6:** Relative entropy.

$$\phi_{eff} = \exp(S_{REL})$$

**Supplementary Equation S7:** Fraction of effective frames.

### Supplementary Figures

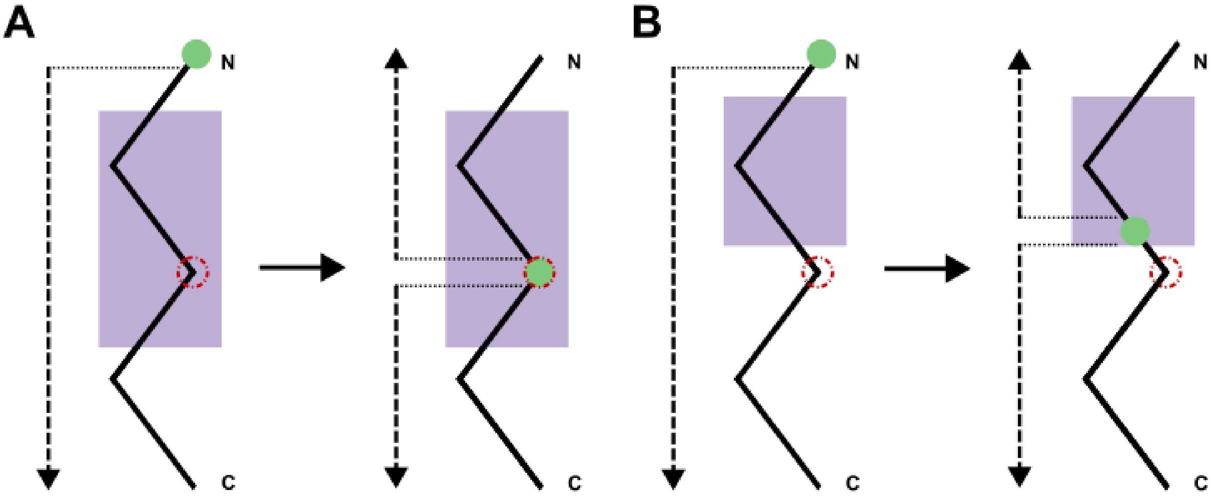

**Figure S1. Manipulation of the FoldTree morphology to minimize propagation of conformational changes.** For each case, the default FoldTree morphology is shown, followed by the optimized version. The protein chain (black line) is represented from N- to C-terminus, with conformational changes propagating through the chain from the residue assigned as the root vertex (green), in the direction indicated by the dashed arrows. The "middle" residue of the chain (red) and an example non-sampled protein region whose structure should remain conserved (lilac). **A.** If the "middle" residue of the protein chain is in a non-sampled region, that residue is assigned as the root vertex. **B.** If the "middle" residue of the protein chain is not in a non-sampled region, the closest residue to the "middle" residue that is also contained in a non-sampled region is assigned as the new root vertex.

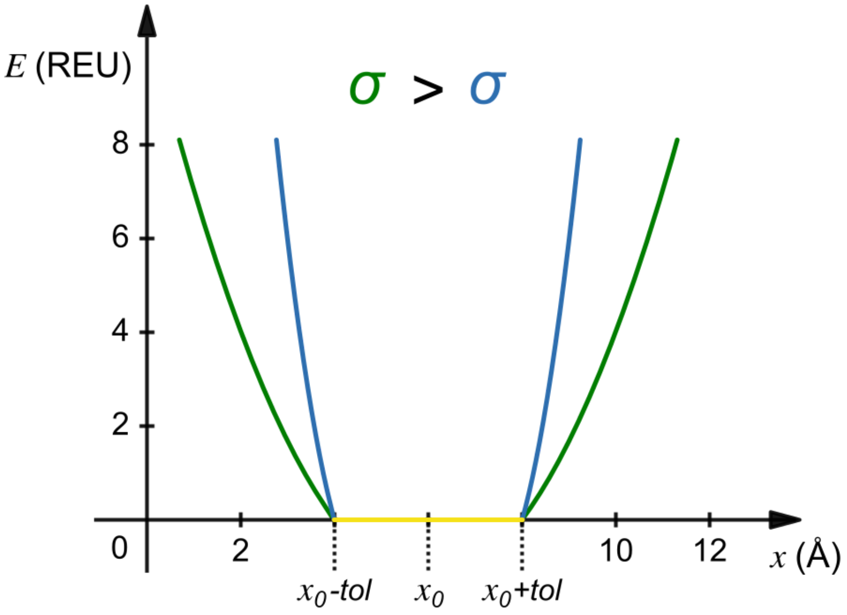

**Figure S2. Graphical representation of the restraint function.** The *x*-axis represents the inter-atomic distance between restrained atoms after a Monte Carlo move, and the *y*-axis shows the restraint score (energy, *E*) in Rosetta Energy Units (REU). In this example, the initial inter-atomic distance is 6 Å, with a tolerance (*tol*) of 2 Å. In the plot we can see how decreasing the σ parameter results in more restrictive restraints, as the energy penalty increases more sharply as we stray further from $x_0$.

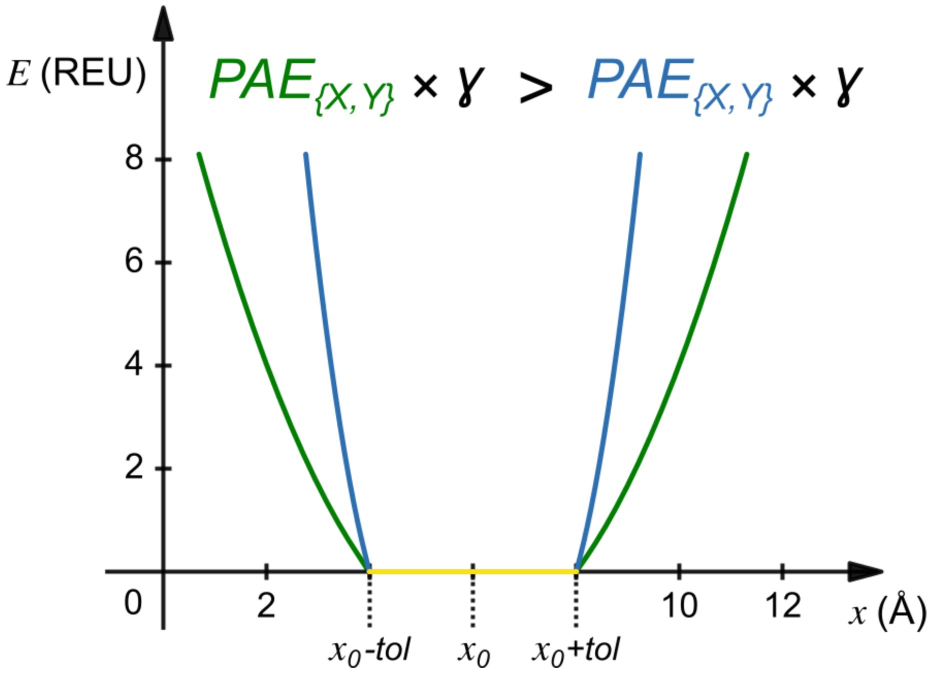

**Figure S3. Graphical representation of the restraint function using PAE values.** The *x*-axis represents the inter-atomic distance between restrained atoms after a Monte Carlo move, and the *y*-axis shows the restraint score (energy, *E*) in Rosetta Energy Units (REU). In this example, the initial inter-atomic distance is 6 Å, with a tolerance (*tol*) of 2 Å. In the plot we can see how smaller PAE values between pairs of residues result in more restrictive restraints, as the energy penalty increases more sharply as we stray further from $x_0$.

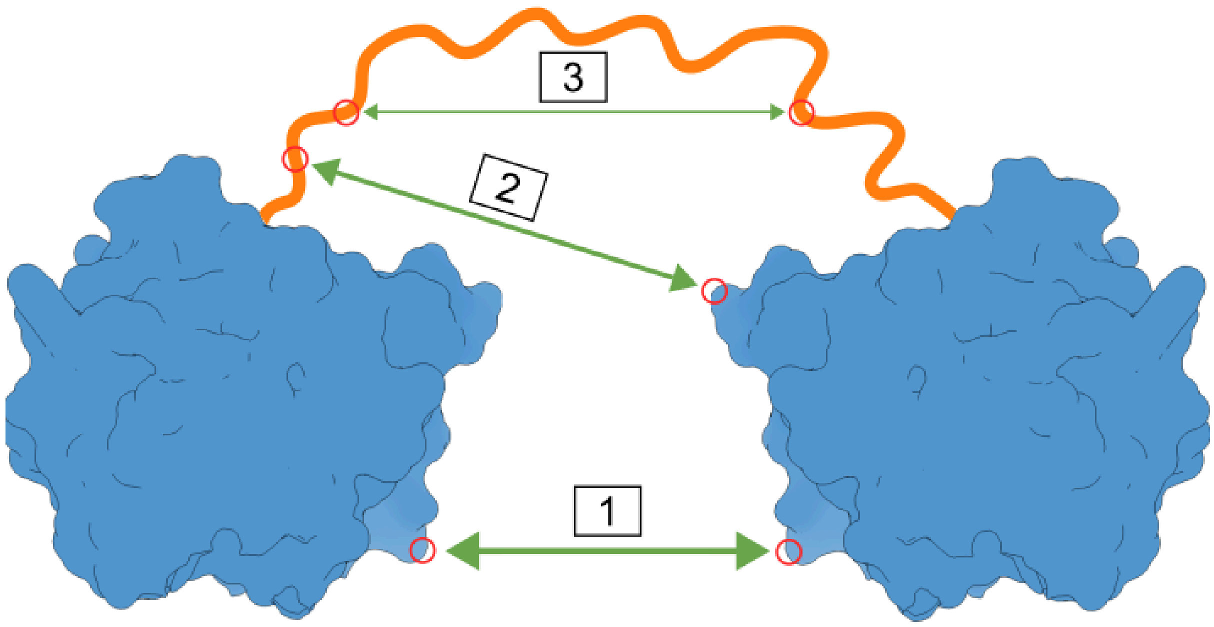

**Figure S4. pLDDT + PAE restraint strength scaling scenarios.** Example protein with two folded domains connected by a disordered linker. Numbering of scenarios follows Supplementary Table S2. The thickness of the arrows is directly proportional to restraint strength.

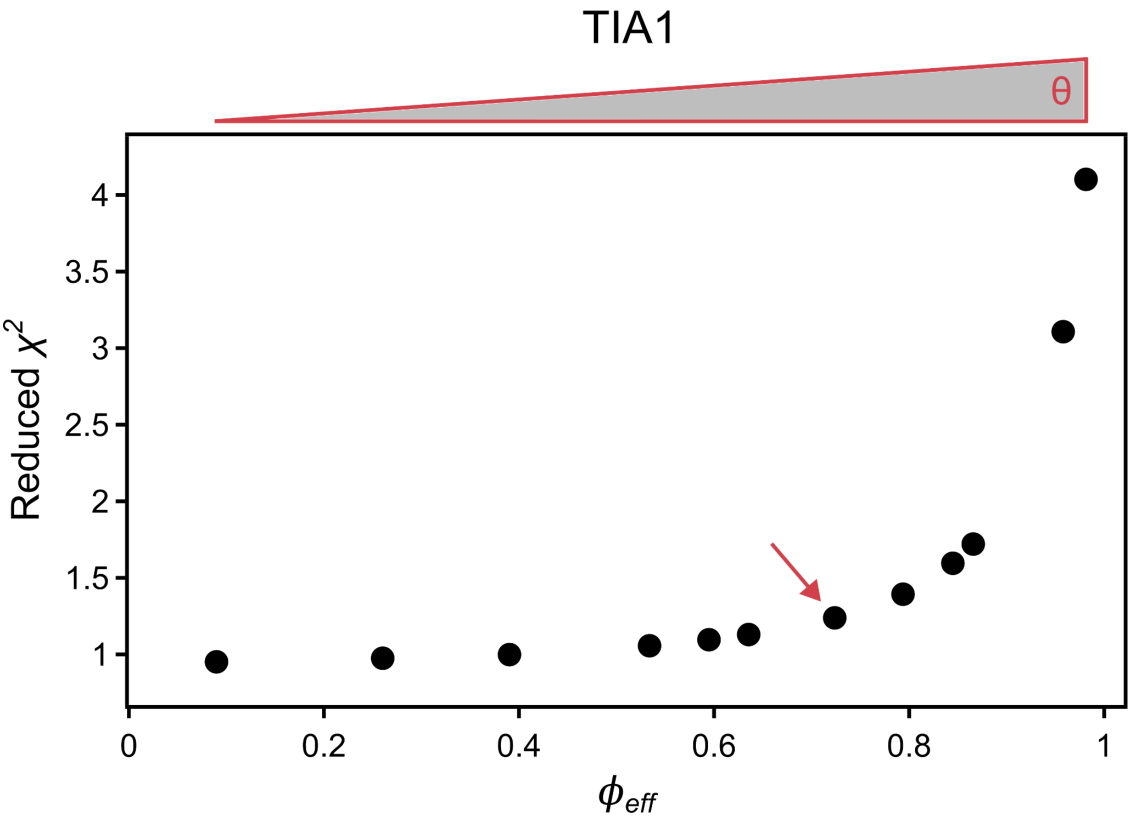

**Figure S5. Elbow plot for an ensemble of 10000 conformers of TIA1.** The plot depicts the number of effective frames ($\phi_{eff}$) vs. reduced $\chi^2$. Each data point was obtained by using a different value for the $\theta$ parameter. The tried values for the $\theta$ parameter were {1, 10, 20, 50, 75, 100, 200, 400, 750, 1000, 5000, 10000}. The optimal value for $\theta$, the "elbow" of the plot, is highlighted with a red arrow.

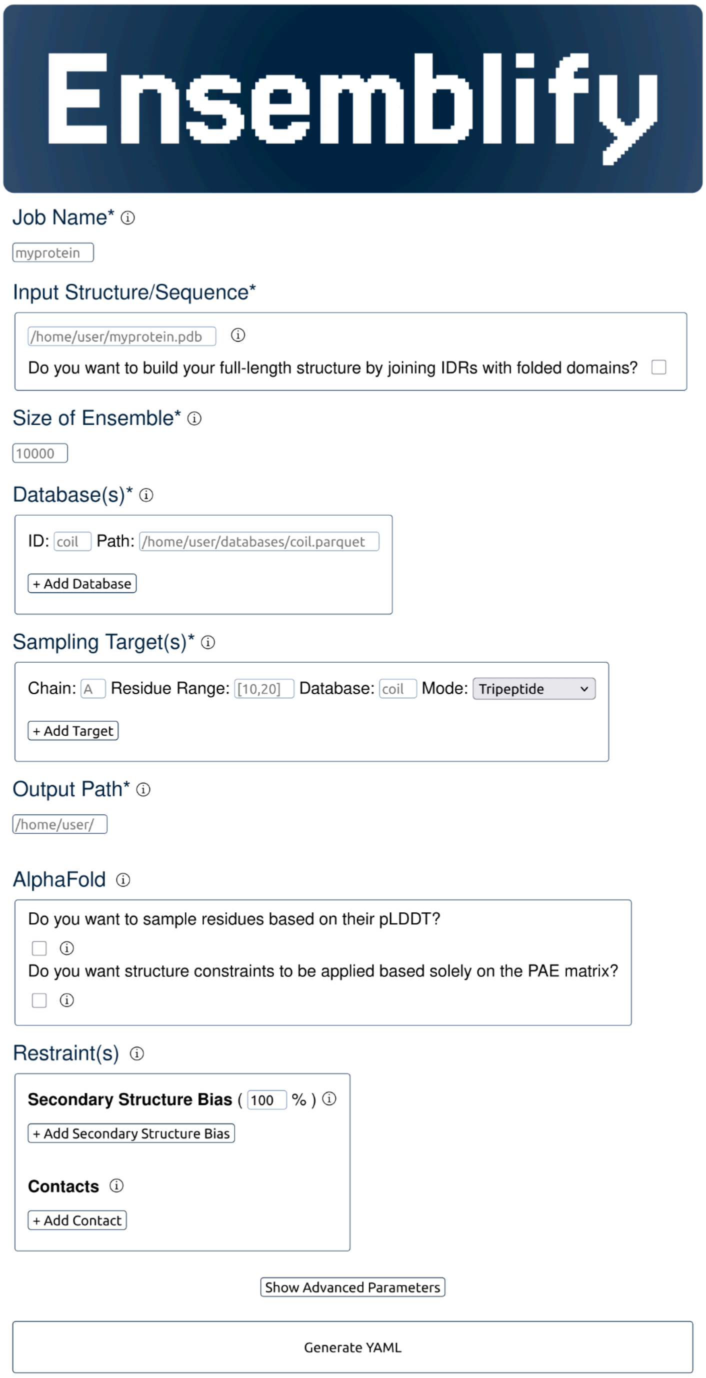

**Figure S6. Snapshot of the Ensemblify HTML parameters form.** Required fields are marked with an asterisk. Users can specify a Job Name, provide an Input Structure path or a protein Sequence (with the option to combine IDRs with folded domains to create a full-length structure), define the Size of the desired ensemble, select Dihedral Angle Databases, specify Sampling Target regions,and set the Output Path. When using an AlphaFold model as an input structure, the pLDDT can guide the definition of sampling regions, while PAE values can be used to apply energy restraints to the structure during sampling. Secondary Structure Biases can be set for specific regions of specific chains in a desired percentage of structures in the final ensemble. Regions involved in intra or inter-chain interfaces can be marked as contacts to conserve their relative positions. Additional customization is available in the Advanced Parameters section. Once the form is completed, a properly formatted YAML parameters file can be generated with a single click.

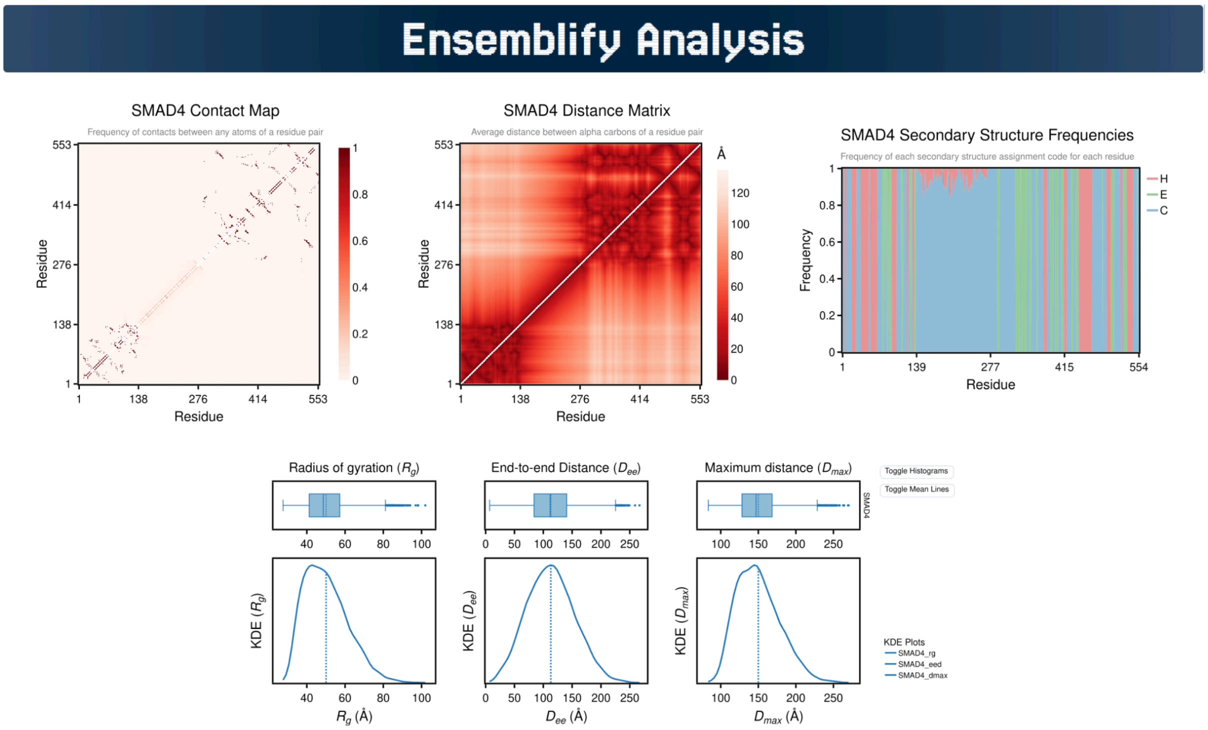

**Figure S7. Snapshot of an Ensemblify analysis dashboard for an ensemble of SMAD4.** Contains analysis figures for an ensemble of SMAD4 generated from a user-defined model. The dashboard includes a Contact Map, a Distance Matrix and a Secondary Structure Assignment Frequency Plot. Additionally, boxplots and probability density distributions for structural metrics such as the radius of gyration ($R_g$), maximum $C_\alpha$ distance ($D_{max}$) and end-to-end distance ($D_{ee}$) are also provided.

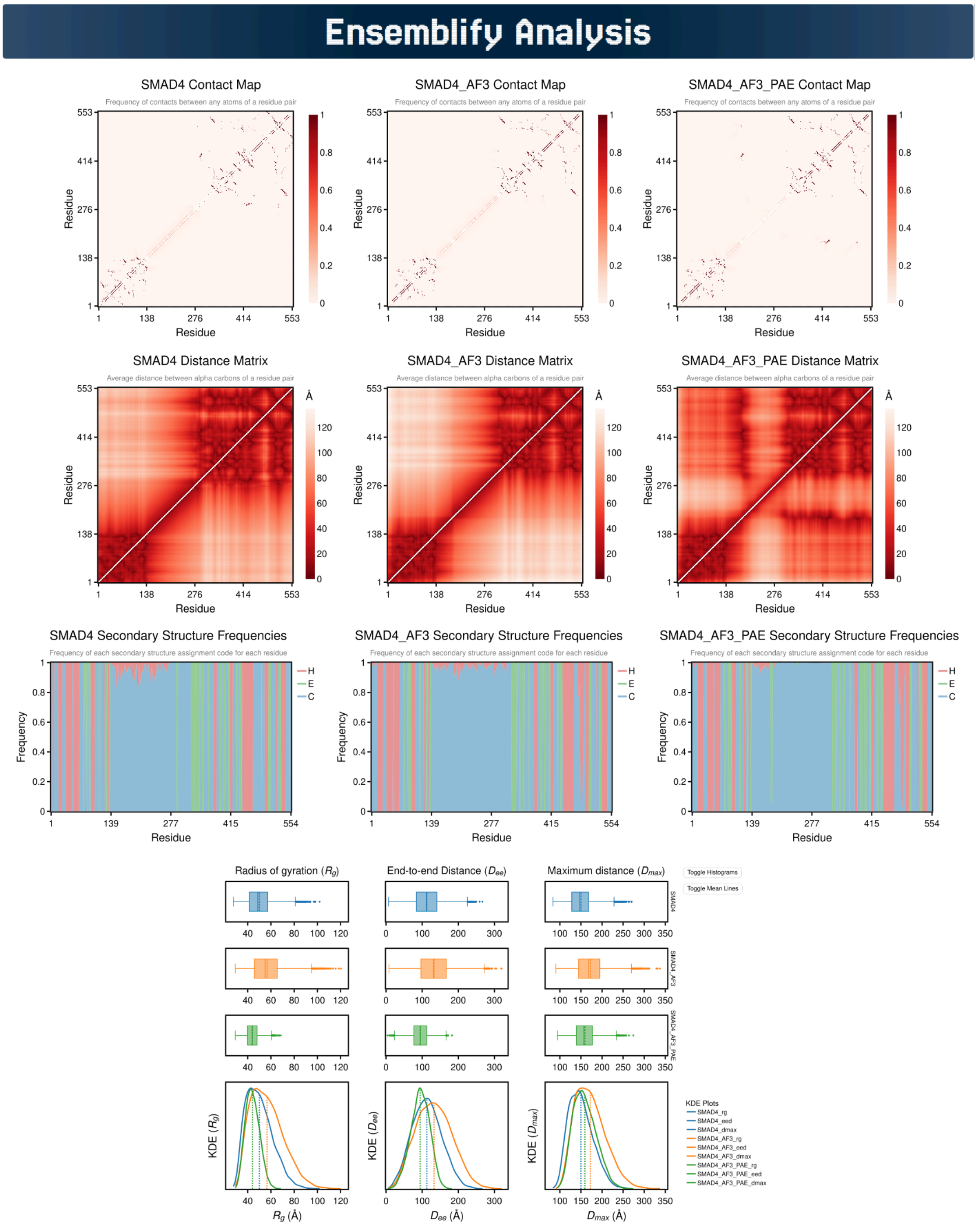

**Figure S8. Snapshot of an Ensemblify analysis dashboard comparing multiple ensembles of SMAD4.** The dashboard displays ensembles generated from: (i) a user-defined model (left), (ii) an AlphaFold3 prediction using the pLDDT to define sampling regions, and (iii) an AlphaFold3 prediction using both the pLDDT (for sampling) and PAE matrix (for restraints). For each ensemble, the dashboard includes a Contact Map, a Distance Matrix and a Secondary Structure Assignment Frequency Plot. Additionally, boxplots and probability density distributions of structural metrics such as the radius of gyration ($R_g$), maximum $C_\alpha$ distance ($D_{max}$) and end-to-end distance ($D_{ee}$) are available in a multiplot figure that facilitates ensemble comparison.

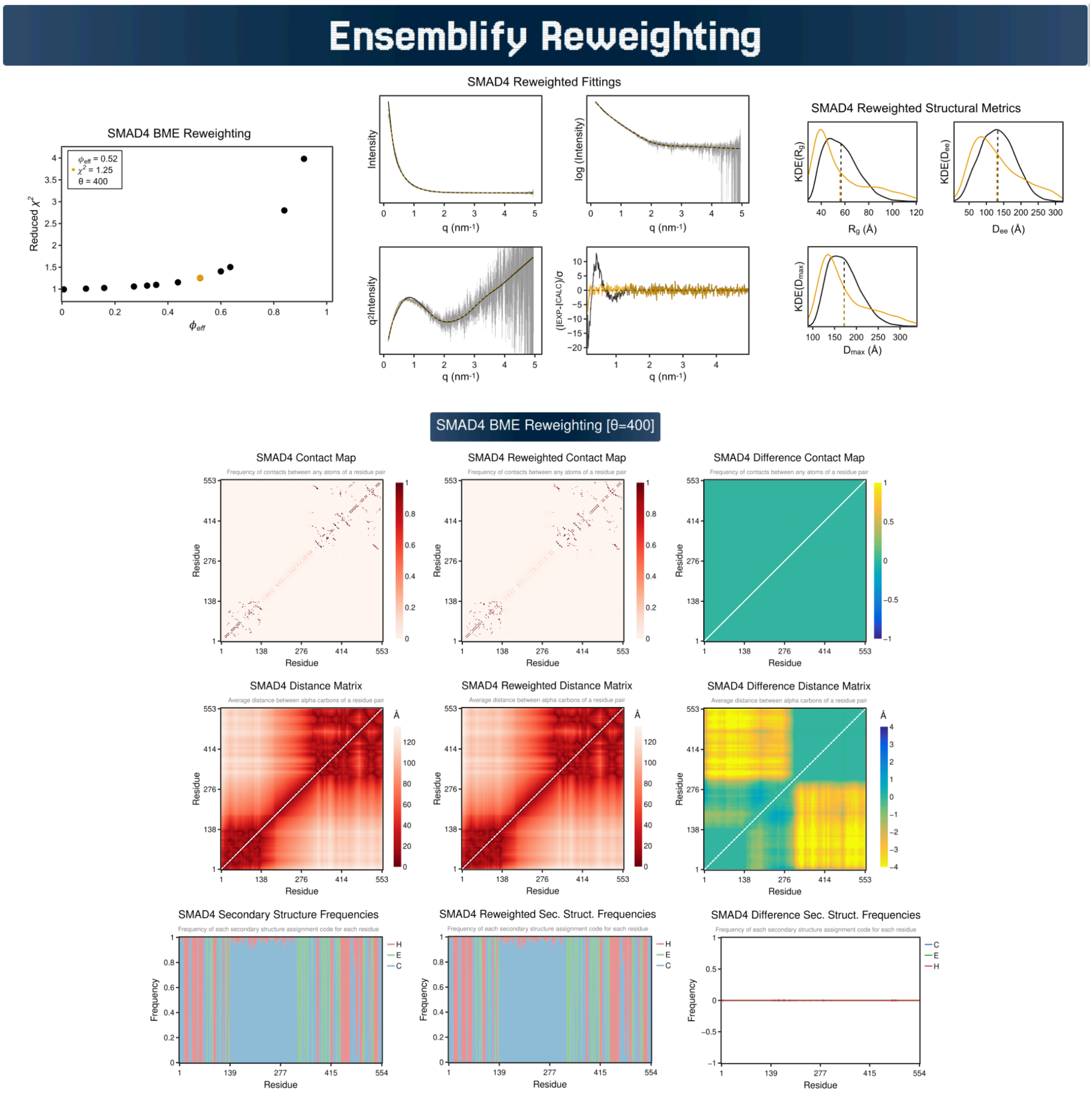

**Figure S9. Snapshot of an Ensemblify reweighting dashboard for an ensemble of SMAD4.** The dashboard was generated from an ensemble of SMAD4 that was created from a user-defined model. For each selected $\theta$ value, the corresponding weights are used to recalculate ensemble analysis metrics, allowing for comparisons between the initial uniformly weighted (uniform) ensemble and its reweighted version. The dashboard includes: (i) a $\phi_{eff}$ versus $\chi^2$ plot across $\theta$ values, (ii) SAXS profiles fittings comparing the experimental data with profiles from uniform and reweighted ensembles, and (iii) probability density distributions for different structural metrics calculated from the uniform and reweighted ensembles. To aid comparison of Contact Maps, Distance Matrices and Secondary Structure Assignment Frequency Plots between ensembles, difference plots are provided, calculated by subtracting the uniform values from the reweighted ones.

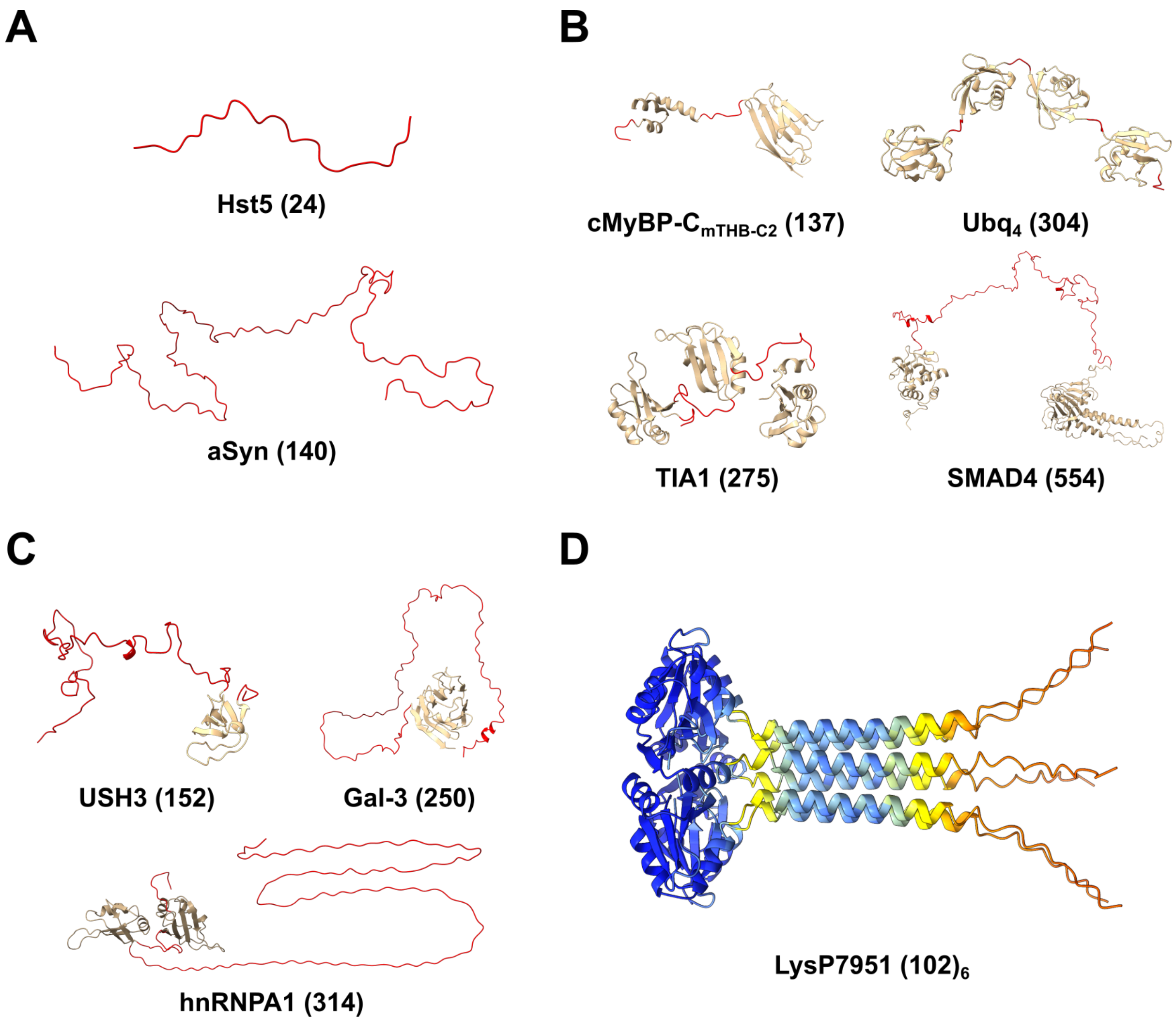

**Figure S10. Tested protein structural architectures.** For each protein,the total number of residues is shown next to its identifier, with sampled regions highlighted in red. The displayed protein structures are user-defined models, except for LysP7951, for which the AF-2 prediction is shown. **A.** Full IDPs: Histatin5 (Hst5) and α-synuclein (aSyn). **B.** Multi-domain proteins connected by flexible linkers: the tri-helix bundle of the m domain of cardiac myosin-binding protein C (cMyBP-C) connected to the C2 domain (cMyBP-C$_{mTHBC2}$), linear tetraubiquitin (Ubq$_4$), T-cell intracellular antigen-1 (TIA1), and "mothers against decapentaplegic homolog 4" (SMAD4). **C.** Folded domain(s) with a long flexible tail: the SH4, Unique and SH3 domains of non-receptor tyrosine kinase Src (USH3), Galectin-3 (Gal-3), and heterogeneous nuclear ribonucleoprotein A1 (hnRNPA1). **D.** Multi-chain protein: AF-2 prediction of a homohexamer construct of the C-terminal product subunit of the endolysin of *S. thermophilus* phage P7951 (LysP7951), coloured by pLDDT using the AF palette.

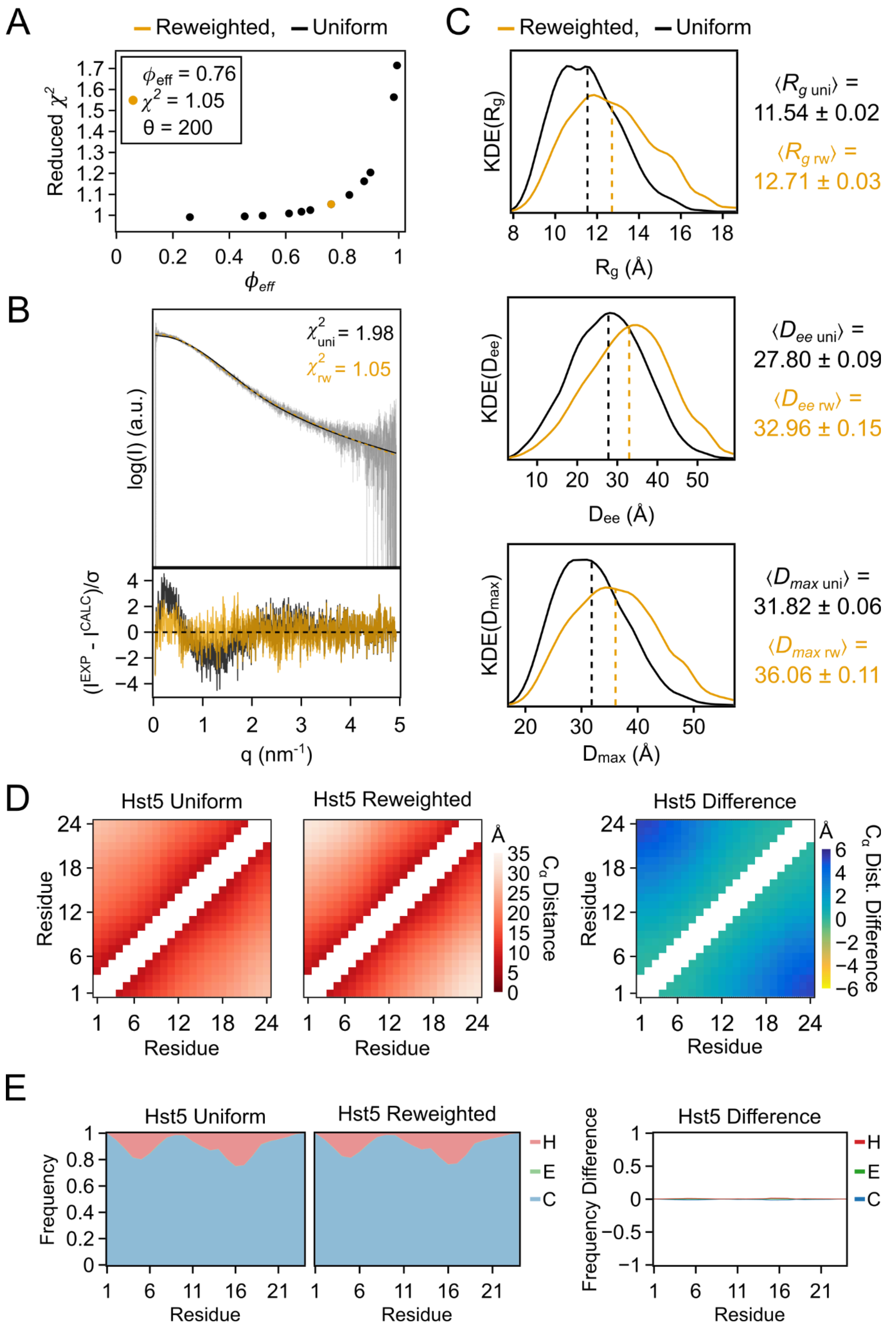
A
Reweighted, Uniform
ϕeff = 0.76
χ2 = 1.05
θ = 200
Reduced χ2
ϕeff
B
χ2uni = 1.98
χ2rw = 1.05
log(I) (a.u.)
(IEXP - ICALC)/σ
q (nm-1)
C
Reweighted, Uniform
KDE(Rg)
Rg (Å)
⟨Rg uni⟩ =
11.54 ± 0.02
⟨Rg rw⟩ =
12.71 ± 0.03
KDE(Dee)
Dee (Å)
⟨Dee uni⟩ =
27.80 ± 0.09
⟨Dee rw⟩ =
32.96 ± 0.15
KDE(Dmax)
Dmax (Å)
⟨Dmax uni⟩ =
31.82 ± 0.06
⟨Dmax rw⟩ =
36.06 ± 0.11
D
Hst5 Uniform
Hst5 Reweighted
Hst5 Difference
Residue
Cα Distance
Cα Dist. Difference
Å
E
Hst5 Uniform
Hst5 Reweighted
Hst5 Difference
Frequency
Frequency Difference
Residue
H
E
C

**Figure S11. Hst5 ensemble analysis and reweighting highlights.** Comparison of calculated structural properties for the uniformly weighted ensemble of Hst5 (“uniform”) and the ensemble after reweighting using SAXS data via BME approach (“reweighted”). Differences are calculated by subtracting the uniformly weighted values from the reweighted ones. **A.** $\phi_{\mathrm{eff}}$ versus $\chi^2$ plot across $\theta$ values , with the selected $\theta$ value highlighted in gold. **B.** SAXS profile fits from the uniform (black) and reweighted (gold) ensembles) against experimental data (top), and the respective standardized residuals (bottom). **C.** Probability density distributions of the radius of gyration ($R_g$), end-to-end distance ($D_{ee}$), and maximum $C_\alpha$-$C_\alpha$ distance ($D_{max}$) for both ensembles (black-uniform; gold-reweighted). Dashed lines indicate weighted average values for each distribution, with its values detailed on the right of the plots along with their respective standard error. **D.** $C_\alpha$-$C_\alpha$ distance matrices for the uniform and reweighted ensembles, and their difference. **E.** Secondary structure assignment frequency plots for the uniform and reweighted ensembles, and their difference.

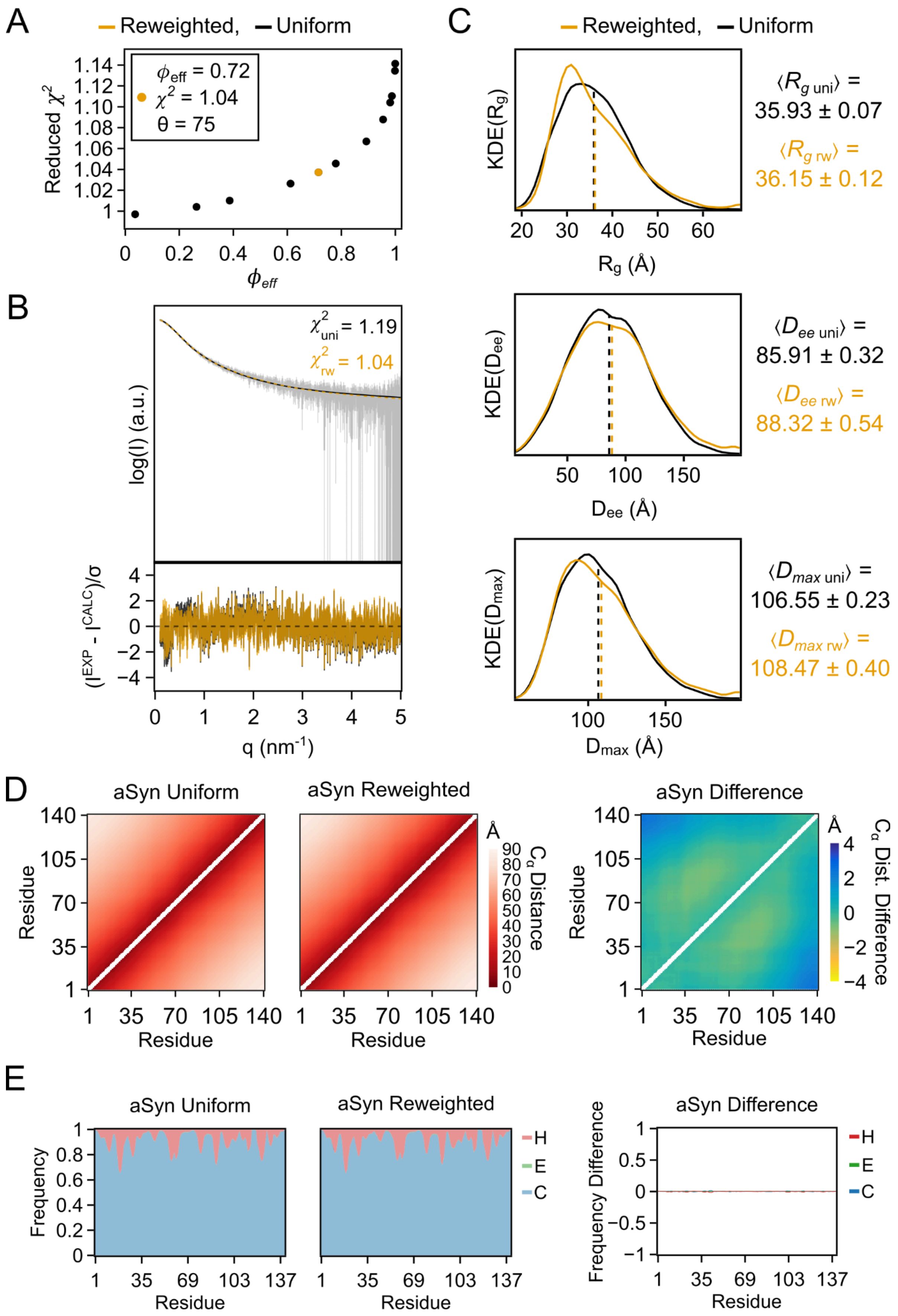

A
Reweighted, Uniform
$\phi_{eff}$ = 0.72
$\chi^2$ = 1.04
θ = 75
Reduced $\chi^2$
$\phi_{eff}$
B
$\chi^2_{uni}$ = 1.19
$\chi^2_{rw}$ = 1.04
log(I) (a.u.)
$(I^{EXP} - I^{CALC})/\sigma$
q (nm$^{-1}$)
C
Reweighted, Uniform
KDE($R_g$)
$R_g$ (Å)
$\langle R_{g\ uni} \rangle$ = 35.93 ± 0.07
$\langle R_{g\ rw} \rangle$ = 36.15 ± 0.12
KDE($D_{ee}$)
$D_{ee}$ (Å)
$\langle D_{ee\ uni} \rangle$ = 85.91 ± 0.32
$\langle D_{ee\ rw} \rangle$ = 88.32 ± 0.54
KDE($D_{max}$)
$D_{max}$ (Å)
$\langle D_{max\ uni} \rangle$ = 106.55 ± 0.23
$\langle D_{max\ rw} \rangle$ = 108.47 ± 0.40
D
aSyn Uniform
aSyn Reweighted
aSyn Difference
Residue
Å
$C_\alpha$ Distance
$C_\alpha$ Dist. Difference
E
aSyn Uniform
aSyn Reweighted
aSyn Difference
Frequency
Frequency Difference
Residue
H
E
C

**Figure S12. aSyn ensemble analysis and reweighting highlights.** Comparison of calculated structural properties for the uniformly weighted ensemble of aSyn (“uniform”) and the ensemble after reweighting using SAXS data via BME approach (“reweighted”). Differences are calculated by subtracting the uniformly weighted values from the reweighted ones. **A.** $\phi_{eff}$ versus $\chi^2$ plot across $\theta$ values , with the selected $\theta$ value highlighted in gold. **B.** SAXS profile fits from the uniform (black) and reweighted (gold) ensembles) against experimental data (top), and the respective standardized residuals (bottom). **C.** Probability density distributions of the radius of gyration ($R_g$), end-to-end distance ($D_{ee}$), and maximum $C_\alpha$-$C_\alpha$ distance ($D_{max}$) for both ensembles (black-uniform; gold-reweighted). Dashed lines indicate weighted average values for each distribution, with its values detailed on the right of the plots along with their respective standard error. **D.** $C_\alpha$-$C_\alpha$ distance matrices for the uniform and reweighted ensembles, and their difference. **E.** Secondary structure assignment frequency plots for the uniform and reweighted ensembles, and their difference.

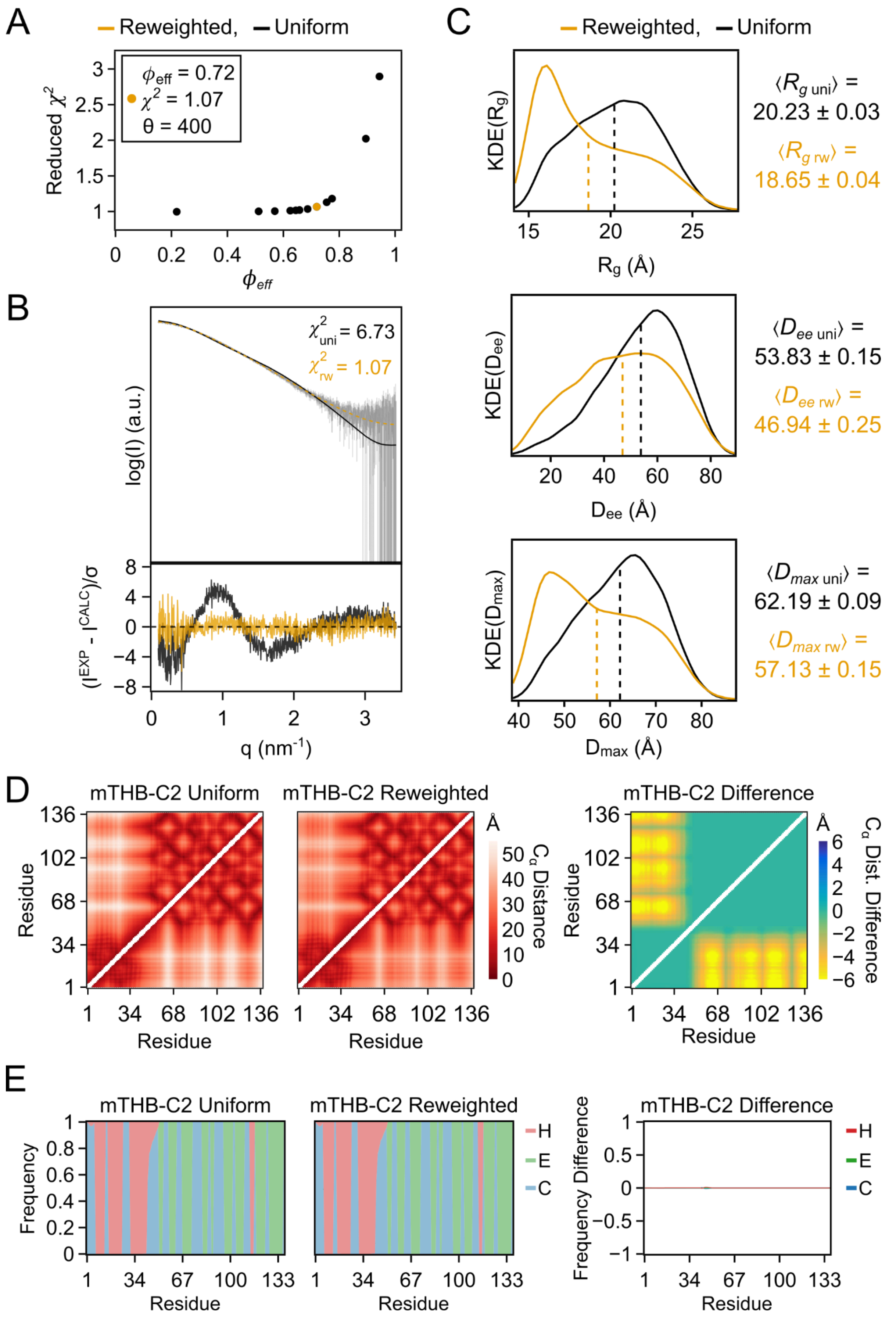

A
Reweighted, Uniform
$\phi_{eff}$ = 0.72
$\chi^2$ = 1.07
θ = 400
Reduced $\chi^2$
$\phi_{eff}$
B
$\chi^2_{uni}$ = 6.73
$\chi^2_{rw}$ = 1.07
log(I) (a.u.)
$(I^{EXP} - I^{CALC})/\sigma$
q (nm$^{-1}$)
C
Reweighted, Uniform
KDE(Rg)
Rg (Å)
⟨$R_{g\ uni}$⟩ = 20.23 ± 0.03
⟨$R_{g\ rw}$⟩ = 18.65 ± 0.04
KDE(Dee)
Dee (Å)
⟨$D_{ee\ uni}$⟩ = 53.83 ± 0.15
⟨$D_{ee\ rw}$⟩ = 46.94 ± 0.25
KDE(Dmax)
Dmax (Å)
⟨$D_{max\ uni}$⟩ = 62.19 ± 0.09
⟨$D_{max\ rw}$⟩ = 57.13 ± 0.15
D
mTHB-C2 Uniform
mTHB-C2 Reweighted
mTHB-C2 Difference
Residue
Cα Distance
Cα Dist. Difference
Å
E
mTHB-C2 Uniform
mTHB-C2 Reweighted
mTHB-C2 Difference
Frequency
Frequency Difference
Residue
H
E
C

**Figure S13. cMyBP-C$_{mTHBC2}$ ensemble analysis and reweighting highlights.** Comparison of calculated structural properties for the uniformly weighted ensemble of cMyBP-C$_{mTHBC2}$ ("uniform") and the ensemble after reweighting using SAXS data via BME approach ("reweighted"). Differences are calculated by subtracting the uniformly weighted values from the reweighted ones. **A.** $\phi_{eff}$ versus $\chi^2$ plot across $\theta$ values , with the selected $\theta$ value highlighted in gold. **B.** SAXS profile fits from the uniform (black) and reweighted (gold) ensembles) against experimental data (top), and the respective standardized residuals (bottom). **C.** Probability density distributions of the radius of gyration ($R_g$), end-to-end distance ($D_{ee}$), and maximum $C_\alpha$-$C_\alpha$ distance ($D_{max}$) for both ensembles (black-uniform; gold-reweighted). Dashed lines indicate weighted average values for each distribution, with its values detailed on the right of the plots along with their respective standard error. **D.** $C_\alpha$-$C_\alpha$ distance matrices for the uniform and reweighted ensembles, and their difference. **E.** Secondary structure assignment frequency plots for the uniform and reweighted ensembles, and their difference.

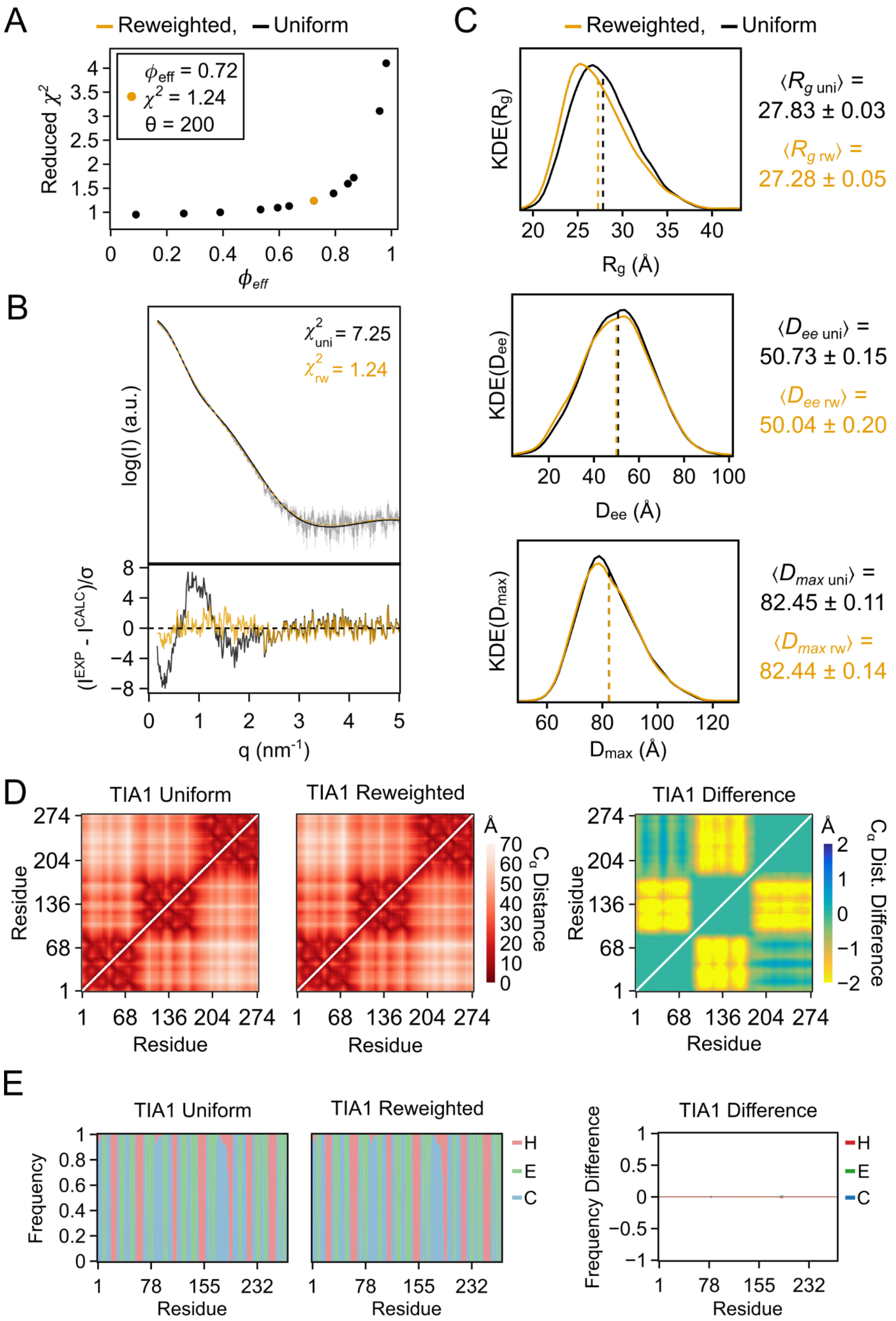
A
Reweighted, Uniform
ϕeff = 0.72
χ2 = 1.24
θ = 200
Reduced χ2
ϕeff
B
χ2uni = 7.25
χ2rw = 1.24
log(I) (a.u.)
(IEXP - ICALC)/σ
q (nm-1)
C
Reweighted, Uniform
KDE(Rg)
Rg (Å)
⟨Rg uni⟩ = 27.83 ± 0.03
⟨Rg rw⟩ = 27.28 ± 0.05
KDE(Dee)
Dee (Å)
⟨Dee uni⟩ = 50.73 ± 0.15
⟨Dee rw⟩ = 50.04 ± 0.20
KDE(Dmax)
Dmax (Å)
⟨Dmax uni⟩ = 82.45 ± 0.11
⟨Dmax rw⟩ = 82.44 ± 0.14
D
TIA1 Uniform
TIA1 Reweighted
TIA1 Difference
Residue
Cα Distance
Cα Dist. Difference
E
TIA1 Uniform
TIA1 Reweighted
TIA1 Difference
Frequency
Frequency Difference
Residue
H
E
C

**Figure S14. TIA1 ensemble analysis and reweighting highlights.** Comparison of calculated structural properties for the uniformly weighted ensemble of TIA1 (“uniform”) and the ensemble after reweighting using SAXS data via BME approach (“reweighted”). Differences are calculated by subtracting the uniformly weighted values from the reweighted ones. **A.** $\phi_{\mathrm{eff}}$ versus $\chi^2$ plot across $\theta$ values , with the selected $\theta$ value highlighted in gold. **B.** SAXS profile fits from the uniform (black) and reweighted (gold) ensembles) against experimental data (top), and the respective standardized residuals (bottom). **C.** Probability density distributions of the radius of gyration ($R_g$), end-to-end distance ($D_{ee}$), and maximum $C_\alpha$-$C_\alpha$ distance ($D_{max}$) for both ensembles (black-uniform; gold-reweighted). Dashed lines indicate weighted average values for each distribution, with its values detailed on the right of the plots along with their respective standard error. **D.** $C_\alpha$-$C_\alpha$ distance matrices for the uniform and reweighted ensembles, and their difference. **E.** Secondary structure assignment frequency plots for the uniform and reweighted ensembles, and their difference.

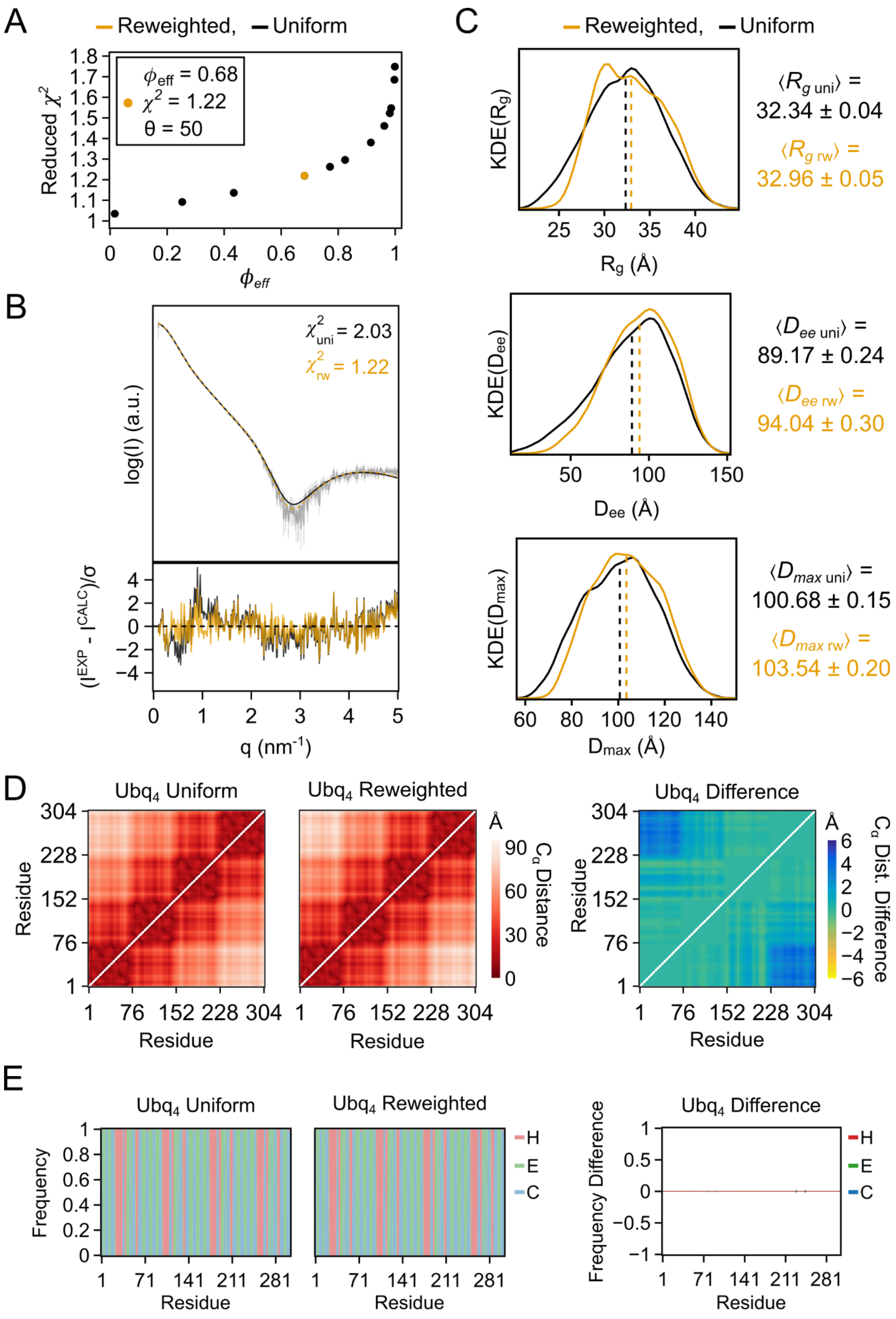

A
Reweighted, Uniform
$\phi_{eff}$ = 0.68
$\chi^2$ = 1.22
θ = 50
Reduced $\chi^2$
$\phi_{eff}$
B
$\chi^2_{uni}$ = 2.03
$\chi^2_{rw}$ = 1.22
log(I) (a.u.)
(I^EXP - I^CALC)/σ
q (nm^-1)
C
Reweighted, Uniform
KDE(Rg)
Rg (Å)
⟨Rg uni⟩ = 32.34 ± 0.04
⟨Rg rw⟩ = 32.96 ± 0.05
KDE(Dee)
Dee (Å)
⟨Dee uni⟩ = 89.17 ± 0.24
⟨Dee rw⟩ = 94.04 ± 0.30
KDE(Dmax)
Dmax (Å)
⟨Dmax uni⟩ = 100.68 ± 0.15
⟨Dmax rw⟩ = 103.54 ± 0.20
D
Ubq4 Uniform
Ubq4 Reweighted
Ubq4 Difference
Residue
Cα Distance
Cα Dist. Difference
Å
E
Ubq4 Uniform
Ubq4 Reweighted
Ubq4 Difference
Frequency
Frequency Difference
Residue
H
E
C

**Figure S15. $Ubq_4$ ensemble analysis and reweighting highlights.** Comparison of calculated structural properties for the uniformly weighted ensemble of $Ubq_4$ ("uniform") and the ensemble after reweighting using SAXS data via BME approach ("reweighted"). Differences are calculated by subtracting the uniformly weighted values from the reweighted ones. **A.** $\phi_{eff}$ versus $\chi^2$ plot across $\theta$ values , with the selected $\theta$ value highlighted in gold. **B.** SAXS profile fits from the uniform (black) and reweighted (gold) ensembles) against experimental data (top), and the respective standardized residuals (bottom). **C.** Probability density distributions of the radius of gyration ($R_g$), end-to-end distance ($D_{ee}$), and maximum $C_\alpha$-$C_\alpha$ distance ($D_{max}$) for both ensembles (black-uniform; gold-reweighted). Dashed lines indicate weighted average values for each distribution, with its values detailed on the right of the plots along with their respective standard error. **D.** $C_\alpha$-$C_\alpha$ distance matrices for the uniform and reweighted ensembles, and their difference. **E.** Secondary structure assignment frequency plots for the uniform and reweighted ensembles, and their difference.

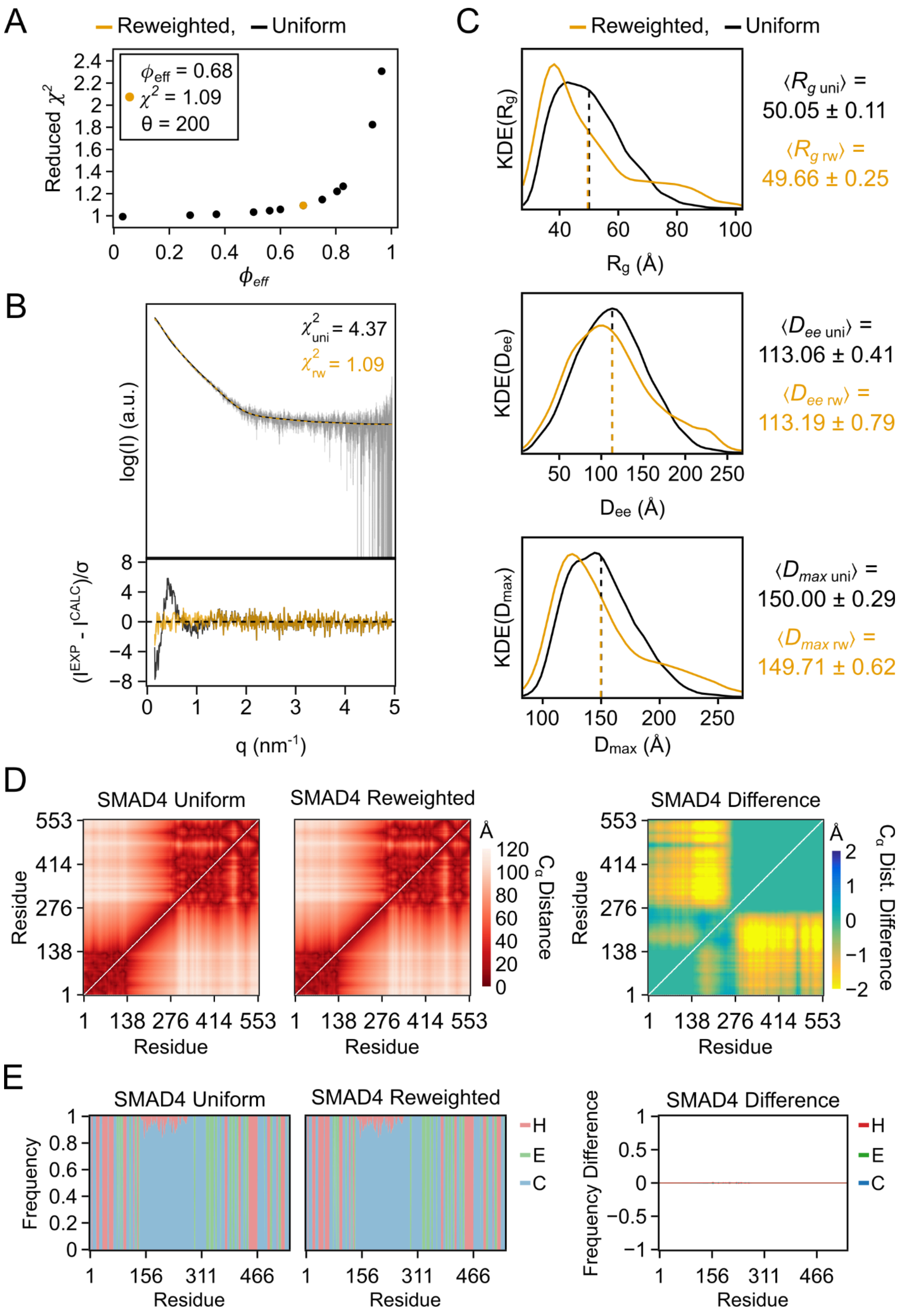

A
Reweighted, Uniform
$\phi_{eff}$ = 0.68
$\chi^2$ = 1.09
θ = 200
Reduced $\chi^2$
$\phi_{eff}$
B
$\chi^2_{uni}$ = 4.37
$\chi^2_{rw}$ = 1.09
log(I) (a.u.)
$(I^{EXP} - I^{CALC})/\sigma$
q (nm$^{-1}$)
C
Reweighted, Uniform
KDE(Rg)
Rg (Å)
$\langle R_{g\ uni} \rangle$ = 50.05 ± 0.11
$\langle R_{g\ rw} \rangle$ = 49.66 ± 0.25
KDE(Dee)
Dee (Å)
$\langle D_{ee\ uni} \rangle$ = 113.06 ± 0.41
$\langle D_{ee\ rw} \rangle$ = 113.19 ± 0.79
KDE(Dmax)
Dmax (Å)
$\langle D_{max\ uni} \rangle$ = 150.00 ± 0.29
$\langle D_{max\ rw} \rangle$ = 149.71 ± 0.62
D
SMAD4 Uniform
SMAD4 Reweighted
SMAD4 Difference
Residue
Cα Distance
Cα Dist. Difference
E
SMAD4 Uniform
SMAD4 Reweighted
SMAD4 Difference
Frequency
Frequency Difference
Residue
H
E
C

**Figure S16. SMAD4 ensemble analysis and reweighting highlights.** Comparison of calculated structural properties for the uniformly weighted ensemble of SMAD4 (“uniform”) and the ensemble after reweighting using SAXS data via BME approach (“reweighted”). Differences are calculated by subtracting the uniformly weighted values from the reweighted ones. **A.** $\phi_{eff}$ versus $\chi^2$ plot across $\theta$ values , with the selected $\theta$ value highlighted in gold. **B.** SAXS profile fits from the uniform (black) and reweighted (gold) ensembles) against experimental data (top), and the respective standardized residuals (bottom). **C.** Probability density distributions of the radius of gyration ($R_g$), end-to-end distance ($D_{ee}$), and maximum $C_\alpha$-$C_\alpha$ distance ($D_{max}$) for both ensembles (black-uniform; gold-reweighted). Dashed lines indicate weighted average values for each distribution, with its values detailed on the right of the plots along with their respective standard error. **D.** $C_\alpha$-$C_\alpha$ distance matrices for the uniform and reweighted ensembles, and their difference. **E.** Secondary structure assignment frequency plots for the uniform and reweighted ensembles, and their difference.

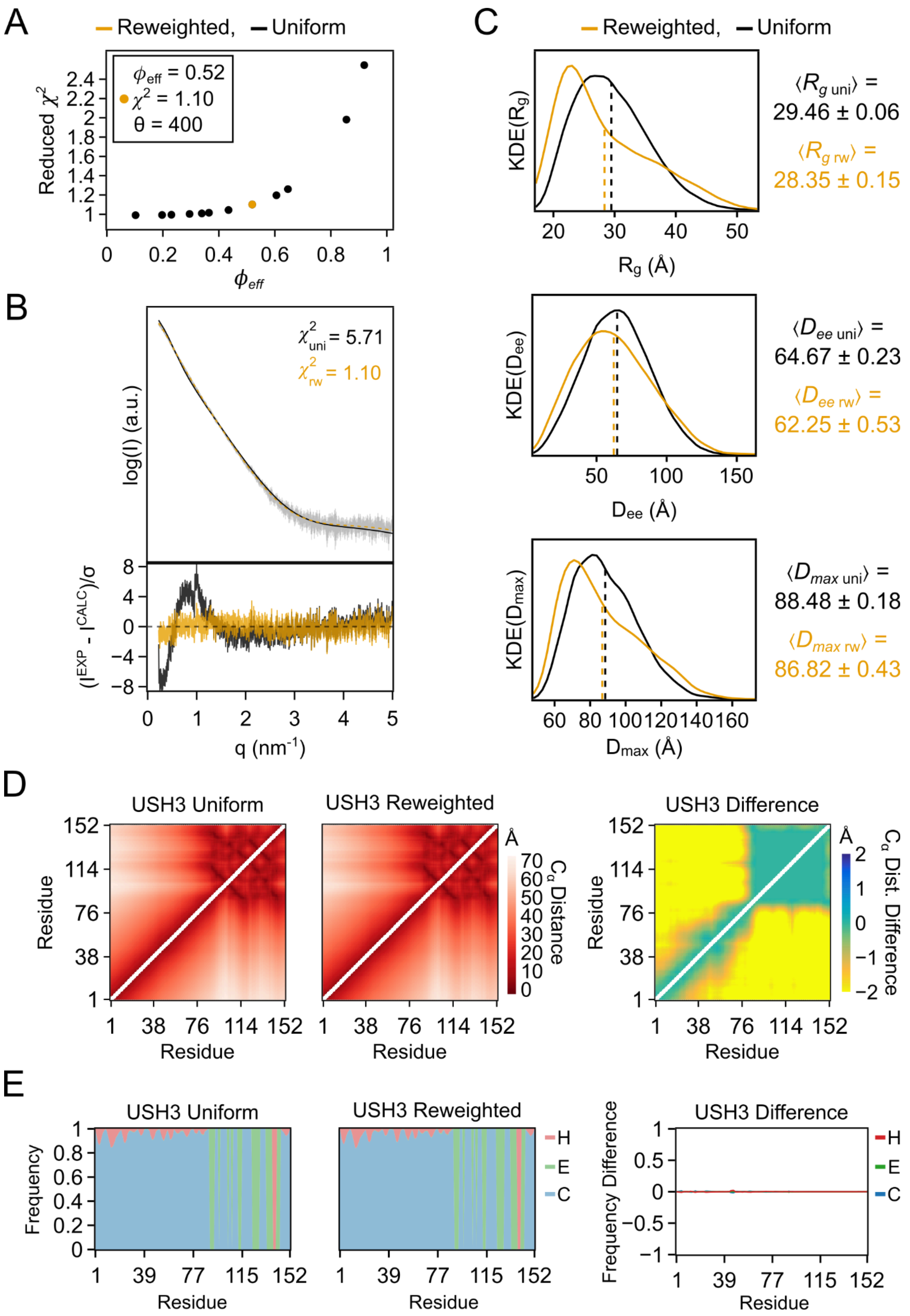

A
Reweighted, Uniform
$\phi_{eff}$ = 0.52
$\chi^2$ = 1.10
θ = 400
Reduced $\chi^2$
$\phi_{eff}$
B
$\chi^2_{uni}$ = 5.71
$\chi^2_{rw}$ = 1.10
log(I) (a.u.)
$(I^{EXP} - I^{CALC})/\sigma$
q (nm$^{-1}$)
C
Reweighted, Uniform
KDE(Rg)
Rg (Å)
⟨$R_{g\ uni}$⟩ = 29.46 ± 0.06
⟨$R_{g\ rw}$⟩ = 28.35 ± 0.15
KDE(Dee)
Dee (Å)
⟨$D_{ee\ uni}$⟩ = 64.67 ± 0.23
⟨$D_{ee\ rw}$⟩ = 62.25 ± 0.53
KDE(Dmax)
Dmax (Å)
⟨$D_{max\ uni}$⟩ = 88.48 ± 0.18
⟨$D_{max\ rw}$⟩ = 86.82 ± 0.43
D
USH3 Uniform
USH3 Reweighted
USH3 Difference
Residue
$C_\alpha$ Distance
$C_\alpha$ Dist. Difference
E
USH3 Uniform
USH3 Reweighted
USH3 Difference
Frequency
Frequency Difference
Residue
H
E
C

**Figure S17. USH3 ensemble analysis and reweighting highlights.** Comparison of calculated structural properties for the uniformly weighted ensemble of USH3 (“uniform”) and the ensemble after reweighting using SAXS data via BME approach (“reweighted”). Differences are calculated by subtracting the uniformly weighted values from the reweighted ones. **A.** $\phi_{\mathrm{eff}}$ versus $\chi^2$ plot across $\theta$ values , with the selected $\theta$ value highlighted in gold. **B.** SAXS profile fits from the uniform (black) and reweighted (gold) ensembles) against experimental data (top), and the respective standardized residuals (bottom). **C.** Probability density distributions of the radius of gyration ($R_g$), end-to-end distance ($D_{ee}$), and maximum $C_\alpha$-$C_\alpha$ distance ($D_{max}$) for both ensembles (black-uniform; gold-reweighted). Dashed lines indicate weighted average values for each distribution, with its values detailed on the right of the plots along with their respective standard error. **D.** $C_\alpha$-$C_\alpha$ distance matrices for the uniform and reweighted ensembles, and their difference. **E.** Secondary structure assignment frequency plots for the uniform and reweighted ensembles, and their difference.

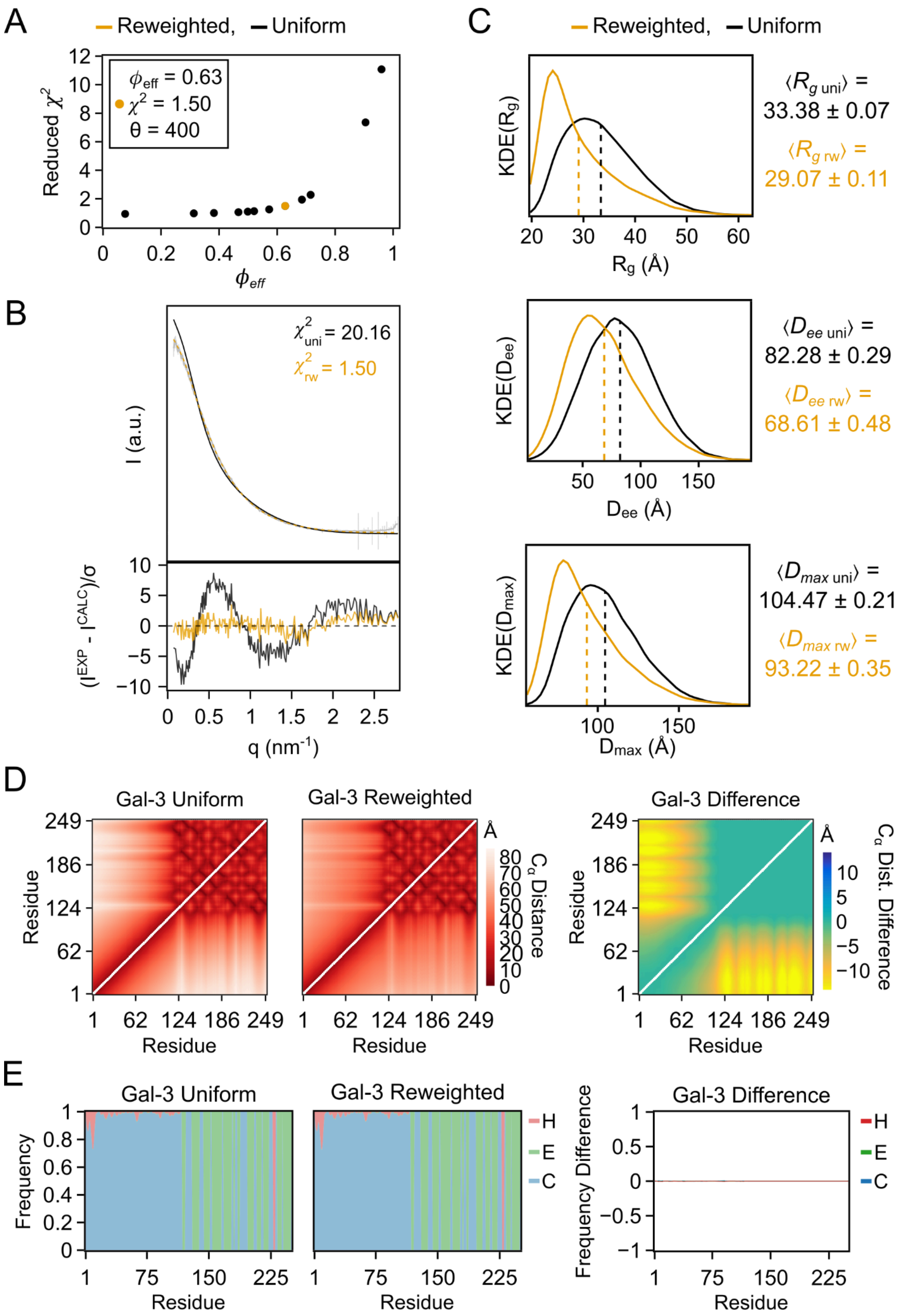
A
Reweighted, Uniform
$\phi_{eff}$ = 0.63
$\chi^2$ = 1.50
θ = 400
Reduced $\chi^2$
$\phi_{eff}$
B
$\chi^2_{uni}$ = 20.16
$\chi^2_{rw}$ = 1.50
I (a.u.)
$(I^{EXP} - I^{CALC})/\sigma$
q (nm$^{-1}$)
C
Reweighted, Uniform
KDE(Rg)
Rg (Å)
⟨$R_{g\ uni}$⟩ = 33.38 ± 0.07
⟨$R_{g\ rw}$⟩ = 29.07 ± 0.11
KDE(Dee)
Dee (Å)
⟨$D_{ee\ uni}$⟩ = 82.28 ± 0.29
⟨$D_{ee\ rw}$⟩ = 68.61 ± 0.48
KDE(Dmax)
Dmax (Å)
⟨$D_{max\ uni}$⟩ = 104.47 ± 0.21
⟨$D_{max\ rw}$⟩ = 93.22 ± 0.35
D
Gal-3 Uniform
Gal-3 Reweighted
Gal-3 Difference
Residue
Å
Cα Distance
Cα Dist. Difference
E
Gal-3 Uniform
Gal-3 Reweighted
Gal-3 Difference
Frequency
Frequency Difference
Residue
H
E
C

**Figure S18. Gal-3 ensemble analysis and reweighting highlights.** Comparison of calculated structural properties for the uniformly weighted ensemble of Gal-3 ("uniform") and the ensemble after reweighting using SAXS data via BME approach ("reweighted"). Differences are calculated by subtracting the uniformly weighted values from the reweighted ones. **A.** $\phi_{\mathrm{eff}}$ versus $\chi^2$ plot across $\theta$ values , with the selected $\theta$ value highlighted in gold. **B.** SAXS profile fits from the uniform (black) and reweighted (gold) ensembles) against experimental data (top), and the respective standardized residuals (bottom). **C.** Probability density distributions of the radius of gyration ($R_g$), end-to-end distance ($D_{ee}$), and maximum $C_\alpha$-$C_\alpha$ distance ($D_{max}$) for both ensembles (black-uniform; gold-reweighted). Dashed lines indicate weighted average values for each distribution, with its values detailed on the right of the plots along with their respective standard error. **D.** $C_\alpha$-$C_\alpha$ distance matrices for the uniform and reweighted ensembles, and their difference. **E.** Secondary structure assignment frequency plots for the uniform and reweighted ensembles, and their difference.

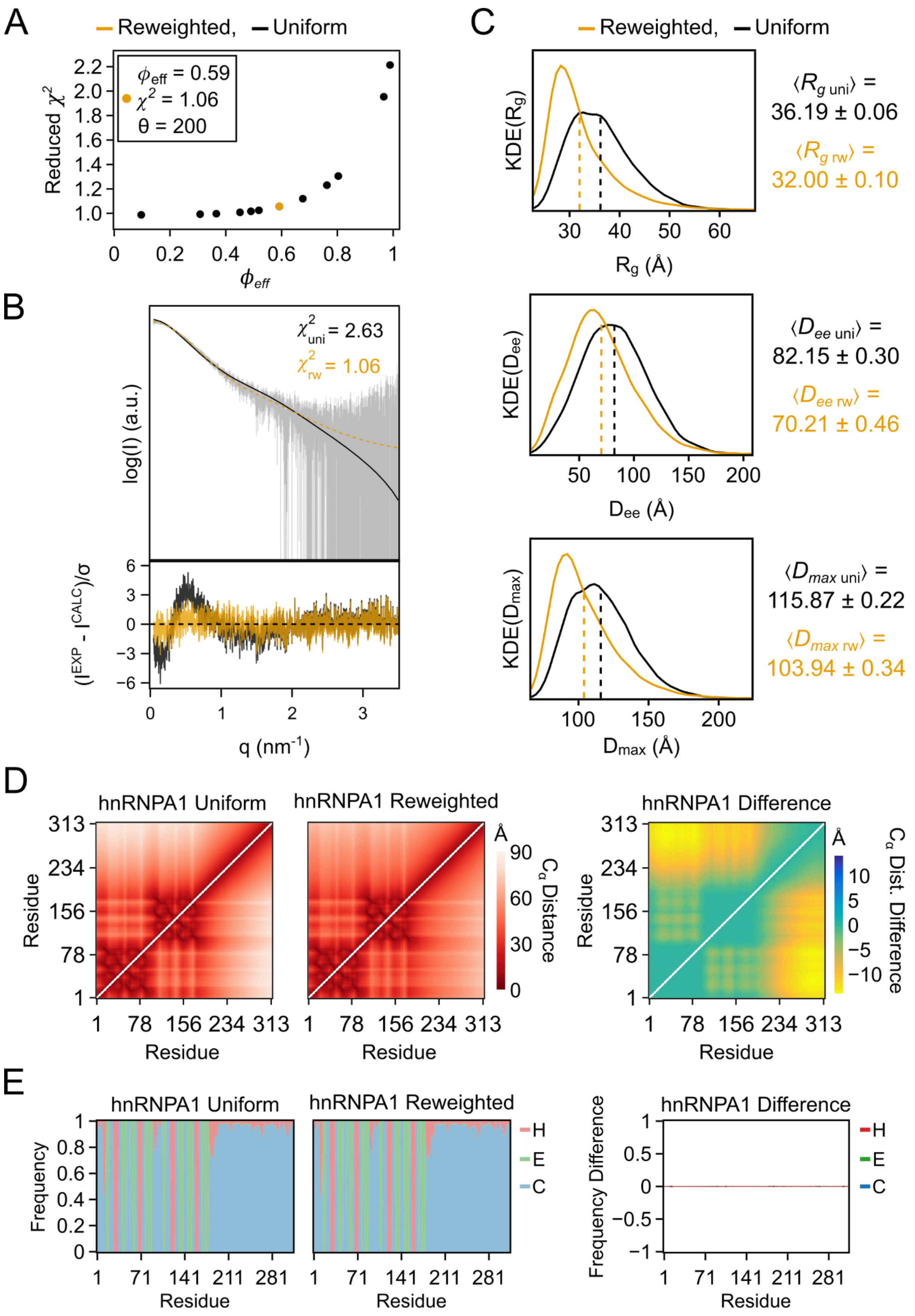

A
Reweighted, Uniform
Reduced $\chi^2$
$\phi_{eff}$ = 0.59
$\chi^2$ = 1.06
θ = 200
$\phi_{eff}$
B
log(I) (a.u.)
$\chi^2_{uni}$ = 2.63
$\chi^2_{rw}$ = 1.06
$(I^{EXP} - I^{CALC})/\sigma$
q (nm$^{-1}$)
C
KDE($R_g$)
$R_g$ (Å)
$\langle R_{g\ uni} \rangle$ = 36.19 ± 0.06
$\langle R_{g\ rw} \rangle$ = 32.00 ± 0.10
KDE($D_{ee}$)
$D_{ee}$ (Å)
$\langle D_{ee\ uni} \rangle$ = 82.15 ± 0.30
$\langle D_{ee\ rw} \rangle$ = 70.21 ± 0.46
KDE($D_{max}$)
$D_{max}$ (Å)
$\langle D_{max\ uni} \rangle$ = 115.87 ± 0.22
$\langle D_{max\ rw} \rangle$ = 103.94 ± 0.34
D
hnRNPA1 Uniform
hnRNPA1 Reweighted
hnRNPA1 Difference
Residue
$C_\alpha$ Distance
$C_\alpha$ Dist. Difference
E
hnRNPA1 Uniform
hnRNPA1 Reweighted
hnRNPA1 Difference
Frequency
Frequency Difference
H
E
C
Residue

**Figure S19. hnRNPA1 ensemble analysis and reweighting highlights.** Comparison of calculated structural properties for the uniformly weighted ensemble of hnRNPA1 (“uniform”) and the ensemble after reweighting using SAXS data via BME approach (“reweighted”). Differences are calculated by subtracting the uniformly weighted values from the reweighted ones. **A.** $\phi_{\mathrm{eff}}$ versus $\chi^2$ plot across $\theta$ values , with the selected $\theta$ value highlighted in gold. **B.** SAXS profile fits from the uniform (black) and reweighted (gold) ensembles) against experimental data (top), and the respective standardized residuals (bottom). **C.** Probability density distributions of the radius of gyration ($R_g$), end-to-end distance ($D_{ee}$), and maximum $C_\alpha$-$C_\alpha$ distance ($D_{max}$) for both ensembles (black-uniform; gold-reweighted). Dashed lines indicate weighted average values for each distribution, with its values detailed on the right of the plots along with their respective standard error. **D.** $C_\alpha$-$C_\alpha$ distance matrices for the uniform and reweighted ensembles, and their difference. **E.** Secondary structure assignment frequency plots for the uniform and reweighted ensembles, and their difference.

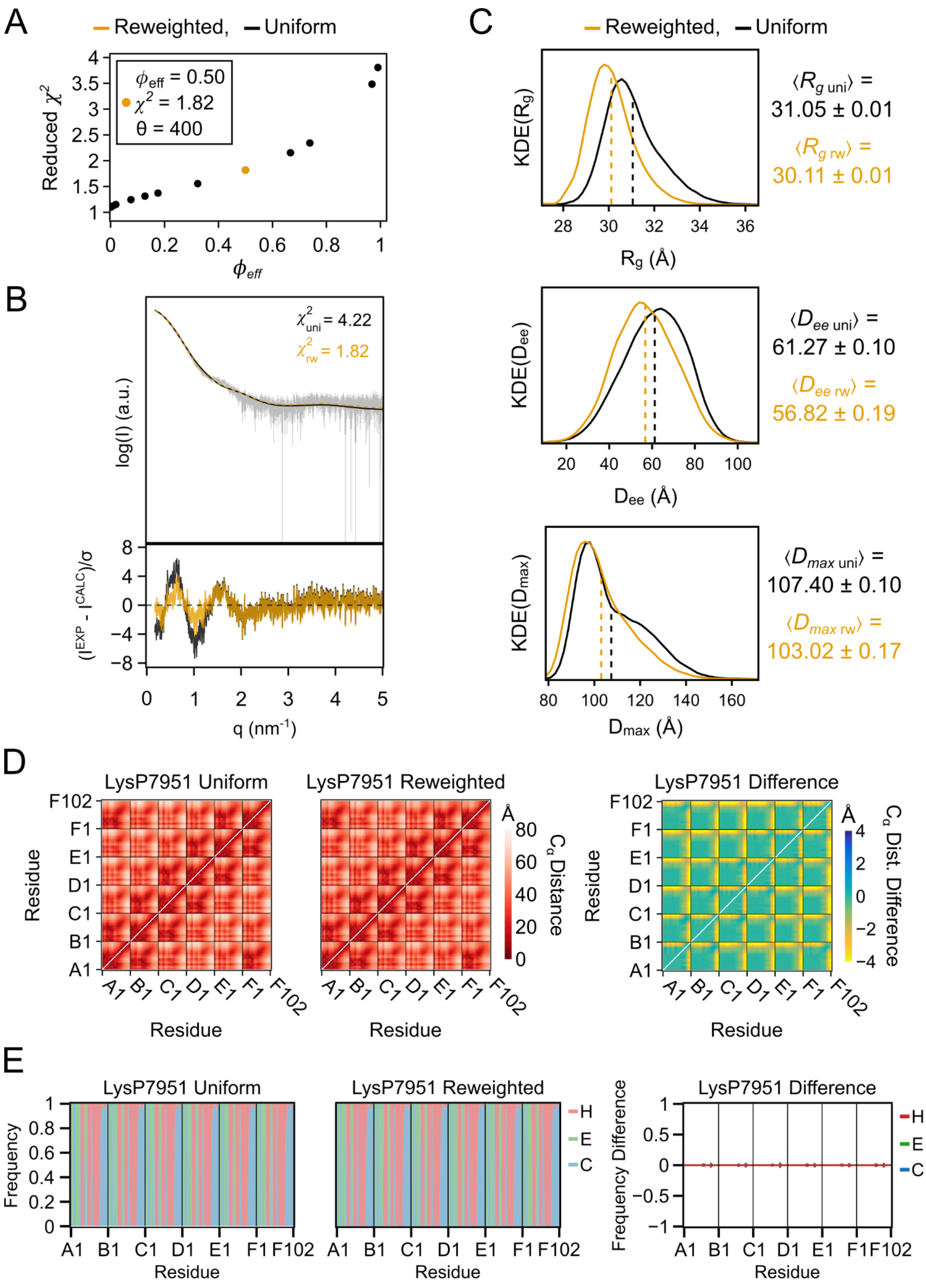

A
Reweighted, Uniform
Reduced χ²
ϕeff = 0.50
χ² = 1.82
θ = 400
ϕeff
B
χ²uni = 4.22
χ²rw = 1.82
log(I) (a.u.)
(I^EXP - I^CALC)/σ
q (nm⁻¹)
C
Reweighted, Uniform
KDE(Rg)
Rg (Å)
⟨Rg uni⟩ = 31.05 ± 0.01
⟨Rg rw⟩ = 30.11 ± 0.01
KDE(Dee)
Dee (Å)
⟨Dee uni⟩ = 61.27 ± 0.10
⟨Dee rw⟩ = 56.82 ± 0.19
KDE(Dmax)
Dmax (Å)
⟨Dmax uni⟩ = 107.40 ± 0.10
⟨Dmax rw⟩ = 103.02 ± 0.17
D
LysP7951 Uniform
LysP7951 Reweighted
LysP7951 Difference
Residue
Cα Distance
Cα Dist. Difference
E
LysP7951 Uniform
LysP7951 Reweighted
LysP7951 Difference
Frequency
Frequency Difference
H
E
C
Residue

**Figure S20. LysP7951 ensemble analysis and reweighting highlights.** Comparison of calculated structural properties for the uniformly weighted ensemble of LysP7951 ("uniform") and the ensemble after reweighting using SAXS data via BME approach ("reweighted"). Differences are calculated by subtracting the uniformly weighted values from the reweighted ones. **A.** $\phi_{\mathrm{eff}}$ versus $\chi^2$ plot across $\theta$ values , with the selected $\theta$ value highlighted in gold. **B.** SAXS profile fits from the uniform (black) and reweighted (gold) ensembles) against experimental data (top), and the respective standardized residuals (bottom). **C.** Probability density distributions of the radius of gyration ($R_g$), end-to-end distance ($D_{ee}$), and maximum $C_\alpha$-$C_\alpha$ distance ($D_{max}$) for both ensembles (black-uniform; gold-reweighted). Dashed lines indicate weighted average values for each distribution, with its values detailed on the right of the plots along with their respective standard error. **D.** $C_\alpha$-$C_\alpha$ distance matrices for the uniform and reweighted ensembles, and their difference. **E.** Secondary structure assignment frequency plots for the uniform and reweighted ensembles, and their difference.

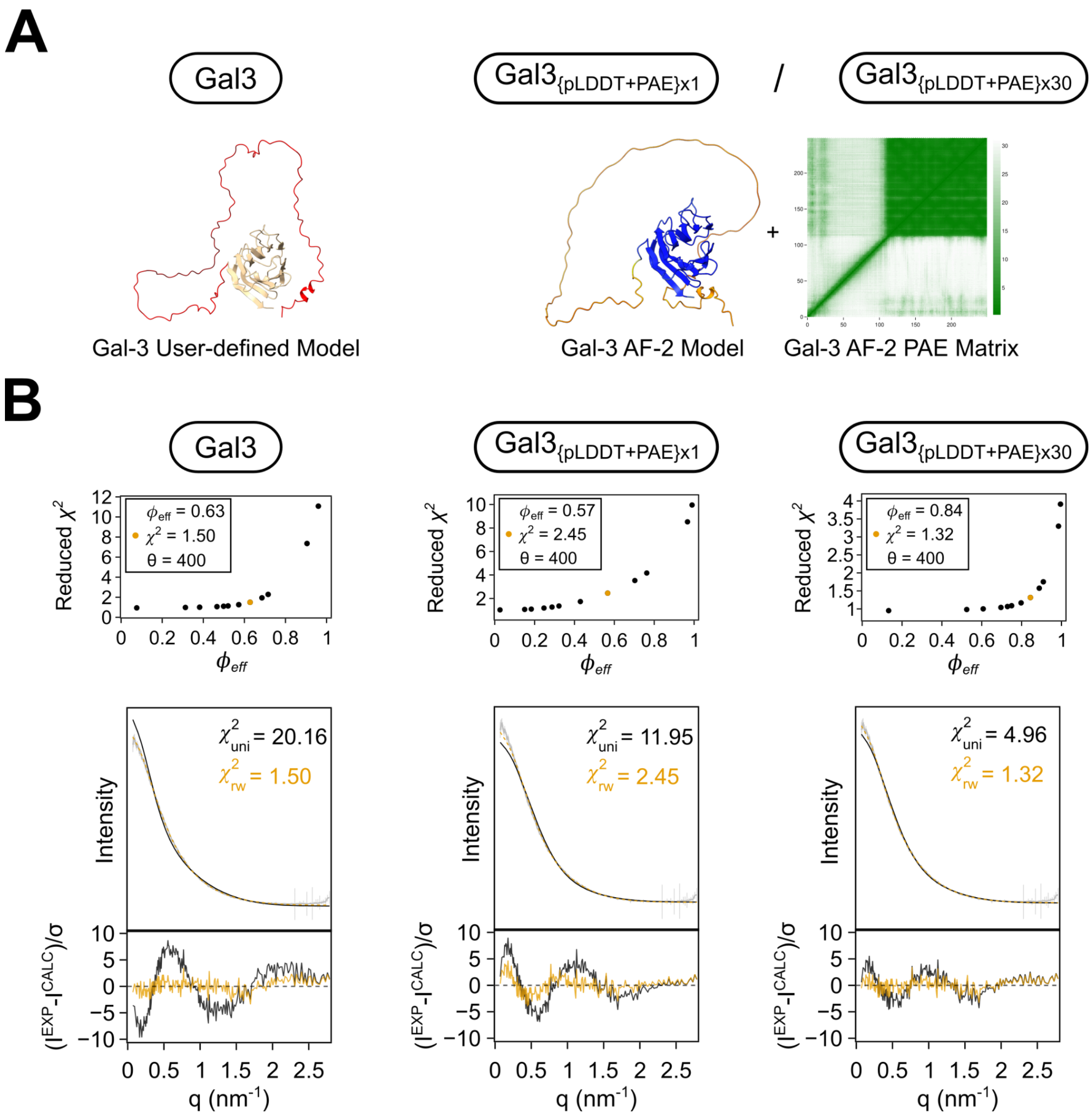

**Figure S21. AlphaFold-guided sampling can improve ensemble accuracy. A.** Inputs for generating Gal-3 ensembles: a user-defined model was used to generate an ensemble with custom sampling regions (Gal3). The AF-2 prediction was used to generate two ensembles, with sampling regions chosen based on low pLDDT values and the addition of inter-residue energy restraints based on PAE values. These differed in the value for the scaling factor applied to the PAE restraints: either no scaling was applied, with a value of 1 ($Gal3_{\{pLDDT+PAE\}x1}$), or a scaling factor of 30 was applied ($Gal3_{\{pLDDT+PAE\}x30}$). **B.** Fittings of Gal-3 ensembles to experimental SAXS data, before and after BME reweighting. For each ensemble, the $\phi_{eff}$ versus $\chi^2$ plot is shown, along with the fitting of theoretical SAXS data calculated from the ensemble (black) to experimental SAXS data of Gal-3 (grey). After applying BME reweighting with a value of 400 for the $\theta$ parameter, a new theoretical SAXS curve was calculated (gold) and re-fitted to experimental data. The reduced $\chi^2$ values of fitting each curve to the experimental data are shown, with the standardized residuals of each fitting presented below the plot.

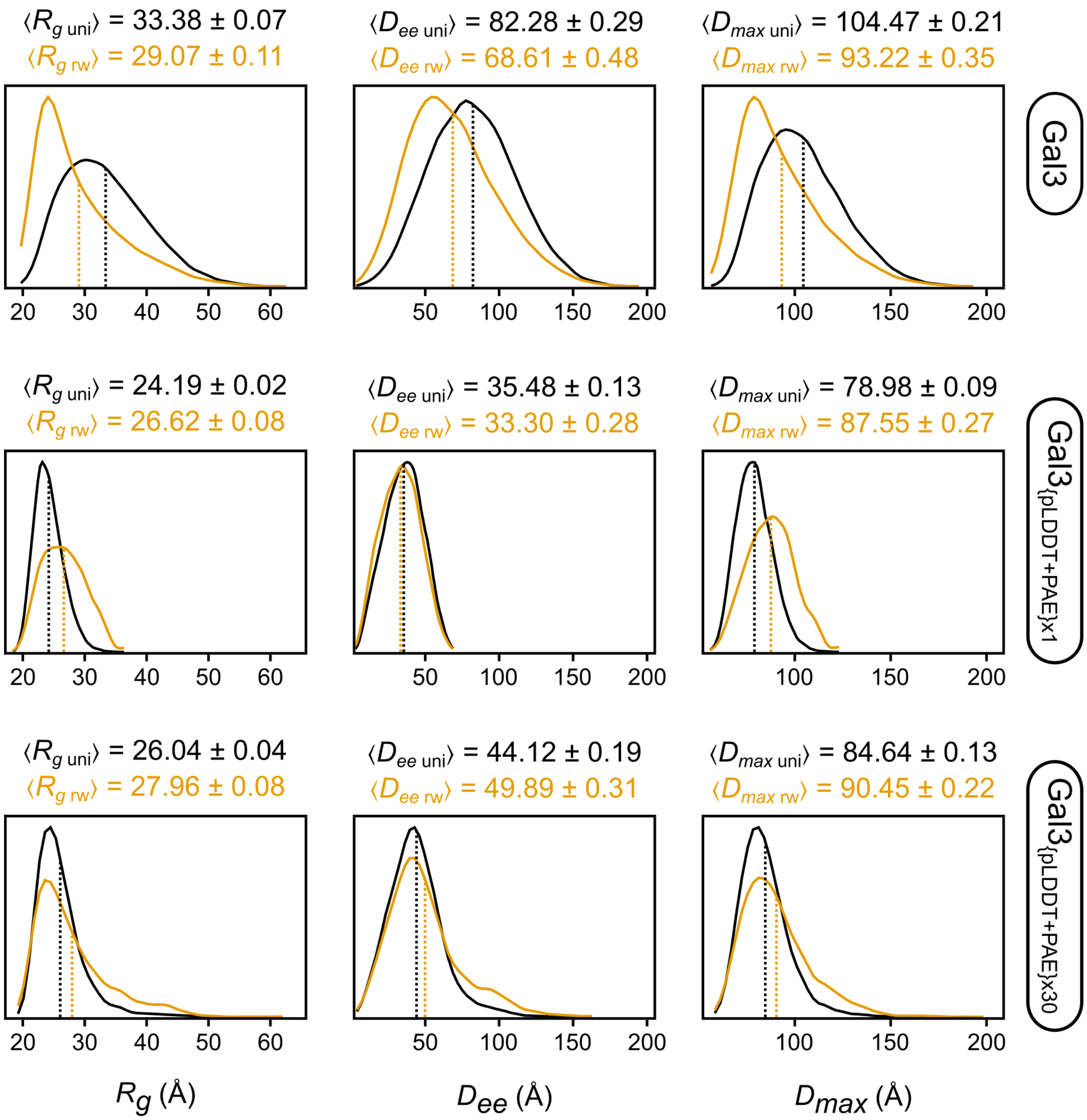

**Figure S22. Structural metrics distributions for the $R_g$, $D_{ee}$ and $D_{max}$ for generated ensembles of Gal-3, before and after BME reweighting.** A value of 400 for the $\theta$ parameter was used for reweighting all ensembles, and new structural metrics distributions were calculated using the optimized weights. Above each plot are the average values for the distributions of ensembles with uniform weights (black) and reweighted ensembles (gold).

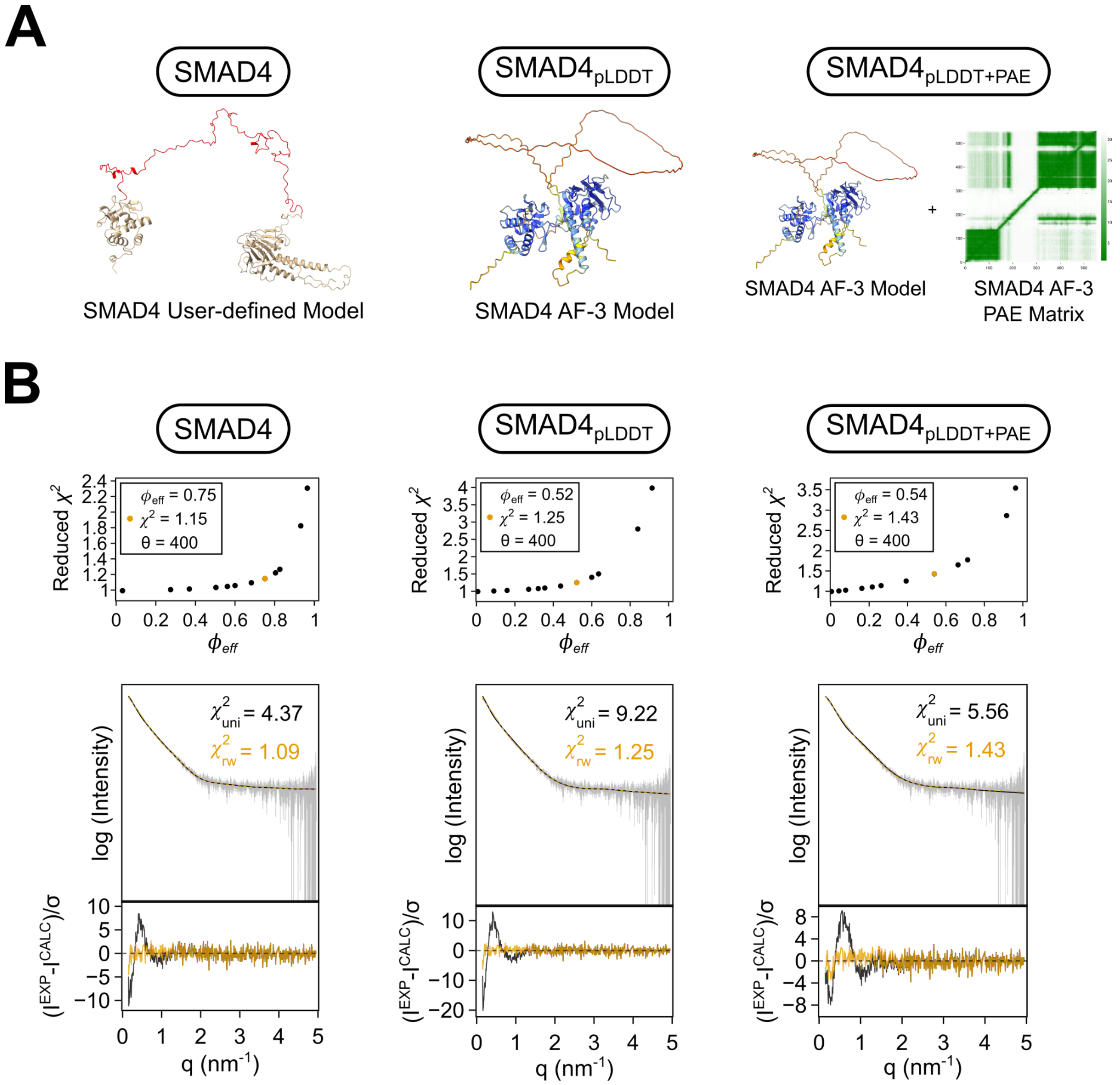

**Figure S23. AlphaFold-guided sampling can enable the study of specific structural states. A.** Inputs for generating SMAD4 ensembles: a user-defined model was used to generate an ensemble with custom sampling regions (SMAD4). The AF-3 prediction was used to generate two ensembles, one only choosing sampling regions based on low pLDDT values ($SMAD4_{pLDDT}$) and another using the pLDDT to choose sampling regions and the PAE matrix for the creation of inter-residue energy restraints based on PAE values ($SMAD4_{pLDDT+PAE}$). **B.** Fittings of SMAD4 ensembles to experimental SAXS data, before and after BME reweighting. For each ensemble, the $\phi_{eff}$ versus $\chi^2$ plot is shown, along with the fitting of theoretical SAXS data calculated from the ensemble (black) to experimental SAXS data of SMAD4 (grey). After applying BME reweighting with a value of 400 for the $\theta$ parameter, a new theoretical SAXS curve was calculated (gold) and re-fitted to experimental data. The reduced $\chi^2$ values of fitting each curve to the experimental data are shown, with the standardized residuals of each fitting presented below the plot.

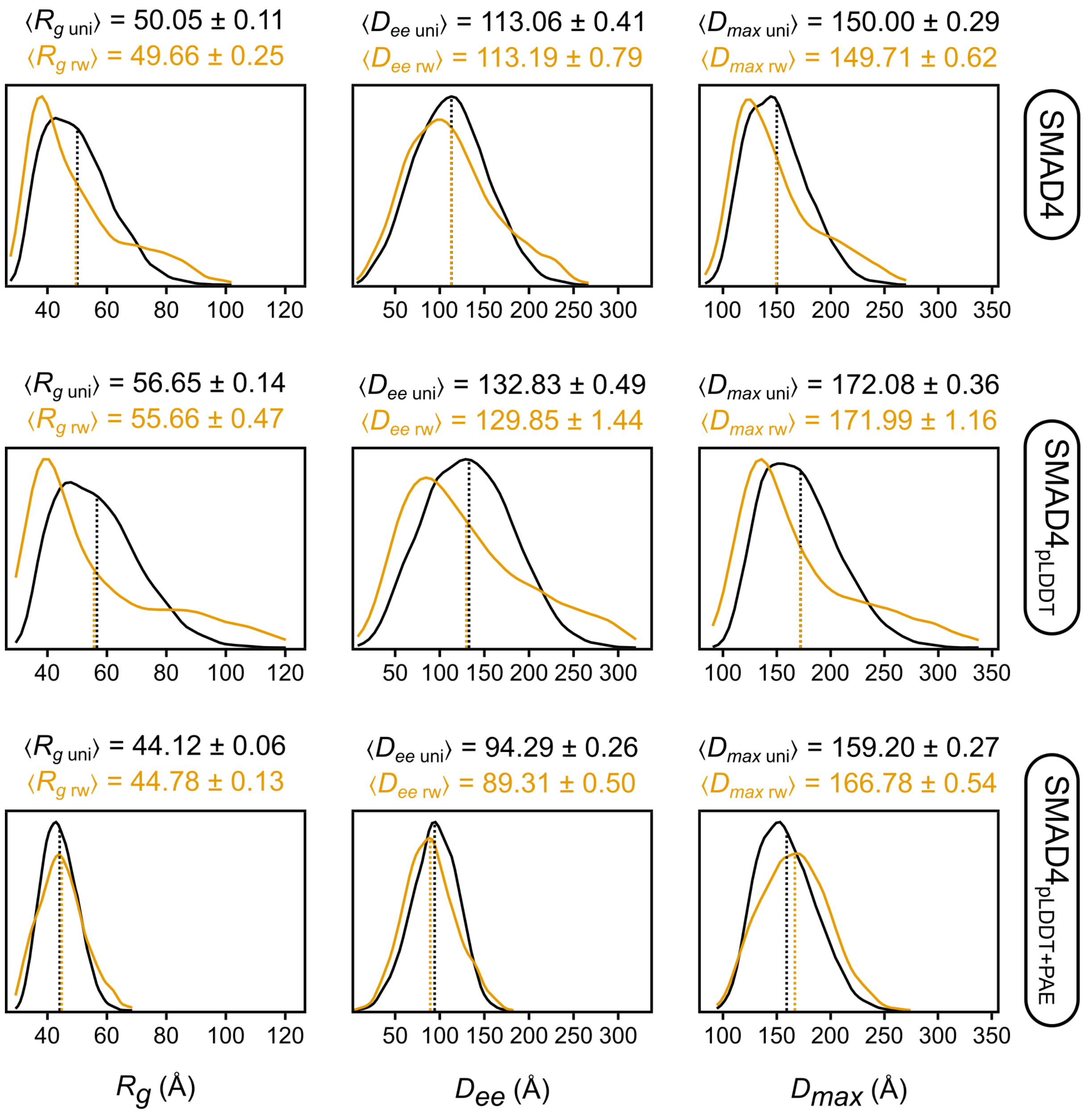

**Figure S24. Structural metrics distributions for the $R_g$, $D_{ee}$ and $D_{max}$ for generated ensembles of SMAD4, before and after BME reweighting.** A value of 400 for the $\theta$ parameter was used for reweighting all ensembles, and new structural metrics distributions were calculated using the optimized weights. Above each plot are the average values for the distributions of ensembles with uniform weights (black) and reweighted ensembles (gold).

Jung JE, Ewing MA, Valentine SJ *et al.* Structural Insights into Linkage-Specific Ubiquitin Chains Using Ion Mobility Mass Spectrometry. *J Am Soc Mass Spectrom*

2024;**35**:982–91.

Jussupow A, Messias AC, Stehle R *et al.* The dynamics of linear polyubiquitin. *Sci Adv* 2020;**6**, DOI: 10.1126/sciadv.abc3786.

Larsen AH, Wang Y, Bottaro S *et al.* Combining molecular dynamics simulations with small-angle X-ray and neutron scattering data to study multi-domain proteins in solution. *PLoS Comput Biol* 2020;**16**:e1007870.

Leaver-Fay A, Tyka M, Lewis SM *et al.* ROSETTA3: an object-oriented software suite for the simulation and design of macromolecules. *Meth Enzymol* 2011;**487**:545–74.

Lepur A, Salomonsson E, Nilsson UJ *et al.* Ligand induced galectin-3 protein self-association. *J Biol Chem* 2012;**287**:21751–6.

Levengood JD, Potoyan D, Penumutchu S *et al.* Thermodynamic coupling of the tandem RRM domains of hnRNP A1 underlie its pleiotropic RNA binding functions. *Sci Adv* 2024;**10**:eadk6580.

Lin Y-H, Qiu D-C, Chang W-H *et al.* The intrinsically disordered N-terminal domain of galectin-3 dynamically mediates multisite self-association of the protein through fuzzy interactions. *J Biol Chem* 2017;**292**:17845–56.

Martin EW, Thomasen FE, Milkovic NM *et al.* Interplay of folded domains and the disordered low-complexity domain in mediating hnRNPA1 phase separation. *Nucleic Acids Res* 2021;**49**:2931–45.

McKinney W. Data structures for statistical computing in python. *Proceedings of the 9th Python in Science Conference*. SciPy, 2010, 56–61.

Melino S, Rufini S, Sette M *et al.* Zn(2+) ions selectively induce antimicrobial salivary peptide histatin-5 to fuse negatively charged vesicles. Identification and characterization of a zinc-binding motif present in the functional domain. *Biochemistry* 1999;**38**:9626–33.

Metropolis N, Rosenbluth AW, Rosenbluth MN *et al.* Equation of state calculations by fast computing machines. *J Chem Phys* 1953;**21**:1087.

Michie KA, Kwan AH, Tung C-S *et al.* A Highly Conserved Yet Flexible Linker Is Part of a Polymorphic Protein-Binding Domain in Myosin-Binding Protein C. *Structure* 2016;**24**:2000–7.

Oppenheim FG, Xu T, McMillian FM *et al.* Histatins, a novel family of histidine-rich proteins in human parotid secretion. Isolation, characterization, primary structure, and fungistatic effects on Candida albicans. *J Biol Chem* 1988;**263**:7472–7.

Pally D, Bhat R. N-terminal tail prolines of Gal-3 mediate its oligomerization/phase separation. *Proc Natl Acad Sci USA* 2021;**118**, DOI: 10.1073/pnas.2107023118.

Pan Z, Mu J, Chen H-F. Balanced Three-Point Water Model OPC3-B for Intrinsically Disordered and Ordered Proteins. *J Chem Theory Comput* 2023;**19**:4837–50.

Piana S, Donchev AG, Robustelli P *et al.* Water dispersion interactions strongly influence simulated structural properties of disordered protein states. *J Phys Chem B* 2015;**119**:5113–23.

Pinto D, Gonçalo R, Louro M *et al.* On the Occurrence and Multimerization of Two-Polypeptide Phage Endolysins Encoded in Single Genes. *Microbiol Spectr* 2022;**10**:e0103722.

Qin B, Lam SS, Lin K. Crystal structure of a transcriptionally active Smad4 fragment. *Structure* 1999;**7**:1493–503.

Raj PA, Marcus E, Sukumaran DK. Structure of human salivary histatin 5 in aqueous and nonaqueous solutions. *Biopolymers* 1998.

Ramis R, Ortega-Castro J, Casasnovas R *et al.* A Coarse-Grained Molecular Dynamics Approach to the Study of the Intrinsically Disordered Protein α-Synuclein. *J Chem Inf Model* 2019;**59**:1458–71.

Ritsch I, Lehmann E, Emmanouilidis L *et al.* Phase Separation of Heterogeneous Nuclear Ribonucleoprotein A1 upon Specific RNA-Binding Observed by Magnetic Resonance. *Angew Chem Int Ed* 2022;**61**:e202204311.

Rohl CA, Strauss CEM, Misura KMS *et al.* Protein structure prediction using Rosetta. *Meth Enzymol* 2004;**383**:66–93.

Sato S. Why does galectin-3 have a unique intrinsically disordered region? “Raison d’être” for the disordered structure and liquid–liquid phase separation –Part 1–. *Glycoforum* 2023a.

Sato S. Why does galectin-3 have a unique intrinsically disordered region? “Raison d’être” for the disordered structure and liquid–liquid phase separation –Part 2–. *Glycoforum* 2023b.

Schwalbe M, Ozenne V, Bibow S *et al.* Predictive atomic resolution descriptions of intrinsically disordered hTau40 and α-synuclein in solution from NMR and small angle scattering. *Structure* 2014;**22**:238–49.

Shah M, Noronha V, Shetty O *et al.* SMAD4: A case-based review of the literature and current treatment options. *Cancer Research, Statistics, and Treatment* 2023;**6**.

Shrestha UR, Juneja P, Zhang Q *et al.* Generation of the configurational ensemble of an intrinsically disordered protein from unbiased molecular dynamics simulation. *Proc Natl Acad Sci USA* 2019;**116**:20446–52.

Shrestha UR, Smith JC, Petridis L. Full structural ensembles of intrinsically disordered proteins from unbiased molecular dynamics simulations. *Commun Biol* 2021;**4**:243.

Sonntag M, Jagtap PKA, Simon B *et al.* Segmental, Domain-Selective Perdeuteration and Small-Angle Neutron Scattering for Structural Analysis of Multi-Domain Proteins. *Angew Chem Int Ed* 2017;**56**:9322–5.

Strelkova MA, Tolstova AP, Mitkevich VA *et al.* Structure of Full-Length Src Kinase and Its Key Phosphorylated States: Molecular Dynamics Study. *Int J Mol Sci* 2024;**25**, DOI: 10.3390/ijms252212391.

Thomasen FE, Skaalum T, Kumar A *et al.* Rescaling protein-protein interactions improves Martini 3 for flexible proteins in solution. *Nat Commun* 2024;**15**:6645.

Tofaris GK. Initiation and progression of α-synuclein pathology in Parkinson’s disease. *Cell Mol Life Sci* 2022;**79**:210.

Volles MJ, Lansbury PT. Zeroing in on the pathogenic form of alpha-synuclein and its mechanism of neurotoxicity in Parkinson’s disease. *Biochemistry* 2003;**42**:7871–8.

Wang C, Bradley P, Baker D. Protein-protein docking with backbone flexibility. *J Mol Biol* 2007;**373**:503–19.

Wang I, Hennig J, Jagtap PKA *et al.* Structure, dynamics and RNA binding of the multi-domain splicing factor TIA-1. *Nucleic Acids Res* 2014;**42**:5949–66.
